# RCX – an R package adapting the Cytoscape Exchange format for biological networks

**DOI:** 10.1101/2021.10.26.466001

**Authors:** Florian Auer, Frank Kramer

## Abstract

**Motivation:** The Cytoscape Exchange (CX) format is a JSON-based data structure designed for the transmission of biological networks using standard web technologies. It was developed by the network data exchange (NDEx), which itself serves as online commons to share and collaborate on biological networks. Furthermore, the Cytoscape software for the analysis and visualization of biological networks contributes structure elements to capture the visual layout within the CX format. However, there is a fundamental difference between data handling in web standards and R. A manual conversion requires detailed knowledge of the CX format to reproduce and work with the networks.

**Results:** Here we present a software package to create, handle, validate, visualize and convert networks in CX format to standard data types and objects within R. Networks in this format can serve as a source for biological knowledge, and also capture the results of the analysis of those while preserving the visual layout across all platforms. The RCX package connects the R environment for statistical computing with outside platforms for storage and collaboration, as well as further analysis and visualization of biological networks.

**Availability:** RCX is a free and open-source R package, available on Bioconductor from release 3.15 (https://bioconductor.org/packages/RCX) and via GitHub (https://github.com/frankkramer-lab/RCX).

**Supplementary information:** Supplementary data are available at *Bioinformatics Advances* online.

## 1 Introduction

Biological networks are a common and widely used resource to capture associations between any types of biological entities such as genes, transcripts, proteins, metabolites, ligands, diseases, or drugs. Furthermore, the data formats used for encoding the network information differ heavily depending on the contained data and their intended use.

A variety of public databases provide their biological knowledge in domain-specific exchange formats. In subsequent analyses, those networks are further enriched with heterogeneous data and therefore require a more flexible format for capturing their content. Additionally, the layout and visualization of networks are often not considered as part of the network and omitted.

The Cytoscape exchange (CX) format covers the above shortcomings by following an aspect-based design: The network is split into independent modules (aspects) with specific schemes for the information they contain. For example, the *edges* aspect comprises the interactions between the nodes defined in the *nodes* aspect, and the *cartesianLayout* aspect provides the coordinates to position the nodes in space. The CX format was developed by the Network Data Exchange (NDEx), an online commons for biological networks (Pratt et al. 2015). The schemes for aspects responsible for storing visual attributes are derived from Cytoscape (Shannon et al. 2003), one of the most popular open-source software tools for the analysis and visualization of biomedical networks. Both Cytoscape and NDEx use the CX format for the exchange of the networks between their platforms with consistent visualizations.

Users of the statistical programming language R (R Development Core Team 2008) can use existing packages like rBiopaxParser (Kramer et al. 2013) to retrieve biological knowledge from public databases and conduct further analyses. The ndexr package (Auer et al. 2018) interfaces with the NDEx platform to store subsequent results. However, the included data model is a simple conversion of the JSON structure that conflicts with the table-oriented approach in R. Consequently, constructing valid networks requires advance knowledge and hinders the adoption of the software by a large user base. The RCX package implements the aspect-oriented design while allowing the user to focus on the networks instead of the underlying data structure.

## 2 Features

The RCX package provides custom functions for the creation and modification of aspects and networks in RCX format. Additional functions are provided to validate data types, aspects properties, and references between the aspects, even after manual editing. The CX format also requires a meta-data aspect that provides an overview of the included aspects (e.g., number of elements within the aspects). Within RCX objects, this meta-data is created and updated automatically.

The RCX package not only provides accessibility of networks in CX format, but it also provides conversion to and from objects of iGraph (Csardi and Nepusz 2006) and Bioconductor graph (Gentleman et al. 2021), both widely used libraries for graph manipulation and network analysis (Fig. 1).

**Fig. 1.**
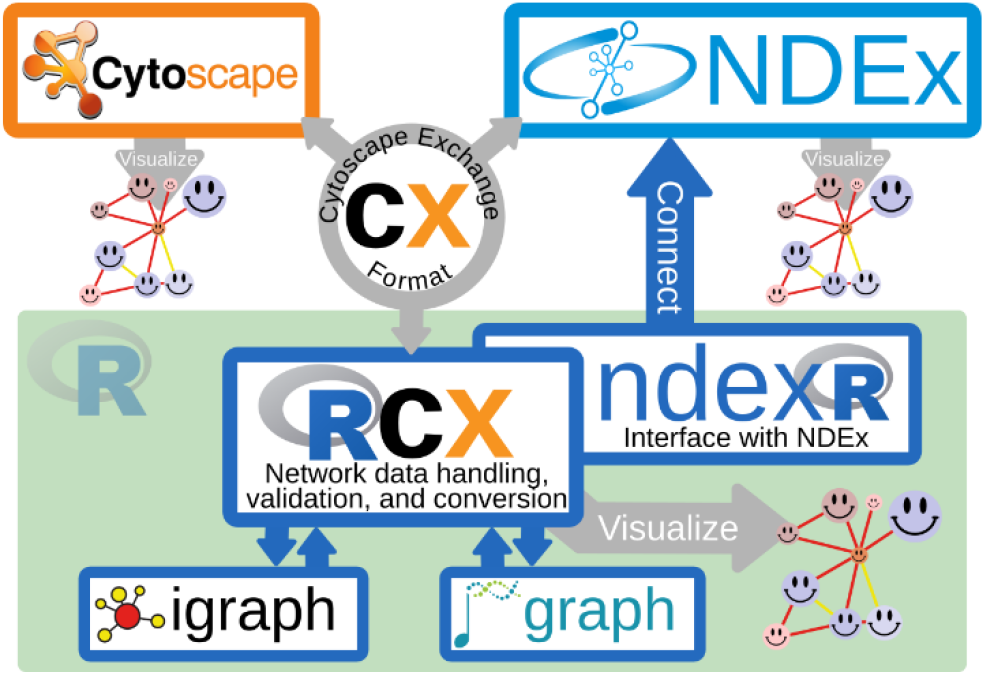
The RCX package connects the CX transmission format with established analysis libraries in R. The CX format is shared between RCX, Cytoscape and NDEx, while the consistent visual representation is preserved.

The R-based visualization of the networks is congruent with its representation in both the NDEx platform and Cytoscape. It also can be exported as an HTML file for further use. Since the visual representation is saved as an aspect within the network, it can easily be reused to layout additional networks in the same style without modification.

A key feature of the aspect-oriented design of the CX format is to allow the definition of custom aspects. Therefore, the RCX package was designed with a focus on extensibility with additional functions for the creation, modification, conversion, and validation of custom aspects.

Detailed documentation and examples can be found in the package manual and vignettes. The Supplementary Materials contain code examples for working with RCX networks and their creation.

## 3 Implementation

The RCX package builds upon on several R packages for data processing and graph representation. The CX networks are read and written with the readr package. The obtained JSON is transformed and further processed using the jsonlite (Ooms 2014) and tidyr packages (Wickham 2011).

The visualization was realized with the JavaScript library cytoscape.js (Franz et al. 2016) and a custom script to map the visual properties between the in CX used Cytoscape properties to cytoscape.js compatible layout definitions.

## 4 Conclusion

The RCX package is a freely available R software tool that enables the lossless conversion between the object-oriented JSON format of the CX data structure and the table-like paradigm of data in R. The data model was designed to enhance usability and enrich functionality by a better adjustment to fundamental R data structures and adding high-level functions for data manipulation.

Integrated conversion to igraph and Bioconductor compatible graph objects fosters the accessibility to advanced network analysis tools. Furthermore, extensibility was increased by facilitating the creation of custom aspects that cover specialized extensions to the CX data model.

By implementing this software, we ease the task of handling network data available via NDEx within the R Framework for Statistical Computing. Enriched networks as results of investigations and their visualizations can be easily created and translated to the CX format, that connects analysis, visualization, and collaboration.

## Supporting information

RCX package overview

Creating RCX from scratch

Extending the RCX Data Model

The RCX and CX Data Model

RCX Reference Manual

RCX Cheat Sheet

