## Supplementary material for "RCX – an R package adapting the Cytoscape Exchange format for biological networks": RCX package overview

***Florian J. Auer***[1*]

[1]Augsburg of University

[*]

**2021-10-26**

### Contents

1 Introduction
  - 1.1 The Cytoscape Exchange (CX)
  - 1.2 The NDEx platform
  - 1.3 Cytoscape
  - 1.4 RCX - an adaption of the CX format

2 Installation

3 The basics
  - 3.1 Read and write CX files
  - 3.2 Explore the RCX object
  - 3.3 Visualize the network
  - 3.4 Validation

4 Get information about the networks

5 Conversion to R graph data models
  - 5.1 igraph

6 Bioconductor graph (graphNEL)

7 Session info

### 1 Introduction

Networks are a powerful and flexible methodology for expressing biological knowledge for computation and communication. Biological networks can hold a variety of different types of information, like genetic or metabolic interactions, gene, and transcriptional regulation, protein-protein interaction (PPI), or cell signaling networks and pathways. They often form a valuable resource for hypothesis generation and further investigations, and in the course of the analyses, they are processed and enriched with additional information from experiments. As a result further networks are generated, whether as intermediate results that should be documented in the process, as the outcome of those analyses, or as visual representations and illustrations used in reports and publications. As a consequence, these resulting networks do not follow anymore the strict rules the source networks were subjected, therefore a more flexible format is needed to capture their content.

In addition, suitable solutions for transmission conflict with those for storage, or usage in applications and analyses. Therefore, seamless conversion between those different formats becomes as important as the data itself.

#### 1.1 The Cytoscape Exchange (CX)

A possible solution as flexible transmission format is provided by the Cytoscape Exchange (CX) format. CX is a JSON-based, aspect-oriented data structure, which means that the network is divided into several independent modules (“aspects”). This way, every aspect of a network, meaning nodes, edges, its attributes, and visual representations can be handled individually without interference. Each aspect has its own schema for its contained information, and links between aspects are realized by referencing the unique internal IDs of other aspects. The CX data model was developed by the NDEx project, in collaboration with the Cytoscape Consortium (http://www.cytoscapeconsortium.org/) as a transmission format between their tools, and since adopted by many others. More details about the CX data model can be found on its documentation website: https://home.ndexbio.org/data-model/

#### 1.2 The NDEx platform

The Network Data Exchange, or NDEx, is an online commons for biological networks (Pratt et al., 2015, Cell Systems 1, 302-305, October 28, 2015 ©2015 Elsevier Inc. ScienceDirect). It is an open-source software framework to manipulate, store, and exchange networks of various types and formats. NDEx can be used to upload, share and publicly distribute networks while providing an output in formats, that can be used by plenty of other applications.

The public NDEx server is a network data commons that provides pathway collections like the Pathway Interaction Database of the NCI (http://www.ndexbio.org/#/user/301a91c6-a37b-11e4-bda0-000c29202374) and the Cancer Cell Maps Initiative (http://www.ndexbio.org/#/user/b47268a6-8112-11e6-b0a6-06603eb7f303). Public networks can be searched and retrieved from the platform for further use. Own networks can be uploaded and shared with certain collaborators or groups privately or provided publicly to the community. Furthermore, private installation of the NDEx platform can be used to store and collaborate on networks locally.

The ndexr package available on Bioconductor (https://doi.org/doi:10.18129/B9.bioc.ndexr) allows connecting with the NDEx platform from within R. This package provides an interface to query the public NDEx server, as well as private installations, to upload, download or modify biological networks.

#### 1.3 Cytoscape

The most prominent software environment for biological network analysis and visualization is Cytoscape (https://cytoscape.org/). It provides support for large networks and comes with a rich set of features for custom visualization, and advanced layout and analysis algorithms. One of these visualization features is the “attribute-to-visual mapping”, where the network’s data translates to its visual representation. Based on this visualization strategy, Cytoscape contributed aspects to the CX-format to capture the visual representation as part of the network. Because of these aspects, the visualization not only can be documented along with the network, but also reproduced on other platforms, and even shared between networks with the same attributes used for creating the visualization.

#### 1.4 RCX - an adaption of the CX format

CX is a JSON-based data structure designed as a flexible model for transmitting networks with a focus on flexibility, modularity, and extensibility. Although those features are widely used in common REST protocols they don’t quite fit the R way of thinking about data.

This package provides an adaption of the CX format to standard R data formats and types to create and modify, load, export, and visualize those networks. This document aims to help the user to install and benefit from the wide range of functionality of this implementation. For an overview of the differences of the RCX implementation to the CX specification see Appendix: The RCX and CX Data Model

### 2 Installation

For installing packages from github the `devtools` package is the most common approach. However, it requires XML libraries installed on the system which can cause problems while installation due to unmet dependencies. The `remotes` package covers the functionality to download and install R packages stored in 'GitHub', 'GitLab', 'Bitbucket', 'Bioconductor', or plain 'subversion' or 'git' repositories without depending on XML libraries. If `devtools` is already installed, of course it can be used, otherwise it is recommended to use the lightweight `remotes` package.

**From github using remotes:**

```
if (!"remotes" %in% installed.packages()) install.packages("remotes")
if (!"RCX" %in% installed.packages()) remotes::install_github("frankkramer-lab/RCX")
library(RCX)
```

**From github using devtools:**

```
if (!"devtools" %in% installed.packages()) install.packages("devtools")
if (!"RCX" %in% installed.packages()) devtools::install_github("frankkramer-lab/RCX")
library(RCX)
```

### 3 The basics

In the following, it will be explained, how to read and write networks from/to CX files, create and modify RCX networks, validate its contents and finally visualize them.

#### 3.1 Read and write CX files

Networks can be downloaded from the NDEx plattform as CX files in JSON format. Those files can be read, and are automatically transformed into RCX networks that can be used in R. Here we load a provided example network from file:

```
cxFile <- system.file(
  "extdata",
  "Imatinib-Inhibition-of-BCR-ABL-66a902f5-2022-11e9-bb6a-0ac135e8bacf.cx",
  package = "RCX"
)

rcx = readCX(cxFile)
```

This network also can be accessed and downloaded from NDEx at https://www.ndexbio.org/viewer/networks/66a902f5-2022-11e9-bb6a-0ac135e8bacf

RCX networks can be saved in a similar manner:

```
writeCX(rcx, "path/to/some-file.cx")
```

However, there might some errors occur while reading CX file. This might happen, because the definition of the CX has changed over time, and so the definition of some aspects. Therefore it is possible, that there are still some networks stored at the NDEx platform following a deprecated format. In those cases it might be helpful to process the CX network step by step:

**1.** just read the JSON content without parsing

```
json <- readJSON(cxFile)

substr(json, 1, 77)
```

```
## [{"numberVerification":[{"longNumber":281474976710655}]},{"metaData":[{"name"
```

This also allows to handle a CX network in JSON format, even if it comes from a different source instead of a file.

**2.** parse the JSON

```
aspectList <- parseJSON(json)

str(aspectList, 2)
```

```
## List of 12
##  $ :List of 1
##   ..$ numberVerification:List of 1
##  $ :List of 1
##   ..$ metaData:List of 9
##  $ :List of 1
##   ..$ provenanceHistory:List of 1
##  $ :List of 1
##   ..$ nodes:List of 75
##  $ :List of 1
##   ..$ edges:List of 159
##  $ :List of 1
##   ..$ networkAttributes:List of 10
##  $ :List of 1
##   ..$ nodeAttributes:List of 1129
##  $ :List of 1
```

```
##   ..$ edgeAttributes:List of 229
##  $ :List of 1
##   ..$ cartesianLayout:List of 75
##  $ :List of 1
##   ..$ cyVisualProperties:List of 3
##  $ :List of 1
##   ..$ cyHiddenAttributes:List of 1
##  $ :List of 1
##   ..$ status:List of 1
```

The result of the parsing are nested lists containing all aspects and its contents. This format not easy to handle in R, but allows error correction previous to forming aspect and RCX objects.

**3.** process the aspect data

```
rcx <- processCX(aspectList)
```

All the above function for processing CX networks come with an option to show the performed steps. This might be helpful for finding occurring errors:

```
rcx <- readCX(cxFile, verbose = TRUE)
```

```
## Read file "RCX/extdata/Imatinib-Inhibition-of-BCR-ABL-66a902f5-2022-11e9-bb6a-0ac135e8bac
## Parse json...done!
## Parsing nodes...create aspect...done!
## Create RCX from parsed nodes...done!
## Parsing edges...create aspect...done!
## Add aspect "edges" to RCX...done!
## Parsing node attributes...create aspect...done!
## Add aspect "nodeAttributes" to RCX...done!
## Parsing edge attributes...create aspect...done!
## Add aspect "edgeAttributes" to RCX...done!
## Parsing network attributes...create aspect...done!
## Add aspect "networkAttributes" to RCX...done!
## Parsing cartesian layout...create aspect...done!
## Add aspect "cartesianLayout" to RCX...done!
## Parsing Cytoscape visual properties...done!
## - Create sub-objects...done!
## - Create aspect...done!
## Add aspect "cyVisualProperties" to RCX...done!
## Parsing Cytoscape hidden attributes...create aspect...done!
## Add aspect "cyHiddenAttributes" to RCX...done!
## Parsing meta-data...done!
## Ignore "numberVerification" aspect, not necessary in RCX!
## Can't process aspect "numberVerification", so skip it...done!
## Don't know what to do with a "provenanceHistory" aspect!
## Can't process aspect "provenanceHistory", so skip it...done!
## Ignore "status" aspect, not necessary in RCX!
## Can't process aspect "status", so skip it...done!
```

This shows, that some aspects that are contained in the CX file are ignored while creating the RCX network. Those are for example aspects needed for transmission of the CX (`status`, `numberVerification`), or deprecated aspects (`provenanceHistory`). For more details about the differences in aspects see Appendix: The RCX and CX Data Model.

#### 3.2 Explore the RCX object

The simplest way to have a look at the content of an RCX object is by printing it:

```
print(rcx)
## OR:
rcx
```

However, especially for large networks this can produce long and hardly readable output. To get a better overview of the contained aspects, the mandatory and automatically generated `metaData` aspect provides information about the contained aspects. This includes information about the number of elements or the highest used ID, if an aspect uses internal IDs:

```
rcx$metaData
```

```
## Meta-data:
##                  name version idCounter elementCount consistencyGroup
## 1               nodes     1.0     11551           75                1
## 2               edges     1.0     11554          159                1
## 3      nodeAttributes     1.0        NA         1129                1
## 4      edgeAttributes     1.0        NA          229                1
## 5   networkAttributes     1.0        NA           10                1
## 6     cartesianLayout     1.0        NA           75                1
## 7 cyVisualProperties     1.0        NA            3                1
## 8 cyHiddenAttributes     1.0        NA            1                1
```

Besides exploring the RCX-object manually, a summary of the object, or single aspects, can provide more insight on them:

```
summary(rcx$nodeAttributes)
```

```
##       propertyOf        name                 value
##  Total      : 1129   Length:1129        Boolean:185
##  Unique ids:   75    Unique:34          Double : 92
##  Min.      :11321    Class :character   Integer: 68
##  Max.      :11551                       String :784
```

We already can quickly see that there are many different node attributes are used. The different node attributes are:

```
unique(rcx$nodeAttributes$name)
```

```
##  [1] "sbo"                    "metaId"                 "compartmentCode"
##  [4] "sbml type"              "sbml id"                "reversible"
##  [7] "sbml compartment"       "cyId"                   "label"
## [10] "sbml type ext"          "fast"                   "constant"
## [13] "units"                  "boundaryCondition"      "derivedUnits"
## [16] "sbml initial amount"    "value"                  "substanceUnits"
## [19] "hasOnlySubstanceUnits" "uniprot"                "inchikey"
## [22] "chebi"                  "biocyc"                 "cas"
## [25] "chemspider"             "kegg.compound"          "inchi"
## [28] "pubchem.compound"       "hmdb"                   "scale"
## [31] "exponent"               "kind"                   "multiplier"
## [34] "unitSid"
```

#### 3.3 Visualize the network

This package provides simple functions to visualize the network encoded in the RCX object.

```
visualize(rcx)
```

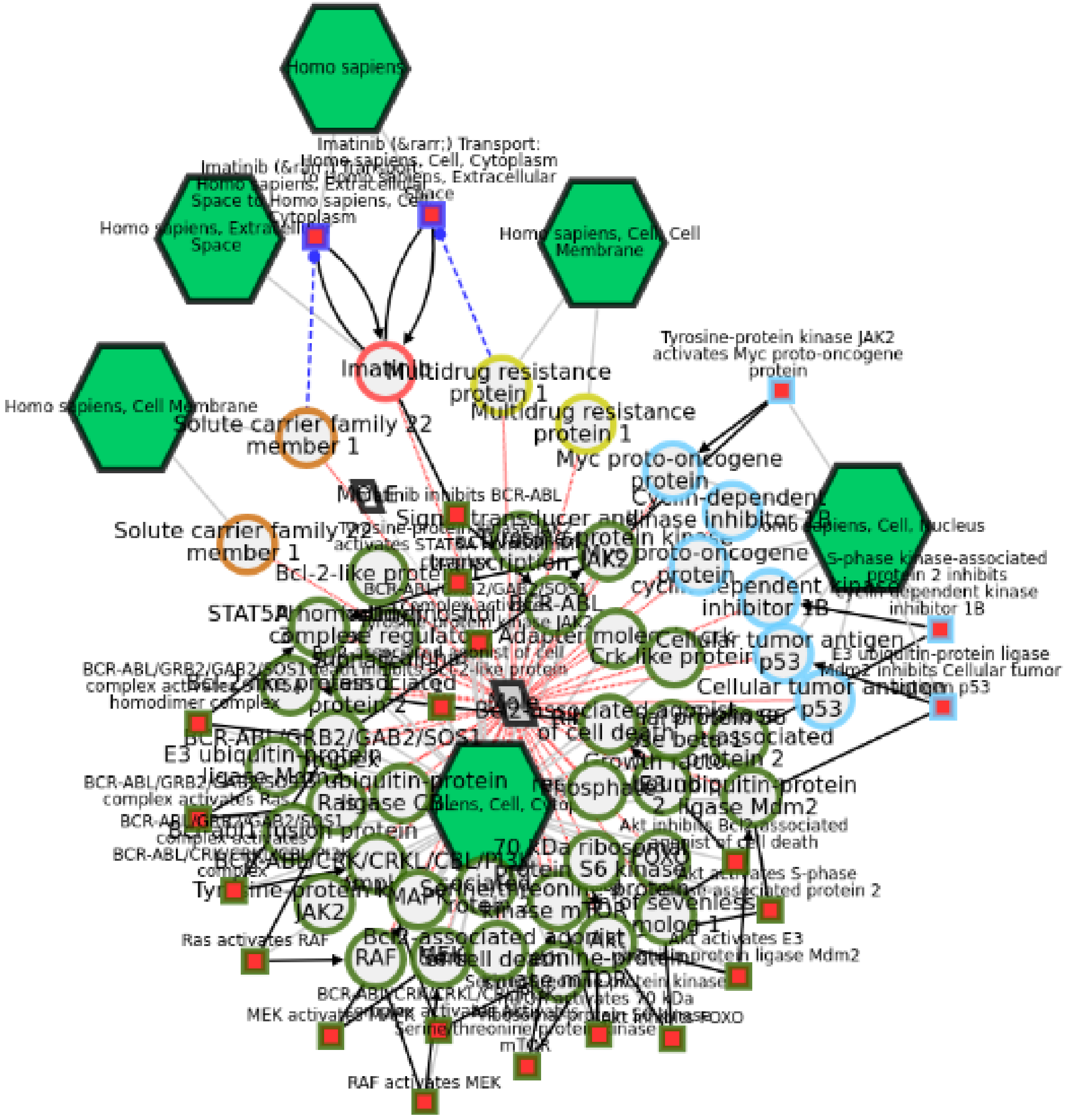


Imatinib Inhibition of BCR-ABL (66a902f5-2022-11e9-bb6a-0ac135e8bacf)

The visualization also utilizes the same JavaScript library as the NDEx platform. Therefore the visual result is the same as when the network is uploaded to the NDEx platform. Additionally, this allows the visualization to be exported as a single HTML file, which can directly be hosted on a web-server or included in existing websites:

```
writeHTML(rcx, "path/to/some-file.html")
```

Networks with many nodes and edges, or those without a provided layout, often are difficult to interpret visually. To untangle the "hairball" it is possible to apply different layout option provided by the Cytoscape.js framework. A force driven layout is a good starting point in those cases. For demonstration purposes, let's delete the visual layout of our network first:

```r
## save them for later
originalVisualProperties <- rcx$cyVisualProperties

## and delete them from the RCX network
rcx$cyVisualProperties <- NULL
rcx <- updateMetaData(rcx)

rcx$metaData
```

```
## Meta-data:
##                   name version idCounter elementCount consistencyGroup
## 1                nodes     1.0     11551           75                1
## 2                edges     1.0     11554          159                1
## 3       nodeAttributes     1.0        NA         1129                1
## 4       edgeAttributes     1.0        NA          229                1
## 5    networkAttributes     1.0        NA           10                1
## 6      cartesianLayout     1.0        NA           75                1
## 7 cyHiddenAttributes     1.0        NA            1                1
```

Lets have a look at the visualization now:

```r
visualize(rcx, layout = c(name = "cose"))
```

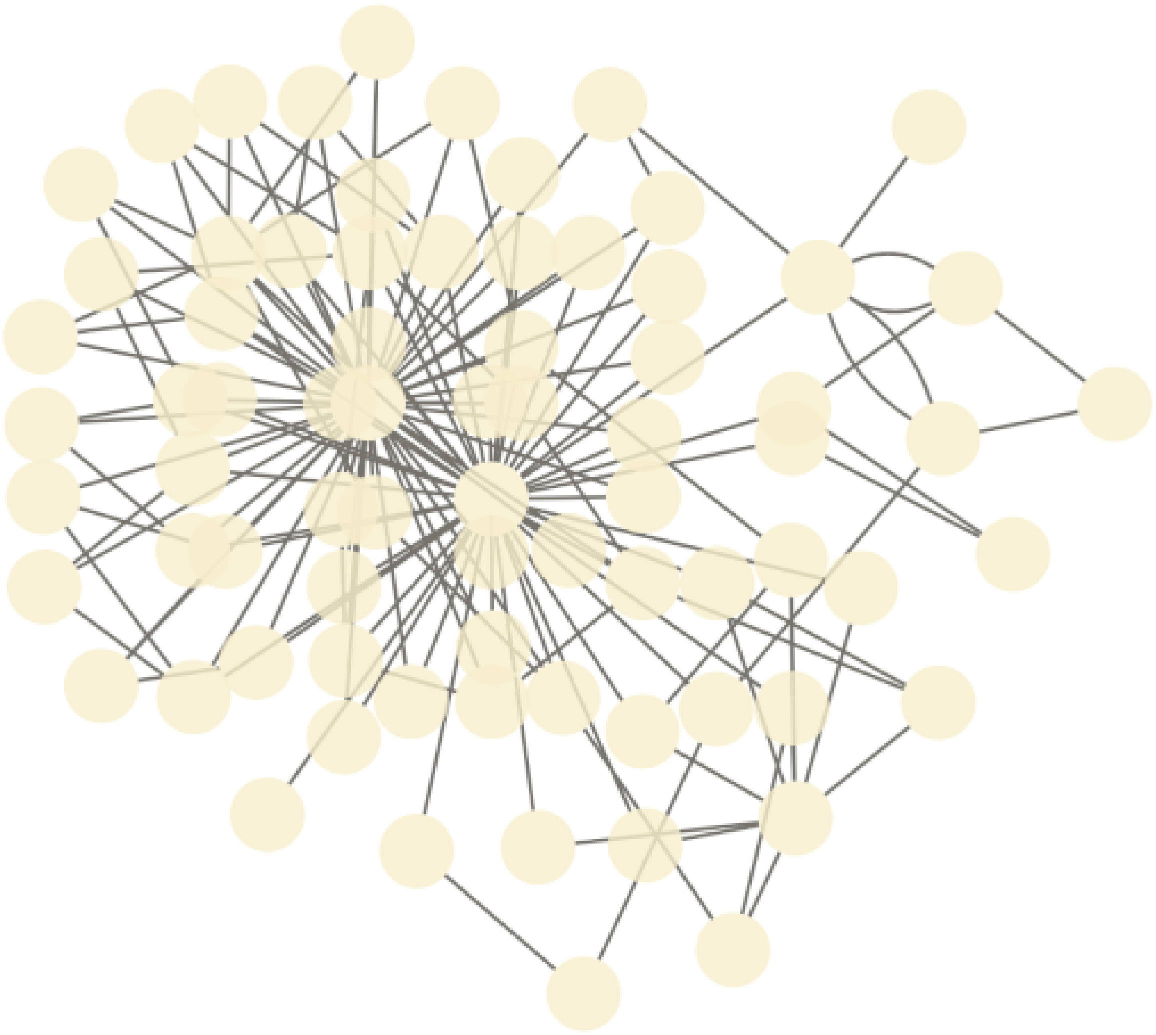

Force-driven visualization of the network without visual properties

Unfortunately no labels are shown in the network. We can fix that by defining a simple pass-through mapping for the node labels and add it to the network:

```
cyMapping <- createCyVisualPropertyMappings(
  name= "NODE_LABEL" ,
  type = "PASSTHROUGH",
  definition = "COL=label,T=string"
)

cyVisualProperties <- createCyVisualProperties(
  defaultNodes = createCyVisualProperty(
    mappings = cyMapping
  )
)

rcx <- updateCyVisualProperties(rcx, cyVisualProperties)
```

By default the visualization is opened in RStudio, but it also can be forced to open in an external browser. The `Cytoscape.js` parameters used to customize the layout algorithm can be found in it documentation at https://js.cytoscape.org/#layouts/cose

```
visualize(
  rcx,
  layout = c(
    name="cose",
    idealEdgeLength="80",
    edgeElasticity="250"),
  openExternal = TRUE)
```

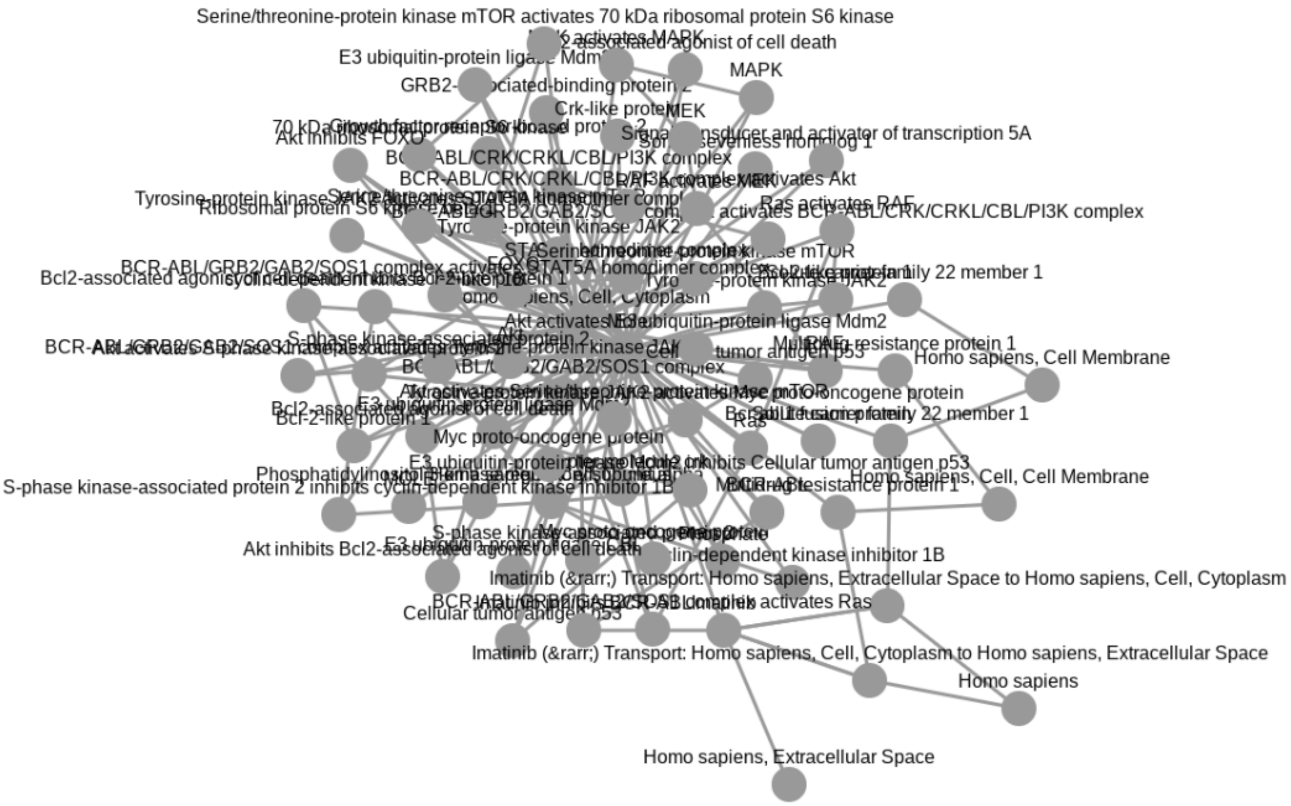

#### 3.4 Validation

The correctness of the RCX network is important for the conversion to CX, and therefore to be used at all platforms and tools. The validity of the an RCX network can be checked simply:

```
validate(rcx)
```

```
## Checking Nodes Aspect:
## - Is object of class "NodesAspect"...OK
## - All required columns present (id)...OK
## - Column (id) doesn't contain any NA values...OK
## - At least one node present...OK
## - Column (id) contains only unique values...OK
## - Column (id) only contains numeric values...OK
## - Column (id) only contains positive (>=0) values...OK
## - No merge artefacts present (i.e. column with old ids: oldId)...OK
## - Only allowed columns present (id, name, represents)...OK
## >> Nodes Aspect: OK
##
## Checking Edges Aspect:
## - Is object of class "EdgesAspect"...OK
## - All required columns present (id, source, target)...OK
## - Column (id) doesn't contain any NA values...OK
## - Column (id) contains only unique values...OK
## - Column (id) only contains numeric values...OK
## - Column (id) only contains positive (>=0) values...OK
## - Column (source) doesn't contain any NA values...OK
## - Column (source) only contains numeric values...OK
## - Column (source) only contains positive (>=0) values...OK
## - Column (target) doesn't contain any NA values...OK
## - Column (target) only contains numeric values...OK
## - Column (target) only contains positive (>=0) values...OK
## - No merge artefacts present (i.e. column with old ids: oldId)...OK
## - Only allowed columns present (id, source, target, name, interaction)...OK
## >> Edges Aspect: OK
##
## Checking Node Attributes Aspect:
## - Is object of class "NodeAttributesAspect"...OK
## - All required columns present (propertyOf, name, value, dataType, isList)...OK
## - Combination of columns (propertyOf, name) contains only unique values...OK
## - Column (propertyOf) doesn't contain any NA values...OK
## - Column (propertyOf) only contains numeric values...OK
## - Column (propertyOf) only contains positive (>=0) values...OK
## - Is the column (name) a character vector...OK
## - Is the column (value) a list...OK
## - Column (dataType) doesn't contain any NA values...OK
## - Is the column (dataType) a character vector...OK
## - All values of dataType are in the allowed set (boolean, integer, long, double, string).
## - Column (isList) doesn't contain any NA values...OK
## - Column (isList) only contains logical values...OK
## - Only allowed columns present (propertyOf, name, value, dataType, isList)...OK
## >> Node Attributes Aspect: OK
##
## Checking Edge Attributes Aspect:
## - Is object of class "EdgeAttributesAspect"...OK
## - All required columns present (propertyOf, name, value, dataType, isList)...OK
## - Combination of columns (propertyOf, name) contains only unique values...OK
```

```
## - Column (propertyOf) doesn't contain any NA values...OK
## - Column (propertyOf) only contains numeric values...OK
## - Column (propertyOf) only contains positive (>=0) values...OK
## - Is the column (name) a character vector...OK
## - Is the column (value) a list...OK
## - Column (dataType) doesn't contain any NA values...OK
## - Is the column (dataType) a character vector...OK
## - All values of dataType are in the allowed set (boolean, integer, long, double, string).
## - Column (isList) doesn't contain any NA values...OK
## - Column (isList) only contains logical values...OK
## - Only allowed columns present (propertyOf, name, value, dataType, isList)...OK
## >> Edge Attributes Aspect: OK
##
## Checking Network Attributes Aspect:
## - Is object of class "NetworkAttributesAspect"...OK
## - All required columns present (name, value, dataType, isList)...OK
## - Column (name) contains only unique values...OK
## - Is the column (name) a character vector...OK
## - Is the column (value) a list...OK
## - Column (dataType) doesn't contain any NA values...OK
## - Is the column (dataType) a character vector...OK
## - All values of dataType are in the allowed set (boolean, integer, long, double, string).
## - Column (isList) doesn't contain any NA values...OK
## - Column (isList) only contains logical values...OK
## - Only allowed columns present (name, value, dataType, isList)...OK
## >> Network Attributes Aspect: OK
##
## Checking Cartesian Layout Aspect:
## - Is object of class "CartesianLayoutAspect"...OK
## - All required columns present (node, x, y)...OK
## - Column (node) contains only unique values...OK
## - Column (node) only contains numeric values...OK
## - Column (node) doesn't contain any NA values...OK
## - Column (x) only contains numeric values...OK
## - Column (x) doesn't contain any NA values...OK
## - Column (y) only contains numeric values...OK
## - Column (y) doesn't contain any NA values...OK
## - Only allowed columns present (node, x, y, z)...OK
## >> Cartesian Layout Aspect: OK
##
## Checking Cytoscape Visual Properties Aspect:
## - Is object of class "CyVisualPropertiesAspect"...OK
## - Is object a list...OK
## - Is a named list ("network", "nodes", "edges", "defaultNodes" or "defaultEdges")...OK
## - The list only contains entries of class "CyVisualProperty"...OK
## - At least one of the elements present (network, nodes, edges, defaultNodes, defaultEdges
## For defaultNodes: Checking Cytoscape Visual Property sub-aspect:
## - Is object of class "CyVisualProperty"...OK
## - Is object a list...OK
## - Is a named list ("appliesTo", "view", "properties", "dependencies" or "mappings")...OK
## - List element (appliesTo) only contains numeric values...OK
## - List element (view) only contains numeric values...OK
## - Combination of columns (appliesTo, view) contains only unique values...OK
## - At least one of the elements present (properties, dependencies, mappings)...OK
##   Checking Cytoscape Visual Property Mappings:
## - Is mappings a list...OK
## - The list only contains entries of class "CyVisualPropertyMappings"...OK
## - All required columns present (name, type, definition)...OK
## - All list elements of name contain only character values...OK
## - All list elements of name don't contain any NA values...OK
## - All list elements (name) contain only unique values...OK
```

```
## - All list elements of type contain only character values...OK
## - All list elements of type don't contain any NA values...OK
## - All list elements of definition contain only character values...OK
## - All list elements of definition don't contain any NA values...OK
## - Only allowed columns present (name, type, definition)...OK
## >> Cytoscape Visual Property sub-aspect: OK
## >> Cytoscape Visual Properties Aspect: OK
##
## Checking Cytoscape Hidden Attributes Aspect:
## - Is object of class "CyHiddenAttributesAspect"...OK
## - All required columns present (name, value, dataType, isList)...OK
## - Column (name) contains only unique values...OK
## - Is the column (name) a character vector...OK
## - Is the column (value) a list...OK
## - Column (dataType) doesn't contain any NA values...OK
## - Is the column (dataType) a character vector...OK
## - All values of dataType are in the allowed set (boolean, integer, long, double, string).
## - Column (isList) doesn't contain any NA values...OK
## - Column (isList) only contains logical values...OK
## - Only allowed columns present (name, value, dataType, isList)...OK
## >> Cytoscape Hidden Attributes Aspect: OK
##
## Checking RCX:
## - Is object of class "RCX"...OK
## - nodes aspect is present...OK
## - Validate nodes aspect...OK
## - Validate edges aspect...OK
##   - Reference aspect (nodes) present and correct...OK
##   - All id references exist (EdgesAspect$source ids in NodesAspect$id)...OK
##   - All id references exist (EdgesAspect$target ids in NodesAspect$id)...OK
## - Validate node attributes aspect...OK
##   - Reference aspect (nodes) present and correct...OK
##   - All id references exist (NodeAttributesAspect$propertyOf ids in NodesAspect$id)...OK
## - Validate edge attributes aspect...OK
##   - Reference aspect (edges) present and correct...OK
##   - All id references exist (EdgeAttributesAspect$propertyOf ids in EdgesAspect$id)...OK
## - Validate network attributes aspect...OK
## - Validate cartesian layout aspect...OK
##   - Reference aspect (nodes) present and correct...OK
##   - All id references exist (CartesianLayoutAspect$node ids in NodesAspect$id)...OK
## - Validate cytoscape visual property aspect...OK
## - Validate cytoscape hidden attributes aspect...OK
## >> RCX: OK
```

The verbose output also shows at which tests are performed. Let's manually introduce an error in an aspect and validate only this one again. Therefore we simply duplicate a node ID.

```
nodes <- rcx$nodes
nodes$id[1] <- nodes$id[2]

test <- validate(nodes)
```

```
## Checking Nodes Aspect:
## - Is object of class "NodesAspect"...OK
## - All required columns present (id)...OK
## - Column (id) doesn't contain any NA values...OK
## - At least one node present...OK
## - Column (id) contains only unique values...FAIL
## - Column (id) only contains numeric values...OK
```

```
## - Column (id) only contains positive (>=0) values...OK
## - No merge artefacts present (i.e. column with old ids: oldId)...OK
## - Only allowed columns present (id, name, represents)...OK
## >> Nodes Aspect: FAIL
```

```
test
```

```
## [1] FALSE
```

As expected, the test fails and returns `FALSE`.

### 4 Get information about the networks

It is always useful to get some basic information about the network. For example this can be the number of elements in the different aspects of the network:

```
countElements(rcx)
```

```
##              NodesAspect            MetaDataAspect               EdgesAspect
##                       75                        NA                       159
##     NodeAttributesAspect      EdgeAttributesAspect   NetworkAttributesAspect
##                     1129                       229                        10
##    CartesianLayoutAspect CyHiddenAttributesAspect CyVisualPropertiesAspect
##                       75                         1                         1
```

This also works for single aspects:

```
countElements(rcx$nodes)
```

```
## [1] 75
```

To determine, if an aspect contains IDs (on the contrary to knowing it beforehand), this can be checked with:

```
hasIds(rcx$nodes)
```

```
## [1] TRUE
```

If an aspect has IDs, one can check what the highest used ID is, to know at which ID the next elements have to continue before adding them. This can even be done for the whole network at once.

```
maxId(rcx$nodes)
```

```
## [1] 11551
```

```
maxId(rcx)
```

```
## NodesAspect EdgesAspect
##       11551       11554
```

Since we now know, that the nodes aspect has IDs, we can also determine the name of the property that holds the IDs:

```
idProperty(rcx$nodes)
```

```
## [1] "id"
```

Other aspects use those IDs to reference to them. Let’s find out, which aspect is referred by others:

```
referredBy(rcx)
```

```
## $NodesAspect
## [1] "EdgesAspect"           "NodeAttributesAspect"  "CartesianLayoutAspect"
##
## $EdgesAspect
## [1] "EdgeAttributesAspect"
```

The nodes aspect is referred by the edges aspect, so we can find out which properties of the edges aspect refer to it:

```
refersTo(rcx$edges)
```

```
##        source        target
## "NodesAspect" "NodesAspect"
```

It might have gotten to your attention, that there is a difference between the aspect name and the aspect class. This has been done intentionally to avoid naming conflicts. The different naming can be converted to each other:

```
## all classes
aspectClasses
```

```
##                        rcx                   metaData
##                      "RCX"           "MetaDataAspect"
##                      nodes                      edges
##              "NodesAspect"              "EdgesAspect"
##             nodeAttributes             edgeAttributes
##     "NodeAttributesAspect"     "EdgeAttributesAspect"
##          networkAttributes            cartesianLayout
##  "NetworkAttributesAspect"    "CartesianLayoutAspect"
##                   cyGroups         cyVisualProperties
##           "CyGroupsAspect" "CyVisualPropertiesAspect"
##         cyHiddenAttributes         cyNetworkRelations
## "CyHiddenAttributesAspect" "CyNetworkRelationsAspect"
##               cySubNetworks              cyTableColumn
##      "CySubNetworksAspect"      "CyTableColumnAspect"
```

```
## class of nodes
aspectName2Class("nodes")
```

```
##         nodes
## "NodesAspect"
```

```
## accession name of NodesAspect
aspectClass2Name("NodesAspect")
```

```
## [1] "nodes"
```

```r
## back and forth
class(rcx[[aspectClass2Name("NodesAspect")]])
```

```
## [1] "NodesAspect" "data.frame"
```

**Note:** These function can help especially for writing extension to the RCX data model (see 03 Extending the RCX Data Model).

### 5 Conversion to R graph data models

The RCX package provides conversion to and from objects of iGraph and Bioconductor graph, both widely used libraries for graph manipulation and network analysis.

#### 5.1 igraph

igraph is a library for creating and manipulating graphs and analyzing (especially large) networks. RCX networks can be simply converted to igraph objects as follows:

```r
library(igraph)
```

```
##
## Attaching package: 'igraph'

## The following objects are masked from 'package:stats':
##
##     decompose, spectrum

## The following object is masked from 'package:base':
##
##     union
```

```r
ig <- toIgraph(rcx)
summary(ig)
```

```
## IGRAPH 9c4456e UN-- 75 159 --
## + attr: attribute...name (g/c), description (g/c), name (v/n), id
## | (v/n), sbo (v/c), metaId (v/c), cartesianLayout...x (v/n),
## | cartesianLayout...y (v/n), id (e/n), interaction (e/c), sbo (e/c),
## | stoichiometry (e/c)
```

To ensure a consistent conversion in both direction, some conventions have to be matched: For example avoiding collisions between `name` used in nodes, `name` used as nodes attribute, and the `name` used by igraph, the first two were re-named in igraph to `nodeName` and `attribute...name` respectively. A similar convention is used for the cartesian coordinates: In the igraph object they are saved as `cartesianLayout...x`, `cartesianLayout...y`, and `cartesianLayout...z`. If the vertex attributes match this naming convention, they assigned to the correct aspect when converted back:

```r
rcxFromIg <- fromIgraph(ig)
```

Igraph objects can not hold information about the visual representation of the network, and therefore the Cytoscape aspects are lost in the conversion. But we can simply add the `CyVisualProperties` aspect we saved previously to get back to the original layout:

```
rcxFromIg <- updateCyVisualProperties(rcxFromIg, originalVisualProperties)
```

```
visualize(rcxFromIg)
```

### 6 Bioconductor graph (graphNEL)

Bioconductor graph is a package that implements some simple graph handling capabilities. It can handle multi-edges, but only if the graph is directed and the source and target start and end not between the same nodes. Unfortunaltelly this is the case in our sample network. A quick fix is simply switching the direction of source and target for the multi-edges:

```
dubEdges <- duplicated(rcx$edges[c("source", "target")])

s <- rcx$edges$source
rcx$edges$source[dubEdges] <- rcx$edges$target[dubEdges]
rcx$edges$target[dubEdges] <- s[dubEdges]

gNel <- toGraphNEL(rcx, directed = TRUE)
rcxBack <- fromGraphNEL(gNel)
```

Then we can simply convert the RCX to a graphNEL object:

```
gNel <- toGraphNEL(rcx, directed = TRUE)
gNel
```

```
## A graphNEL graph with directed edges
## Number of Nodes = 75
## Number of Edges = 159
```

For the conversion igraph is used as an intermediate format, therefore, the same conventions apply as for the igraph conversion. The conversion back to the RCX object works analogously:

```
rcxBack <- fromGraphNEL(gNel)
rcxBack$metaData
```

```
## Meta-data:
##                 name version idCounter elementCount consistencyGroup
## 1              nodes     1.0     11551           75                1
## 2              edges     1.0     11554          159                1
## 3     nodeAttributes     1.0        NA          148                1
## 4     edgeAttributes     1.0        NA          229                1
## 5  networkAttributes     1.0        NA            2                1
## 6    cartesianLayout     1.0        NA           75                1
```

As igraph, graphNEL objects can not hold information about the visual representation of the network, so here too we can restore the original layout by adding the `CyVisualProperties` aspect we saved previously.

### 7 Session info

```
sessionInfo()
```

```
## R version 4.0.3 (2020-10-10)
## Platform: x86_64-pc-linux-gnu (64-bit)
## Running under: Ubuntu 20.04.3 LTS
##
## Matrix products: default
## BLAS:   /usr/lib/x86_64-linux-gnu/blas/libblas.so.3.9.0
## LAPACK: /usr/lib/x86_64-linux-gnu/lapack/liblapack.so.3.9.0
##
## locale:
##  [1] LC_CTYPE=en_US.UTF-8       LC_NUMERIC=C
##  [3] LC_TIME=de_DE.UTF-8        LC_COLLATE=en_US.UTF-8
##  [5] LC_MONETARY=de_DE.UTF-8    LC_MESSAGES=en_US.UTF-8
##  [7] LC_PAPER=de_DE.UTF-8       LC_NAME=C
##  [9] LC_ADDRESS=C               LC_TELEPHONE=C
## [11] LC_MEASUREMENT=de_DE.UTF-8 LC_IDENTIFICATION=C
##
## attached base packages:
## [1] stats     graphics  grDevices utils     datasets  methods   base
##
## other attached packages:
## [1] igraph_1.2.7     RCX_0.99.0       knitr_1.36       BiocStyle_2.18.1
##
## loaded via a namespace (and not attached):
##  [1] graph_1.68.0        Rcpp_1.0.7          magrittr_2.0.1
##  [4] BiocGenerics_0.36.1 rlang_0.4.12        fastmap_1.1.0
##  [7] stringr_1.4.0       plyr_1.8.6          tools_4.0.3
## [10] parallel_4.0.3      xfun_0.27           jquerylib_0.1.4
## [13] htmltools_0.5.2     yaml_2.2.1          digest_0.6.28
## [16] bookdown_0.24       BiocManager_1.30.16 formatR_1.11
## [19] evaluate_0.14       rmarkdown_2.11      stringi_1.7.5
## [22] compiler_4.0.3      stats4_4.0.3        jsonlite_1.7.2
## [25] pkgconfig_2.0.3
```
