## Supplementary material for "RCX – an R package adapting the Cytoscape Exchange format for biological networks": Creating RCX from scratch

### 02. Creating RCX from scratch

***Florian J. Auer***[1*]

[1]Augsburg of University

[*]

**2021-10-26**

### Contents

1 The Cytoscape Exchange (CX) Format
2 Starting with nodes and edges
3 Adding attributes to the network
3.1 Node Attributes
3.2 Network Attributes
4 Put the nodes into position
5 Create visual layout
5.1 Visual properties of the network
5.2 Visual properties of nodes
5.3 Visual properties of edges
5.4 Create a visual property aspect
6 Table column
7 Visualize the final network
8 Meta-data
9 Create RCX at once
10 Save the network
11 Session info

### 1 The Cytoscape Exchange (CX) Format

CX is an “aspect-oriented” data structure, which means that the network is divided into several independent modules (“aspects”). Each aspect has its own schema for its contained information, and links between aspects are realized by referencing the unique internal IDs of other aspects.

The CX format comes with some pre-defined aspects, which are divided in three groups:

- core aspects, with which the most properties of the network can be defined
  - nodes
  - edges
  - nodeAttributes
  - edgeAttributes
  - networkAttributes
  - cartesianLayout
- cytoscape aspects, which provide additional information to visualize the networks with Cytoscape
  - cyGroups
  - cyVisualProperties
  - cyHiddenAttributes
  - cyNetworkRelations
  - cySubnetworks
  - cyTableColumn
- meta aspects, which are necessary for data transmission
  - metaData
  - status

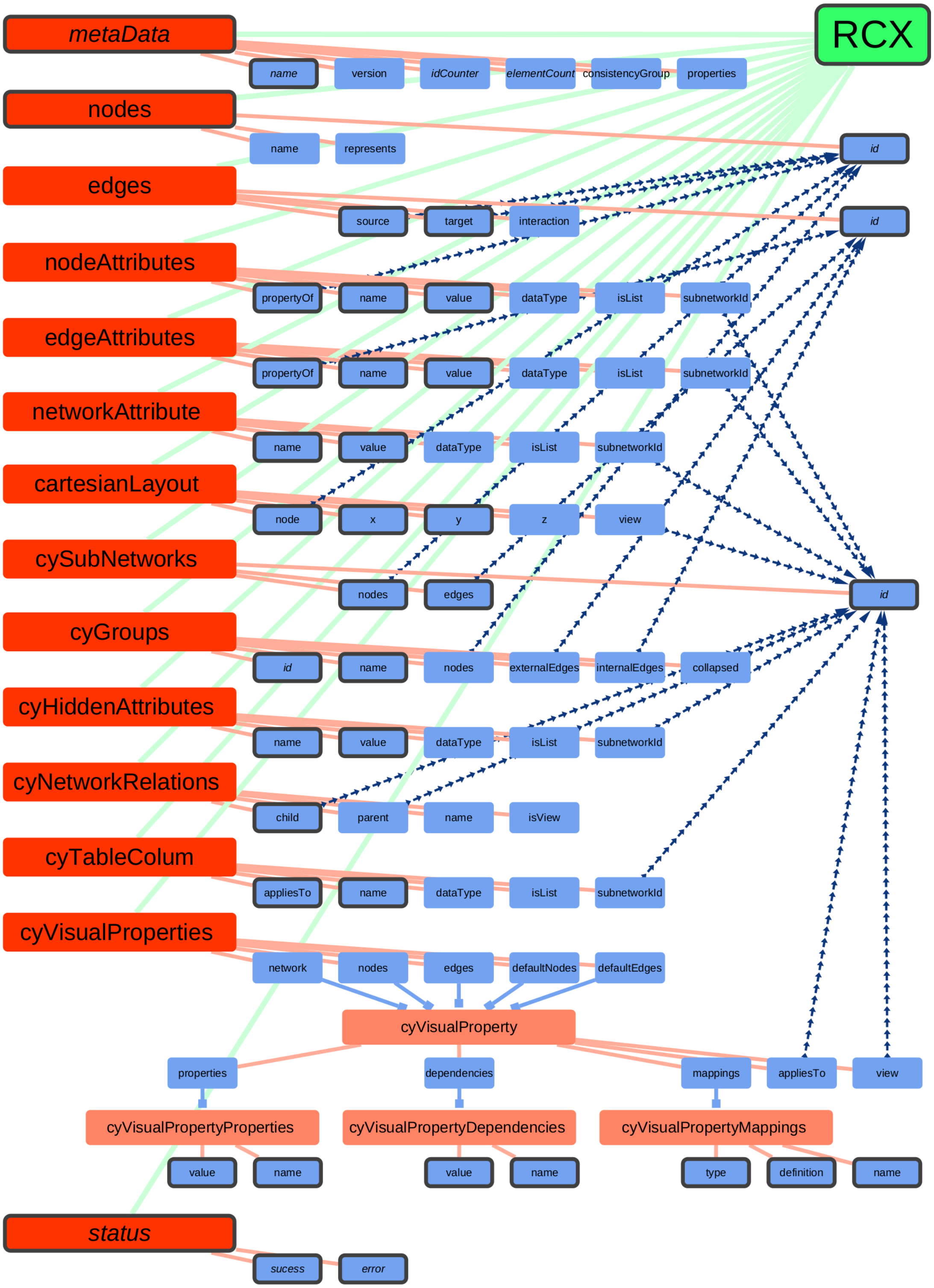


RCX data structure

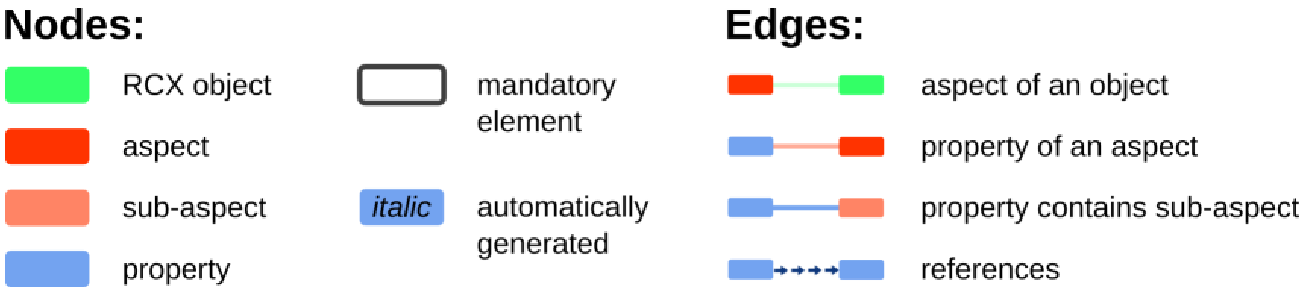


Legend

This network is available at the NDEx platform at https://www.ndexbio.org/viewer/networks/ebdda4da-2ca5-11ec-b3be-0ac135e8bacf

Aspects are the elemental units composing an RCX object. They hold different properties, some are mandatory to the aspect, some are optional. Being mandatory does not necessary mean, that those properties need to be specified at creation. For example, some aspects hold IDs to be referenced by other aspects. Although those IDs are mandatory for the aspect, they will be automatically created if not provided.

For a full documentation of the differences between the RCX and CX, see the vignette Appendix: The RCX and CX Data Model.

In the following, it will be explained, how to create the above RCX network from scratch and save it as CX file ready to be uploaded to the NDEx platform.

### 2 Starting with nodes and edges

Every RCX network needs at least one node, but it is a good starting point to begin the construction of a new network with the creation of nodes and edges. Firstly we create the nodes:

```
nodes <- createNodes(
  name = c(
    "RCX",
    "metaData", "name", "version", "idCounter", "elementCount", "consistencyGroup", "properti
    "nodes", "id", "name", "represents",
    "edges", "id", "source", "target", "interaction",
    "nodeAttributes", "propertyOf", "name", "value", "dataType", "isList", "subnetworkId",
    "edgeAttributes", "propertyOf", "name", "value", "dataType", "isList", "subnetworkId",
    "networkAttribute", "name", "value", "dataType", "isList", "subnetworkId",
    "cartesianLayout", "node", "x", "y", "z", "view",
    "cySubNetworks", "id", "nodes", "edges",
    "cyGroups", "id", "name", "nodes", "externalEdges", "internalEdges", "collapsed",
    "cyHiddenAttributes", "name", "value", "dataType", "isList", "subnetworkId",
    "cyNetworkRelations", "child", "parent", "name", "isView",
    "cyTableColum", "appliesTo", "name", "dataType", "isList", "subnetworkId",
    "cyVisualProperties", "network", "nodes", "edges", "defaultNodes", "defaultEdges",
    "cyVisualProperty", "properties", "dependencies", "mappings", "appliesTo", "view",
    "cyVisualPropertyProperties", "value", "name",
    "cyVisualPropertyDependencies", "value", "name",
    "cyVisualPropertyMappings", "type", "definition", "name",
    "status", "sucess", "error"
  )
)

head(nodes)
```

```
## Nodes:
##   id         name
## 1  0          RCX
## 2  1     metaData
## 3  2         name
## 4  3      version
## 5  4    idCounter
## 6  5 elementCount
```

Next we can create the edges, that link the nodes by its IDs used as source and target of the edge. Additionally, the interaction described by an edge can be specified:

```
edges <- createEdges(
    source = c(
        1, 2, 3, 4, 5, 6, 7, 8, 9, 10, 11, 12, 13, 14, 15, 16, 17, 18, 19, 20,
        21, 22, 23, 24, 25, 26, 27, 28, 29, 30, 31, 32, 33, 34, 35, 36, 37,
        38, 39, 40, 41, 42, 43, 44, 45, 46, 47, 48, 49, 50, 51, 52, 53, 54,
        55, 56, 57, 58, 59, 60, 61, 62, 63, 64, 65, 66, 67, 68, 69, 70, 71,
        72, 73, 74, 75, 76, 78, 79, 80, 81, 82, 84, 85, 87, 88, 90, 91, 92,
        93, 94, 95, 14, 15, 18, 23, 25, 30, 36, 38, 42, 45, 46, 50, 51, 52,
        59, 61, 62, 70, 72, 73, 74, 75, 76, 78, 79, 80, 81, 82
    ),
    target = c(
        0, 1, 1, 1, 1, 1, 1, 0, 8, 8, 8, 0, 12, 12, 12, 12, 0, 17, 17, 17, 17,
        17, 17, 0, 24, 24, 24, 24, 24, 24, 0, 31, 31, 31, 31, 31, 0, 37, 37,
        37, 37, 37, 0, 43, 43, 43, 0, 47, 47, 47, 47, 47, 47, 0, 54, 54, 54,
        54, 54, 0, 60, 60, 60, 60, 0, 65, 65, 65, 65, 65, 0, 71, 71, 71, 71,
        71, 77, 77, 77, 77, 77, 83, 83, 86, 86, 89, 89, 89, 0, 93, 93, 9, 9,
        9, 44, 13, 44, 44, 9, 44, 9, 13, 9, 13, 13, 44, 44, 44, 44, 77, 77,
        77, 77, 77, 83, 86, 89, 44, 44
    ),
    interaction = c(
        "aspectOf", "propertyOf", "propertyOf", "propertyOf", "propertyOf",
        "propertyOf", "propertyOf", "aspectOf", "propertyOf", "propertyOf",
        "propertyOf", "aspectOf", "propertyOf", "propertyOf", "propertyOf",
        "propertyOf", "aspectOf", "propertyOf", "propertyOf", "propertyOf",
        "propertyOf", "propertyOf", "propertyOf", "aspectOf", "propertyOf",
        "propertyOf", "propertyOf", "propertyOf", "propertyOf", "propertyOf",
        "aspectOf", "propertyOf", "propertyOf", "propertyOf", "propertyOf",
        "propertyOf", "aspectOf", "propertyOf", "propertyOf", "propertyOf",
        "propertyOf", "propertyOf", "aspectOf", "propertyOf", "propertyOf",
        "propertyOf", "aspectOf", "propertyOf", "propertyOf", "propertyOf",
        "propertyOf", "propertyOf", "propertyOf", "aspectOf", "propertyOf",
        "propertyOf", "propertyOf", "propertyOf", "propertyOf", "aspectOf",
        "propertyOf", "propertyOf", "propertyOf", "propertyOf", "aspectOf",
        "propertyOf", "propertyOf", "propertyOf", "propertyOf", "propertyOf",
        "aspectOf", "propertyOf", "propertyOf", "propertyOf", "propertyOf",
        "propertyOf", "propertyOf", "propertyOf", "propertyOf", "propertyOf",
        "propertyOf", "propertyOf", "propertyOf", "propertyOf", "propertyOf",
        "propertyOf", "propertyOf", "propertyOf", "aspectOf", "propertyOf",
        "propertyOf", "references", "references", "references", "references",
        "references", "references", "references", "references", "references",
        "references", "references", "references", "references", "references",
        "references", "references", "references", "references", "contains",
        "contains", "contains", "contains", "contains", "contains", "contains",
        "contains", "references", "references"
    )
)

head(edges)
```

```
## Edges:
##   id source target interaction
## 1  0      1      0    aspectOf
## 2  1      2      1  propertyOf
## 3  2      3      1  propertyOf
## 4  3      4      1  propertyOf
```

```
## 5  4      5      1  propertyOf
## 6  5      6      1  propertyOf
```

With the nodes and edges, the RCX object can be created:

```
rcx <- createRCX(nodes = nodes, edges = edges)
```

To investigate the created RCX network, we can simply show the summary of the network:

```
summary(rcx)
```

```
## $nodes
##       id          name
##  Total:96   Length:96
##  Min. : 0   Class :character
##  Max. :95   Mode  :character
##
## $metaData
##       name             version            idCounter      elementCount
##  Length:2           Length:2           Min.   : 95.0   Min.   : 96.0
##  Class :character   Class :character   1st Qu.:100.8   1st Qu.:101.8
##  Mode  :character   Mode  :character   Median :106.5   Median :107.5
##                                        Mean   :106.5   Mean   :107.5
##                                        3rd Qu.:112.2   3rd Qu.:113.2
##                                        Max.   :118.0   Max.   :119.0
##  consistencyGroup
##  Min.   :1
##  1st Qu.:1
##  Median :1
##  Mean   :1
##  3rd Qu.:1
##  Max.   :1
##
## $edges
##       id              source            target    interaction
##  Total:119   Total     :119   Total     :119   Length:119
##  Min. :  0   Unique ids: 91   Unique ids: 22   Class :character
##  Max. :118   Min.      :  1   Min.      :  0   Mode  :character
##              Max.      : 95   Max.      : 93
```

Here we already see that, additionally to the nodes and edges also the metaData is created. Let's have a closer look at the meta data:

```
rcx$metaData
```

```
## Meta-data:
##    name version idCounter elementCount consistencyGroup
## 1 nodes     1.0        95           96                1
## 2 edges     1.0       118          119                1
```

This meta aspect is created and updated automatically when the RCX object is modified and gives an overview of the present aspects, number of elements those contain, and the highest IDs used in the single aspects.

The resulting network already can be visualized, but since no coordinates are set, by default, all nodes would clump together at the same position. To avoid this, we can use a force directed layout to visualize the network:

```
visualize(rcx, layout = c(name = "cose"))
```

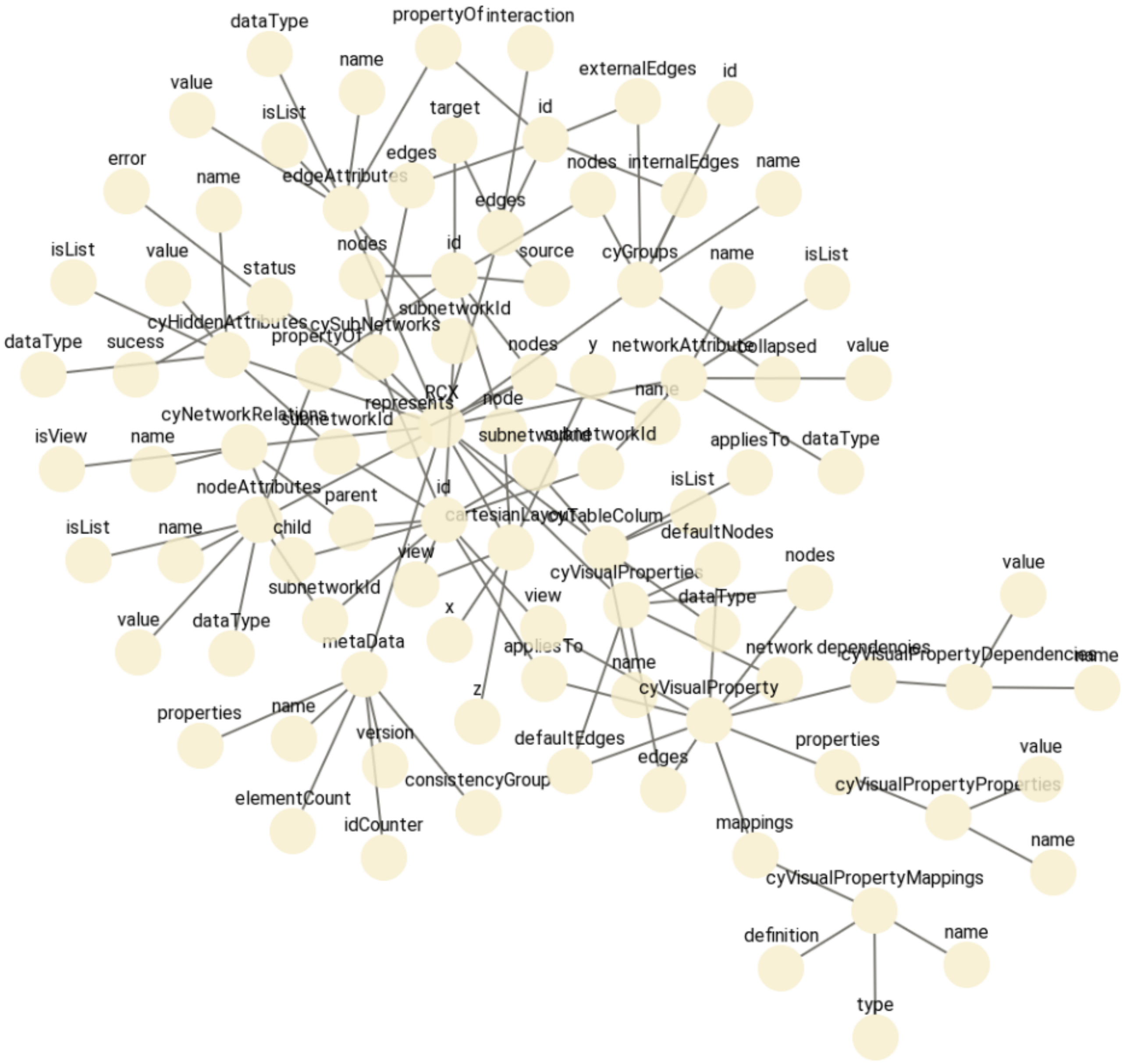


Force directed layout for RCX network with only nodes and edges

### 3 Adding attributes to the network

#### 3.1 Node Attributes

The firstly, let's add the node types. The nodes are either the RCX object, the aspects or the properties of the aspects. Those types are later used to color and size the nodes appropriately.

```
nodeAttributesType = createNodeAttributes(
  propertyOf = nodes$id,
  name = rep("type", length(nodes$id)),
  value = c(
    "object",
    "aspect", "property", "property", "property", "property", "property", "property",
    "aspect", "property", "property", "property",
    "aspect", "property", "property", "property", "property",
    "aspect", "property", "property", "property", "property", "property", "property",
    "aspect", "property", "property", "property", "property", "property", "property",
    "aspect", "property", "property", "property", "property", "property",
    "aspect", "property", "property", "property", "property", "property",
    "aspect", "property", "property", "property",
    "aspect", "property", "property", "property", "property", "property", "property",
    "aspect", "property", "property", "property", "property", "property",
    "aspect", "property", "property", "property", "property",
    "aspect", "property", "property", "property", "property", "property",
    "aspect", "property", "property", "property", "property", "property",
    "subAspect", "property", "property", "property", "property", "property",
    "subAspect", "property", "property",
    "subAspect", "property", "property",
    "subAspect", "property", "property", "property",
    "aspect", "property", "property"
  )
)

rcx = updateNodeAttributes(rcx, nodeAttributesType)
```

Next, mark those aspects and properties that are required in creation of the aspect and RCX object. To simplify the creation of the node attributes aspect, firstly only create a vector containing the node IDS.

```
nodesRequired <- c(
  0,    # RCX
  1,    # metaData
  2,    # metaData:name
  8,    # metaData:nodes
  9,    # metaData:id
  13,   # edge:id
  14,   # edge:source
  15,   # edge:target
  18,   # nodeAttributes:propertyOf
  19,   # nodeAttributes:name
  20,   # nodeAttributes:value
  25,   # edgeAttributes:propertyOf
  26,   # edgeAttributes:name
  27,   # edgeAttributes:value
```

```
  32,   # networkAttribute:name
  33,   # networkAttribute:value
  38,   # cartesianLayout:node
  39,   # cartesianLayout:x
  40,   # cartesianLayout:y
  44,   # cySubNetworks:id
  45,   # cySubNetworks:nodes
  46,   # cySubNetworks:edges
  48,   # cyGroups:id
  49,   # cyGroups:name
  55,   # cyHiddenAttributes:name
  56,   # cyHiddenAttributes:value
  61,   # cyNetworkRelations:child
  66,   # cyTableColum:appliesTo
  67,   # cyTableColum:name
  84,   # cyVisualPropertyProperties:value
  85,   # cyVisualPropertyProperties:name
  87,   # cyVisualPropertyDependencies:value
  88,   # cyVisualPropertyDependencies:name
  90,   # cyVisualPropertyMappings:type
  91,   # cyVisualPropertyMappings:definition
  92,   # cyVisualPropertyMappings:name
  93,   # status
  94,   # status:sucess
  95    # status:error
)
```

Also mark those aspects and properties that are automatically created, when aspects and RCX objects are created. Here also only a vector containing the node IDs is created to keep it simple.

```
nodesAuto <- c(
  1,    # metaData
  2,    # metaData:name
  4,    # metaData:idCounter
  5,    # metaData:elementCount
  9,    # nodes:id
  13,   # edges:id
  44,   # cySubNetworks:id
  48,   # cyGroups:id
  93,   # status
  94,   # status:sucess
  95    # status:error
)
```

Now create the node attributes, combine them to one node attributes aspect and add the aspect to the RCX object. Of course this would be possible in one combined step, but for demonstration purposes it is done here step by step.

The node attributes to mark are simply set to `TRUE`. The remaining node attributes could be set to `FALSE`, but it is not really needed to do so. If you want to assign a mapping to node attributes, that are not `TRUE` you would be required to set them. With not setting them, they are not accessible for the mapping of values (since no value is defined), so the layout falls back to the default style. Since boolean values only have two possible values this only makes a difference, where you define the visual styles: Either in the default style or for both values in the mapping. The later option, as mentioned before, requires you to include the `FALSE` values as well, which increases the size of the CX file.

```
nodeAttributesRequired <- createNodeAttributes(
    propertyOf = nodesRequired, name = rep("required", length(nodesRequired)),
    value = rep(TRUE, length(nodesRequired))
)
```

```
nodeAttributesAuto <- createNodeAttributes(
    propertyOf = nodesAuto, name = rep("auto", length(nodesAuto)),
    value = rep(TRUE, length(nodesAuto))
)

nodeAttributes <- updateNodeAttributes(nodeAttributesRequired, nodeAttributesAuto)
rcx <- updateNodeAttributes(rcx, nodeAttributes)

## nodeAttributesType was already in rcx, so save it all
nodeAttributes <- rcx$nodeAttributes
```

Adding EdgeAttributes would work similarly, but here we have included all needed information already in the `interaction` property of the edges.

#### 3.2 Network Attributes

Some attributes follow a convention on NDEx, especially in the network attributes. The attributes name, author and description are shown on the NDEx platform along the network or in the list overview of your networks.

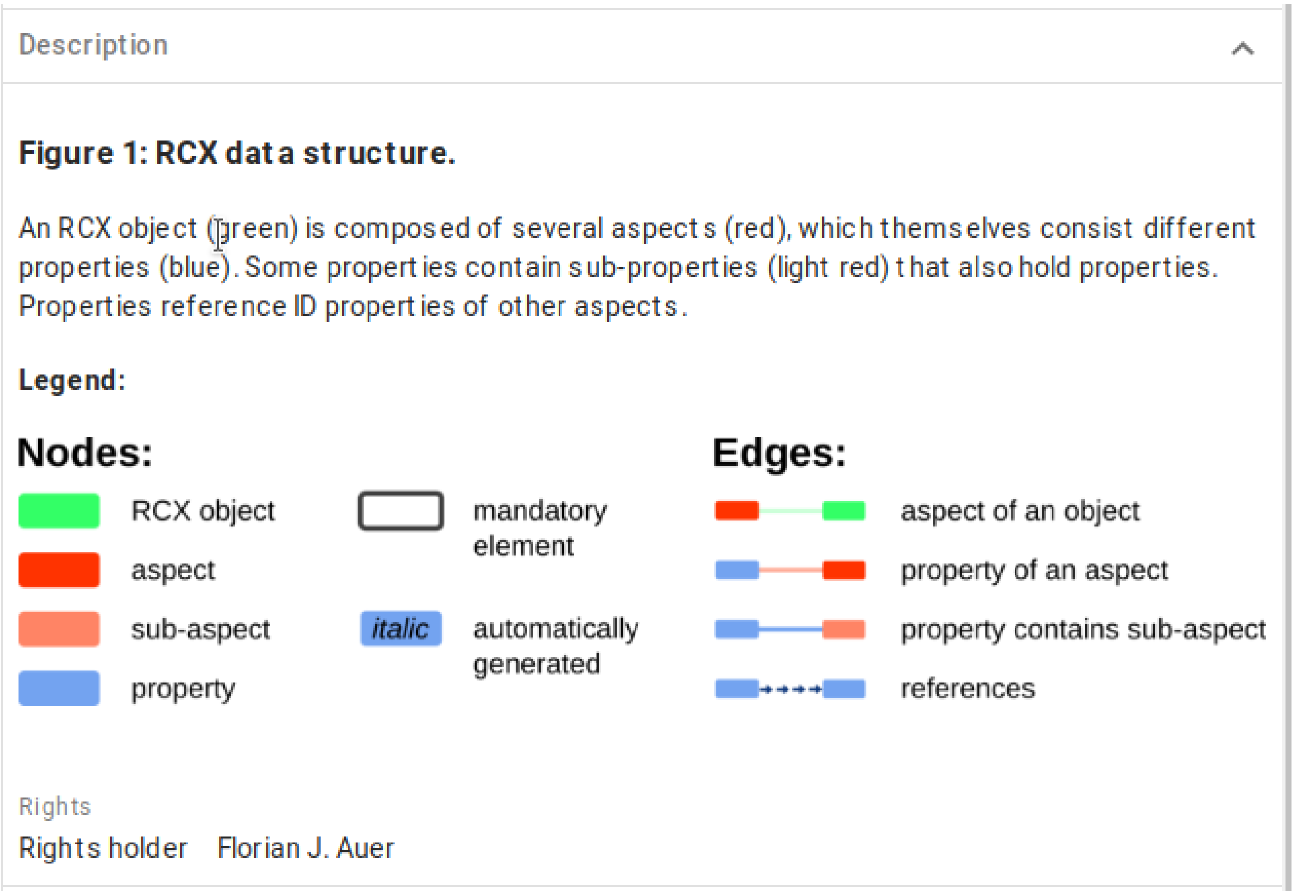


Network attributes shown on the NDEx platform

Therefore, it is useful to include those in the network. The description of the network can be formatted using HTML tags. The NDEx platform also includes an editor for the network attributes. Here we also include the image of the legend in the description of the network. Therefore, we have to do some HTML magic and convert the image to a base64 encoding representation of it.

```r
legendImgFile <- "RCX_Data_Structure_Legend.png"
legendImgRaw <- readBin(legendImgFile, "raw", file.info(legendImgFile)[1, "size"])
legendImgBase64 <- base64enc::base64encode(legendImgRaw)

description <- paste0(
  '<h3>Figure 1: RCX data structure.</h3>
  <p>An RCX object (green) is composed of several aspects (red),
  which themself consist different properties (blue).
  Some properties contain sub-properties (light red) that also hold properties.
  Properties reference ID properties of other aspects.</p>
  <h4>Legend:</h4>
  <p>
  
  </p>'
  )

networkAttributes <- createNetworkAttributes(
  name = c(
    "name",
    "author",
    "description"
  ),
  value = c(
    "RCX Data Structure",
    "Florian J. Auer",
    description
  )
)

rcx <- updateNetworkAttributes(rcx, networkAttributes)
```

Now that we have included the node and network attributes, let's check the network meta-data again to make sure everything worked out fine.

```r
rcx$metaData
```

```
## Meta-data:
##                name version idCounter elementCount consistencyGroup
## 1             nodes     1.0        95           96                1
## 2             edges     1.0       118          119                1
## 3    nodeAttributes     1.0        NA          146                1
## 4 networkAttributes     1.0        NA            3                1
```

### 4 Put the nodes into position

Until now, the nodes don't have any position yet, which means that the layout of the network did not change and only can be layouted using a visualization together with some layout algorithm. Instead, we can set the position of the nodes manually using the cartesian layout aspect.

```
posX <- c(
  1187, # RCX
  219, 430, 540, 650, 760, 870, 980, # metaData
  219, 1189, 431, 541, # nodes
  219, 1189, 545, 655, 765, # edges
  219, 435, 545, 655, 765, 875, 985, # nodeAttributes
  219, 435, 545, 655, 765, 875, 985, # edgeAttributes
  219, 435, 545, 655, 765, 875, # networkAttribute
  219, 435, 545, 655, 765, 875, # cartesianLayout
  219, 1201, 545, 655, # cySubNetworks
  219, 435, 545, 655, 765, 875, 985, # cyGroups
  219, 435, 545, 655, 765, 875, # cyHiddenAttributes
  219, 435, 545, 655, 765, # cyNetworkRelations
  219, 435, 545, 655, 765, 875, # cyTableColum
  219, 435, 545, 655, 765, 875, # cyVisualProperties
  655, 326, 656, 986, 1096, 1206, # cyVisualProperty
  326, 325, 435, # cyVisualPropertyProperties
  656, 655, 765, # cyVisualPropertyDependencies
  986, 985, 1095, 1205, # cyVisualPropertyMappings
  219, 435, 545 # status
)

posY <- c(
  596, # RCX
  595, 646, 646, 646, 646, 646, 646, # metaData
  692, 743, 743, 743, # nodes
  790, 836, 836, 836, 836, # edges
  889, 934, 934, 934, 934, 934, 934, # nodeAttributes
  985, 1031, 1031, 1031, 1031, 1031, 1031, # edgeAttributes
  1081, 1126, 1126, 1126, 1126, 1126, # networkAttribute
  1174, 1219, 1219, 1219, 1219, 1219, # cartesianLayout
  1270, 1316, 1316, 1316, # cySubNetworks
  1364, 1409, 1409, 1409, 1409, 1409, 1409, # cyGroups
  1460, 1506, 1506, 1506, 1506, 1506, # cyHiddenAttributes
  1557, 1602, 1602, 1602, 1602, # cyNetworkRelations
  1653, 1699, 1699, 1699, 1699, 1699, # cyTableColum
  1750, 1795, 1795, 1795, 1795, 1795, # cyVisualProperties
  1872, 1931, 1931, 1931, 1931, 1931, # cyVisualProperty
  2001, 2059, 2059, # cyVisualPropertyProperties
  2001, 2059, 2059, # cyVisualPropertyDependencies
  2001, 2059, 2059, 2059, # cyVisualPropertyMappings
  2136, 2182, 2182 # status
)

cartesianLayout <- createCartesianLayout(
  nodes$id,
  x=posX,
  y=posY)
```

```
rcx <- updateCartesianLayout(rcx, cartesianLayout)
```

Now we can visualize the network with the position assigned to the nodes:

```
visualize(rcx)
```

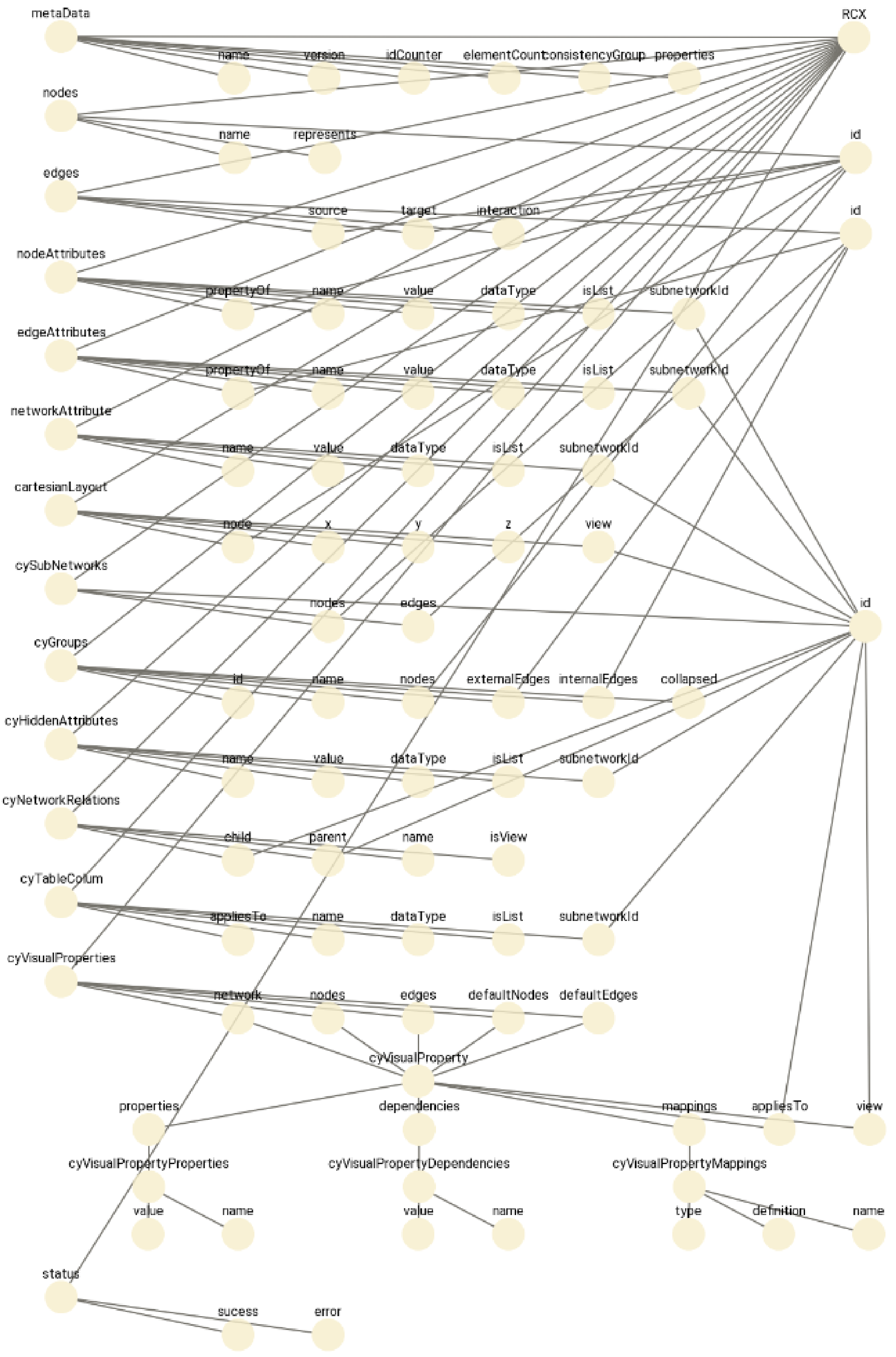


RCX network with a cartesian layout for nodes

### 5 Create visual layout

The visual representation of the network is defined in the `CyVisualProperties` aspect. This aspect holds the definitions for the network, nodes and edges in the form of a `CyVisualProperty` sub-aspect. In this sub-aspect, the layout is defined by

- properties: static properties of networks, nodes or edges (defined by a `CyVisualPropertyProperties` sub-aspect)
- mappings: definition of data dependent discrete, continuous and passthrough mappings (defined by a `CyVisualPropertyMappings` sub-aspect)
- dependencies: dependencies between visual properties (defined by a `CyVisualPropertyMappings` sub-aspect); Currently there are three dependencies supported:
  - Lock Node with and height: `nodeSizeLocked`
  - Fit Custom Graphics to node: `nodeCustomGraphicsSizeSync`
  - Edge color to arrows: `arrowColorMatchesEdge`

Each property (element in `CyVisualProperty`) can be applied to a subnetwork and/or view.

The complete structure of the visual properties looks as follows:

```
CyVisualProperties
├──network = CyVisualProperty
├──nodes = CyVisualProperty
├──edges = CyVisualProperty
├──defaultNodes = CyVisualProperty
└──defaultEdges = CyVisualProperty

CyVisualProperty
├──properties = CyVisualPropertyProperties
│    ├──name
│    └──value
├──dependencies = CyVisualPropertyDependencies
│    ├──name
│    └──value
├──mappings = CyVisualPropertyMappings
│    ├──name
│    ├──type
│    └──definition
├──appliesTo = <reference to subnetwork id>
└──view = <reference to subnetwork id>
```

In the following we will create the `CyVisualProperties` aspect and needed sub-aspects to define a proper visual representation.

#### 5.1 Visual properties of the network

For the network visual properties like the background color can be defined. Here we create the properties for a network and embed it into a new `CyVisualProperty` sub-aspect.

```
cyVisualPropertyPropertiesNetwork <- createCyVisualPropertyProperties(
  name = c(
    "NETWORK_BACKGROUND_PAINT",
    "NETWORK_EDGE_SELECTION",
    "NETWORK_NODE_SELECTION"
  ),
  value = c(
    "#FFFFFF",
    "true",
    "true"
  )
)

cyVisualPropertyNetwork <- createCyVisualProperty(
  properties = cyVisualPropertyPropertiesNetwork
)
```

#### 5.2 Visual properties of nodes

First we define the static properties for the nodes. This includes the default node color and size, the border color and with, etc.

```
cyVisualPropertyPropertiesNodes <- createCyVisualPropertyProperties(
  name = c(
    "NODE_BORDER_PAINT",
    "NODE_BORDER_STROKE",
    "NODE_BORDER_TRANSPARENCY",
    "NODE_BORDER_WIDTH",
    "NODE_DEPTH",
    "NODE_FILL_COLOR",
    "NODE_HEIGHT",
    "NODE_LABEL_COLOR",
    "NODE_LABEL_FONT_FACE",
    "NODE_LABEL_FONT_SIZE",
    "NODE_LABEL_POSITION",
    "NODE_LABEL_TRANSPARENCY",
    "NODE_LABEL_WIDTH",
    "NODE_PAINT",
    "NODE_SELECTED",
    "NODE_SELECTED_PAINT",
    "NODE_SHAPE",
    "NODE_SIZE",
    "NODE_TRANSPARENCY",
    "NODE_VISIBLE",
    "NODE_WIDTH"
  ),
  value = c(
    "#CCCCCC",
    "SOLID",
```

```
      "255",
      "0.0",
      "0.0",
      "#89D0F5",
      "40.0",
      "#000000",
      "SansSerif.plain,plain,12",
      "14",
      "C,C,c,0.00,0.00",
      "255",
      "200.0",
      "#1E90FF",
      "false",
      "#FFFF00",
      "ROUND_RECTANGLE",
      "35.0",
      "255",
      "true",
      "90.0"
    )
  )
```

Next we can define the mappings for the nodes. The node label is simply generated by passing the value of the column `name` through. Here, the other property values are discrete mappings, that only take a column by its name and the corresponding data type, and then simply assign new values to the some selected values from the column.

For example the height of the node is set to 50.0pt and 80.0pt for `aspect` and `object` values from the node attributes column `type`.

Here is the full definition of the node mappings:

```
  cyVisualPropertyMappingNodes <- createCyVisualPropertyMappings(
    name=c(
      "NODE_LABEL",
      "NODE_HEIGHT",
      "NODE_WIDTH",
      "NODE_FILL_COLOR",
      "NODE_BORDER_PAINT",
      "NODE_BORDER_WIDTH",
      "NODE_LABEL_FONT_FACE",
      "NODE_LABEL_FONT_SIZE"
    ),
    type = c(
      "PASSTHROUGH",
      "DISCRETE",
      "DISCRETE",
      "DISCRETE",
      "DISCRETE",
      "DISCRETE",
      "DISCRETE",
      "DISCRETE"
    ),
    definition = c(
      "COL=name,T=string",
      paste0("COL=type,T=string,",
             "K=0=aspect,V=0=50.0,",
             "K=1=object,V=1=80.0,"),
      paste0("COL=type,T=string,",
             "K=0=aspect,V=0=300.0,",
             "K=1=object,V=1=150.0,",
```

```
            "K=2=subAspect,V=2=300.0"),
    paste0("COL=type,T=string,",
            "K=0=aspect,V=0=#FF3300,",
            "K=1=property,V=1=#73A3F0,",
            "K=2=object,V=2=#33FF66,",
            "K=3=subAspect,V=3=#FF8465"),
    paste0("COL=required,T=boolean,",
            "K=0=true,V=0=#3F3F3F"),
    paste0("COL=required,T=boolean,",
            "K=0=true,V=0=10.0"),
    paste0("COL=auto,T=boolean,",
            "K=0=true,V=0=SansSerif.italic,,plain,,14"),
    paste0("COL=type,T=string,",
            "K=0=aspect,V=0=30,",
            "K=1=object,V=1=50,",
            "K=2=subAspect,V=2=20")
  )
)
```

As mentioned before, there are only a few dependencies supported. For nodes there are only the following:

```
cyVisualPropertyDependenciesNodes <- createCyVisualPropertyDependencies(
  name = c(
    "nodeCustomGraphicsSizeSync",
    "nodeSizeLocked"
  ),
  value = c(
    "true",
    "false"
  )
)
```

Finally combine the create sub-aspects to one `CyVisualProperty`. By default this is applied to the whole network and not limited to some subnetworks or views.

```
cyVisualPropertyNodes <- createCyVisualProperty(
  properties = cyVisualPropertyPropertiesNodes,
  mappings = cyVisualPropertyMappingNodes,
  dependencies = cyVisualPropertyDependenciesNodes
)
```

#### 5.3 Visual properties of edges

The definition of visual properties for edges works the same as for nodes. So let's begin again with the static properties:

```
cyVisualPropertyPropertiesEdges <- createCyVisualPropertyProperties(
  name = c(
    "EDGE_CURVED",
    "EDGE_LINE_TYPE",
    "EDGE_PAINT",
    "EDGE_SELECTED",
    "EDGE_SELECTED_PAINT",
    "EDGE_SOURCE_ARROW_SELECTED_PAINT",
    "EDGE_SOURCE_ARROW_SHAPE",
    "EDGE_SOURCE_ARROW_SIZE",
```

```
    "EDGE_SOURCE_ARROW_UNSELECTED_PAINT",
    "EDGE_STROKE_SELECTED_PAINT",
    "EDGE_STROKE_UNSELECTED_PAINT",
    "EDGE_TARGET_ARROW_SELECTED_PAINT",
    "EDGE_TARGET_ARROW_SHAPE",
    "EDGE_TARGET_ARROW_SIZE",
    "EDGE_TARGET_ARROW_UNSELECTED_PAINT",
    "EDGE_TRANSPARENCY",
    "EDGE_VISIBLE"
  ),
  value = c(
    "false",
    "SOLID",
    "#323232",
    "false",
    "#FFFF00",
    "#FFFF00",
    "NONE",
    "6.0",
    "#000000",
    "#FF0000",
    "#848484",
    "#FFFF00",
    "NONE",
    "6.0",
    "#000000",
    "255",
    "true"
  )
)
```

Add a mapping for the edges:

```
cyVisualPropertyMappingEdges <- createCyVisualPropertyMappings(
  name=c(
    "EDGE_LINE_TYPE",
    "EDGE_TARGET_ARROW_SHAPE",
    "EDGE_UNSELECTED_PAINT",
    "EDGE_WIDTH"
  ),
  type = c(
    "DISCRETE",
    "DISCRETE",
    "DISCRETE",
    "DISCRETE"
  ),
  definition = c(
    paste0("COL=interaction,T=string,",
           "K=0=references,V=0=SEPARATE_ARROW"),
    paste0("COL=interaction,T=string,",
           "K=0=contains,V=0=SQUARE,",
           "K=1=references,V=1=DELTA"),
    paste0("COL=interaction,T=string,",
           "K=0=propertyOf,V=0=#FFAD99,",
           "K=1=contains,V=1=#73A3F0,",
           "K=2=aspectOf,V=2=#CBFFD8,",
           "K=3=references,V=3=#0D377C"),
    paste0("COL=interaction,T=string,",
           "K=0=partOf,V=0=10.0,",
           "K=1=propertyOf,V=1=5.0,",
```

```
            "K=2=contains,V=2=5.0,",
            "K=3=aspectOf,V=3=7.0,",
            "K=4=references,V=4=3.0")
  )
)
```

And the one remaining dependency:

```
cyVisualPropertyDependenciesEdges <- createCyVisualPropertyDependencies(
  name = c(
    "arrowColorMatchesEdge"
  ),
  value = c(
    "true"
  )
)
```

And combine the sub-aspects to one `CyVisualProperty`.

```
cyVisualPropertyEdges <- createCyVisualProperty(
  properties=cyVisualPropertyPropertiesEdges,
  mappings = cyVisualPropertyMappingEdges,
  dependencies = cyVisualPropertyDependenciesEdges
)
```

#### 5.4 Create a visual property aspect

Above we create the `CyVisualProperty` sub-aspects for the network, nodes and edges. Finally, we have to combine those to a `CyVisualProperties` aspect, where usually the default properties are added as `defaultNodes` or `defaultEdges`. The final aspect now can be added to the network:

```
cyVisualProperties <- createCyVisualProperties(
  network = cyVisualPropertyNetwork,
  defaultNodes = cyVisualPropertyNodes,
  defaultEdges = cyVisualPropertyEdges
)

rcx <- updateCyVisualProperties(rcx, cyVisualProperties)
```

### 6 Table column

Before we are finished, we can add a table-column aspect to the network as well. This aspect simply holds the information about the node, edge and network attributes, as well as their data type. This seems a little redundant, since the same information can already be found in the network, but Cytoscape seems to depend on it, although it automatically creates this table on import.

Therefore, for completeness, we create the table-column aspect:

```
cyTableColumn <- createCyTableColumn(
  name = c("name", "type", "required", "auto",
           "name", "interaction",
           "name", "author", "description"),
```

```
  appliesTo = c("nodes", "nodes", "nodes", "nodes",
                "edges", "edges",
                "networks", "networks", "networks"),
  dataType = c("string", "string", "boolean", "boolean",
               "string", "string",
               "string", "string", "string")
)

rcx <- updateCyTableColumn(rcx, cyTableColumn)
```

### 7 Visualize the final network

Now that we are finished, we can visualize the network again, this time with the desired visual representation:

```
visualize(rcx)
```

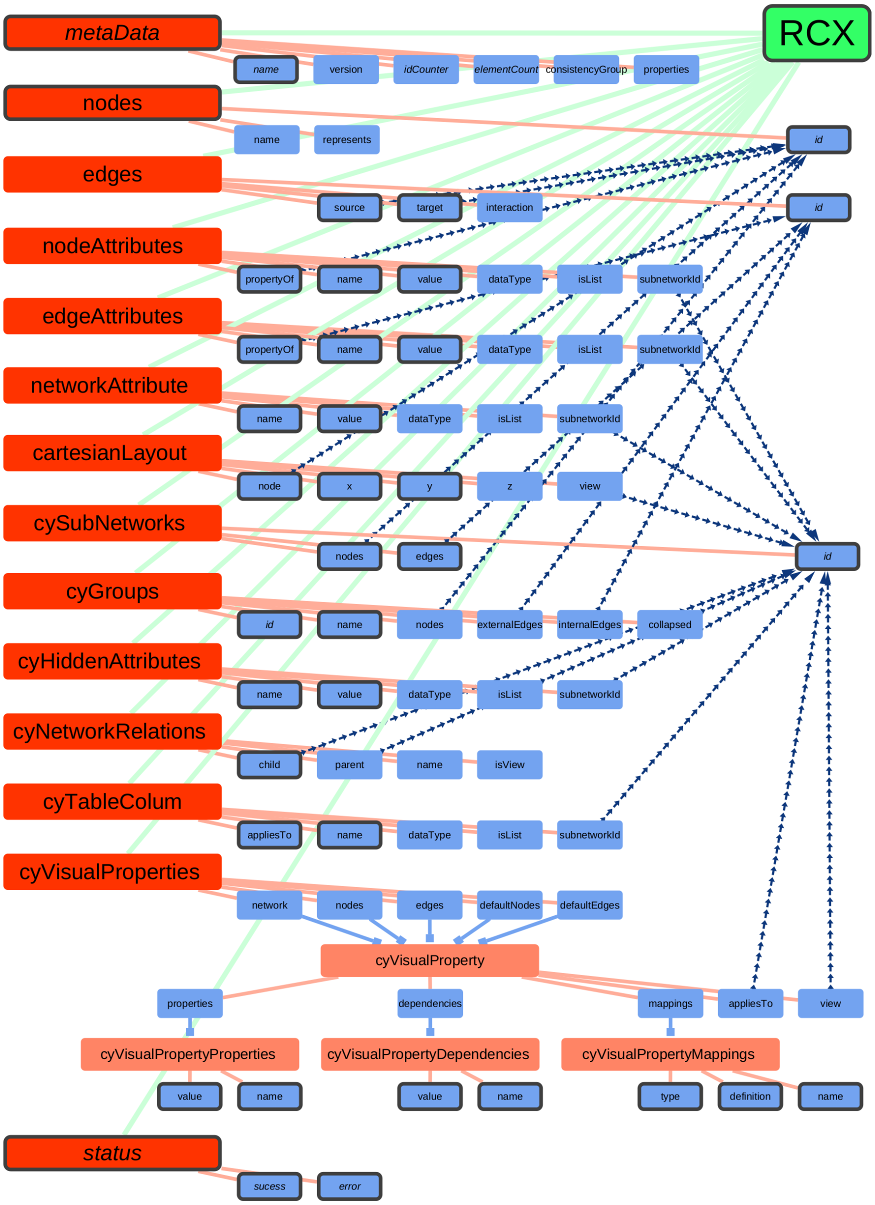


RCX data structure

Doing this manually might be laborious, but it is a good exercise to understand how visual properties and mappings work in detail.

An easier method would be to the current network as CX file, load it in Cytoscape or the online tool NDExEdit (https://frankkramer-lab.github.io/NDExEdit) to create the desired layout.

Afterwards the created layout could be also re-used in other networks that have the same attributes defined:

```
newRcx <- updateCyVisualProperties(newRCX, rcx$cyVisualProperties)
```

### 8 Meta-data

The meta-data of the network is created and updated autimatically. However, we can also update the meta-data manually, and set its `version`, `consisentcyGroup` and `properties`:

```
rcx <- updateMetaData(
  rcx,
  version = c(
    nodes="1.1",
    edges="1.1",
    nodeAttributes="1.1",
    networkAttributes="1.1",
    cartesianLayout="1.1",
    cyVisualProperties="1.1",
    cyTableColumn="1.1"
  ),
  consistencyGroup = c(
    nodes=2,
    edges=2,
    nodeAttributes=2,
    networkAttributes=2,
    cartesianLayout=2,
    cyVisualProperties=2,
    cyTableColumn=2
  ))

rcx$metaData
```

```
## Meta-data:
##                  name version idCounter elementCount consistencyGroup
## 1               nodes     1.1        95           96                2
## 2               edges     1.1       118          119                2
## 3      nodeAttributes     1.1        NA          146                2
## 4   networkAttributes     1.1        NA            3                2
## 5     cartesianLayout     1.1        NA           96                2
## 6 cyVisualProperties     1.1        NA            3                2
## 7       cyTableColumn     1.1        NA            9                2
```

### 9 Create RCX at once

In the above example, we created the RCX network nearly in the beginning and updated it regularly in the course. Instead, we also could simply create the single aspects one by one, and then create an RCX network with all aspects at once:

```
rcx <- createRCX(
  nodes = nodes,
  edges = edges,
  nodeAttributes = nodeAttributes,
  networkAttributes = networkAttributes,
  cartesianLayout = cartesianLayout,
  cyVisualProperties = cyVisualProperties,
  cyTableColumn = cyTableColumn
)
```

### 10 Save the network

After spending this much effort on creating a network, we also want to save it for further use. If we only intend to use it within R, saving it simply as RDS would be sufficient:

```
saveRDS(rcx, "path/to/some-file.rds")
```

For further use of the networks even outside of R, RCX networks can be saved in a similar manner as CX files:

```
writeCX(rcx, "path/to/some-file.cx")
```

Those files can then be used in other platforms like NDEx, Cytoscape or NDExEdit.

In fact, the create network was uploaded to NDEx and can be accessed at https://www.ndexbio.org/viewer/networks/ebdda4da-2ca5-11ec-b3be-0ac135e8bacf

There it is displayed along with its network information and description:

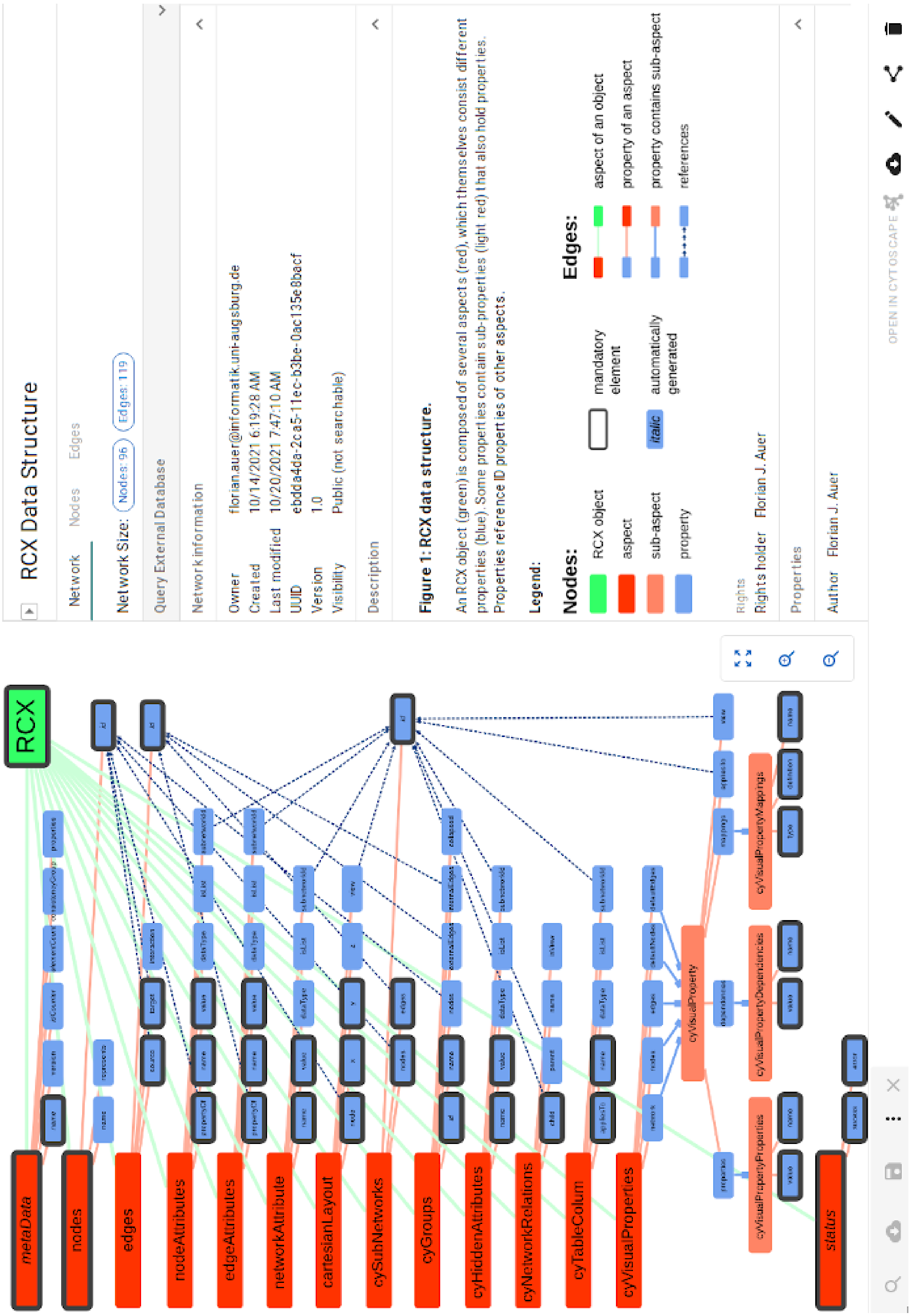


RCX data structure network displayed on NDEx

### 11 Session info

```
sessionInfo()
```

```
## R version 4.0.3 (2020-10-10)
## Platform: x86_64-pc-linux-gnu (64-bit)
## Running under: Ubuntu 20.04.3 LTS
##
## Matrix products: default
## BLAS:   /usr/lib/x86_64-linux-gnu/blas/libblas.so.3.9.0
## LAPACK: /usr/lib/x86_64-linux-gnu/lapack/liblapack.so.3.9.0
##
## locale:
##  [1] LC_CTYPE=en_US.UTF-8       LC_NUMERIC=C
##  [3] LC_TIME=de_DE.UTF-8        LC_COLLATE=en_US.UTF-8
##  [5] LC_MONETARY=de_DE.UTF-8    LC_MESSAGES=en_US.UTF-8
##  [7] LC_PAPER=de_DE.UTF-8       LC_NAME=C
##  [9] LC_ADDRESS=C               LC_TELEPHONE=C
## [11] LC_MEASUREMENT=de_DE.UTF-8 LC_IDENTIFICATION=C
##
## attached base packages:
## [1] stats     graphics  grDevices utils     datasets  methods   base
##
## other attached packages:
## [1] RCX_0.99.0       knitr_1.33       BiocStyle_2.18.1
##
## loaded via a namespace (and not attached):
##  [1] Rcpp_1.0.7          bookdown_0.22       digest_0.6.27
##  [4] plyr_1.8.6          formatR_1.11        magrittr_2.0.1
##  [7] evaluate_0.14       rlang_0.4.11        stringi_1.6.2
## [10] rmarkdown_2.9       tools_4.0.3         stringr_1.4.0
## [13] xfun_0.24           yaml_2.2.1          compiler_4.0.3
## [16] base64enc_0.1-3     BiocManager_1.30.16 htmltools_0.5.1.1
```
