## Supplementary material for "RCX – an R package adapting the Cytoscape Exchange format for biological networks": Extending the RCX Data Model

### 03. Extending the RCX Data Model

***Florian J. Auer***[1*]

[1]Augsburg of University

[*]

**2021-10-26**

### Contents

1 A custom aspect
2 Create the custom aspect in R
3 Update the aspect
    3.1 Providing update methods
    3.2 Update meta-data for the aspect
    3.3 Aspect references
    3.4 Convenience methods
4 Validation of the aspect
5 Conversion to and from CX
    5.1 Convert to CX
    5.2 Conversion from CX/JSON
6 Session info

The CX format is supposed to be flexible, so that custom aspects can be defined by the user. However, the functions provided by the RCX package cannot cover those extensions. To work with those aspects in R, it is necessary to implement functions extending the RCX to support custom aspects. In the following, we will explore how custom aspects can be implemented and integrated in the RCX model.

### 1 A custom aspect

For demonstration purposes we here define our own custom aspect for keeping the network provenance. While similar, this is not the deprecated `provenanceHistory` aspect from previous CX versions! In this example, the JSON structure should look like this:

```
{
  "networkProvenance": [
    {
      "@id": 1,
      "time": 1445437740,
      "action": "created",
      "nodes": [1, 2, 3, 4, 5, 6],
      "source": "https://www.ndexbio.org/viewer/networks/66a902f5-2022-11e9-bb6a-0ac135e8bac
    },
    {
      "@id": 2,
      "time": 1445437770,
      "action": "merged",
      "nodes": [7, 8, 9, 10],
      "source": "https://www.ncbi.nlm.nih.gov/geo/query/acc.cgi?acc=GSE33075"
    },
    {
      "@id": 3,
      "time": 1445437799,
      "action": "filtered",
      "nodes": [],
      "comment": "Some manual filtering was performed"
    }
  ]
}
```

The network provenance structure consists of following properties:

- `@id` : **(required)** and unique ID used in this aspect
- `time` : **(required)** timestamp in seconds since JAN 01 1970. (UTC)
- `action` : **(required)** what was done
- `nodes` : **(required)** list of node IDs, can be empty
- `source` : **(optional)** where new data came from in this step
- `comment` : **(optional)** some commentary on the performed action

It is quite a simple aspect, but it will show the single steps needed to adapt this aspect to the RCX model.

### 2 Create the custom aspect in R

First, load the RCX library:

```
library(RCX)
```

Following the naming convention of the package, we define a simple function to create the aspect in R:

```
createNetworkProvenance <- function(
  id = NULL,
  time,
  action,
  nodes,
  source = NULL,
  comment = NULL){

  ## generate id if not provided
  if(is.null(id)){
    id = 0:(length(name) -1)
  }

  ## create aspect with default values
  res = data.frame(
    id = id,
    time = time,
    action = action,
    nodes = NA,
    source = NA,
    comment = NA,
    stringsAsFactors=FALSE, check.names=FALSE
  )

  ## add nodes
  if(!is.list(nodes)) nodes <- list(nodes)
  res$nodes = nodes


  ## add source if provided
  if(!is.null(source)){
    res$source <- source
  }

  ## add comment if provided
  if(!is.null(comment)){
    res$comment <- comment
  }

  ## add a class name
  class(res) <- append("NetworkProvenanceAspect", class(res))
  return(res)
}
```

Since this is only for demonstration purposes no checks or validations of the data is included. In practice, all data should be checked to avoid mistakes. Also for all following functions validation of the data has been omitted.

Now that we can create objects of our own aspect, let’s do it:

```
networkProvenance <- createNetworkProvenance(
  id = c(1,2,3),
  time = c(1445437740, 1445437770, 1445437799),
  action = c("created", "merged", "filtered"),
  nodes = list(
    c(1, 2, 3, 4, 5, 6),
    c(7, 8, 9, 10),
    c()
  ),
  source = c(
    "https://www.ndexbio.org/viewer/networks/66a902f5-2022-11e9-bb6a-0ac135e8bacf",
    "https://www.ncbi.nlm.nih.gov/geo/query/acc.cgi?acc=GSE33075",
    NA
  ),
  comment = c(NA, NA, "Some manual filtering was performed")
)

networkProvenance
```

```
##   id       time   action             nodes
## 1  1 1445437740  created 1, 2, 3, 4, 5, 6
## 2  2 1445437770   merged      7, 8, 9, 10
## 3  3 1445437799 filtered             NULL
##                                                                       source
## 1 https://www.ndexbio.org/viewer/networks/66a902f5-2022-11e9-bb6a-0ac135e8bacf
## 2                  https://www.ncbi.nlm.nih.gov/geo/query/acc.cgi?acc=GSE33075
## 3                                                                         <NA>
##                               comment
## 1                                <NA>
## 2                                <NA>
## 3 Some manual filtering was performed
```

### 3 Update the aspect

#### 3.1 Providing update methods

For updating the aspect, as well as the RCX object with this aspect, the RCX package uses method dispatch and follows a convention for naming the functions `update<accession name>`. For example for `rcx$nodes` the update function **must** be named `updateNodes`! Firstly, we create for `networkProvenance` a generic function following this convention:

```
updateNetworkProvenance <- function(x, aspect) {
    UseMethod("updateNetworkProvenance", x)
}
```

The first argument must **always** be either an RCX object or a network provenance aspect, the second a network provenance aspect.

Now let's add a method, that merges two network provenance aspects:

```
updateNetworkProvenance.NetworkProvenanceAspect <- function(x, aspect) {
    res <- plyr::rbind.fill(x, aspect)
```

```
    if (!"NetworkProvenanceAspect" %in% class(res)) {
        class(res) <- append("NetworkProvenanceAspect", class(res))
    }
    return(res)
}
```

To test this method, we split our previous example into two parts and merge them. If everything works, we should get the same aspect object as we got from `createNetworkProvenance`.

```
## Split the original example
## Create first part
np1 <- createNetworkProvenance(
  id = c(1,2),
  time = c(1445437740, 1445437770),
  action = c("created", "merged"),
  nodes = list(
    c(1, 2, 3, 4, 5, 6),
    c(7, 8, 9, 10)
  ),
  source = c(
    "https://www.ndexbio.org/viewer/networks/66a902f5-2022-11e9-bb6a-0ac135e8bacf",
    "https://www.ncbi.nlm.nih.gov/geo/query/acc.cgi?acc=GSE33075"
  )
)

## Create second part
np2 <- createNetworkProvenance(
  id = 3,
  time = 1445437799,
  action = "filtered",
  nodes = c(),
  comment = "Some manual filtering was performed"
)

## Merge the parts
networkProvenance <- updateNetworkProvenance(np1, np2)

networkProvenance
```

```
##   id       time   action            nodes
## 1  1 1445437740  created 1, 2, 3, 4, 5, 6
## 2  2 1445437770   merged      7, 8, 9, 10
## 3  3 1445437799 filtered             NULL
##                                                                          source
## 1 https://www.ndexbio.org/viewer/networks/66a902f5-2022-11e9-bb6a-0ac135e8bacf
## 2                  https://www.ncbi.nlm.nih.gov/geo/query/acc.cgi?acc=GSE33075
## 3                                                                          <NA>
##                                 comment
## 1                                  <NA>
## 2                                  <NA>
## 3 Some manual filtering was performed
```

Now that we have the method for two network provenance aspects, we can create another method to merge an RCX object with a network provenance aspects. For this, we simply can use the previous method in this one to merge the two network provenance aspects:

```
updateNetworkProvenance.RCX <- function(x, aspect) {
    rcxAspect <- x$networkProvenance
    if (!is.null(rcxAspect)) {
```

```
        aspect <- updateNetworkProvenance(rcxAspect, aspect)
    }

    x$networkProvenance <- aspect

    x <- updateMetaData(x)

    return(x)
}
```

So if we now update the RCX object with the network provenance parts one by one, we should end up with a combine one in the RCX.

```
## Prepare an RCX object
rcx <- createRCX(
    nodes = createNodes(name = LETTERS[1:10]),
    edges = createEdges(
        source = c(1, 2),
        target = c(2, 3)
    )
)

## Add the first part of network provenance
rcx <- updateNetworkProvenance(rcx, np1)

## Add the second part
rcx <- updateNetworkProvenance(rcx, np2)

rcx
```

```
## [[metaData]] = Meta-data:
##    name version idCounter elementCount consistencyGroup
## 1 nodes     1.0         9           10                1
## 2 edges     1.0         1            2                1
##
## [[nodes]] = Nodes:
##    id name
## 1   0    A
## 2   1    B
## 3   2    C
## 4   3    D
## 5   4    E
## 6   5    F
## 7   6    G
## 8   7    H
## 9   8    I
## 10  9    J
##
## [[edges]] = Edges:
##   id source target
## 1  0      1      2
## 2  1      2      3
##
## [[networkProvenance]] =   id       time   action            nodes
## 1  1 1445437740  created 1, 2, 3, 4, 5, 6
## 2  2 1445437770   merged      7, 8, 9, 10
## 3  3 1445437799 filtered             NULL
##                                                                    source
## 1 https://www.ndexbio.org/viewer/networks/66a902f5-2022-11e9-bb6a-0ac135e8bacf
## 2                  https://www.ncbi.nlm.nih.gov/geo/query/acc.cgi?acc=GSE33075
```

```
## 3                                                                              <NA>
##                                comment
## 1                                 <NA>
## 2                                 <NA>
## 3 Some manual filtering was performed
```

As we see, this works as well, but there is one problem: The meta-data does not contain the aspect yet.

#### 3.2 Update meta-data for the aspect

To include it, we have to provide information about the relation between accession name (`networkProvenance`) and the class of the aspect (`NetworkProvenanceAspect`). This information is kept in the `aspectClasses` vector. To update this vector with a new aspect, we can use the `updateAspectClasses` function:

```
aspectClasses
```

```
##                       rcx                    metaData
##                     "RCX"            "MetaDataAspect"
##                     nodes                       edges
##             "NodesAspect"               "EdgesAspect"
##            nodeAttributes              edgeAttributes
##     "NodeAttributesAspect"     "EdgeAttributesAspect"
##         networkAttributes             cartesianLayout
##  "NetworkAttributesAspect"    "CartesianLayoutAspect"
##                  cyGroups          cyVisualProperties
##            "CyGroupsAspect" "CyVisualPropertiesAspect"
##        cyHiddenAttributes          cyNetworkRelations
## "CyHiddenAttributesAspect" "CyNetworkRelationsAspect"
##              cySubNetworks               cyTableColumn
##      "CySubNetworksAspect"      "CyTableColumnAspect"
```

```
aspectClasses <- updateAspectClasses(
  aspectClasses,
  c(networkProvenance="NetworkProvenanceAspect")
)
aspectClasses
```

```
##                       rcx                    metaData
##                     "RCX"            "MetaDataAspect"
##                     nodes                       edges
##             "NodesAspect"               "EdgesAspect"
##            nodeAttributes              edgeAttributes
##     "NodeAttributesAspect"     "EdgeAttributesAspect"
##         networkAttributes             cartesianLayout
##  "NetworkAttributesAspect"    "CartesianLayoutAspect"
##                  cyGroups          cyVisualProperties
##            "CyGroupsAspect" "CyVisualPropertiesAspect"
##        cyHiddenAttributes          cyNetworkRelations
## "CyHiddenAttributesAspect" "CyNetworkRelationsAspect"
##              cySubNetworks               cyTableColumn
##      "CySubNetworksAspect"      "CyTableColumnAspect"
##         networkProvenance
##  "NetworkProvenanceAspect"
```

The meta-data can be updated manually using the `updateMetaData` function. However, to get the updated `aspectClass` to work, we have to provide it to the update function:

```
rcx <- updateMetaData(rcx)
rcx$metaData
```

```
## Meta-data:
##    name version idCounter elementCount consistencyGroup
## 1 nodes     1.0         9           10                1
## 2 edges     1.0         1            2                1
```

```
rcx <- updateMetaData(rcx, aspectClasses = aspectClasses)
rcx$metaData
```

```
## Meta-data:
##                name version idCounter elementCount consistencyGroup
## 1             nodes     1.0         9           10                1
## 2             edges     1.0         1            2                1
## 3 networkProvenance     1.0        NA            3                1
```

Now the meta data is updated with our custom aspect. To automatically update the meta-data when the provenance history is updated, we can add it to our update method for the RCX object:

```
updateNetworkProvenance.RCX <- function(x, aspect) {
    rcxAspect <- x$networkProvenance
    if (!is.null(rcxAspect)) {
        aspect <- updateNetworkProvenance(rcxAspect, aspect)
    }

    x$networkProvenance <- aspect

    x <- updateMetaData(x, aspectClasses = aspectClasses)

    return(x)
}
```

The `idCounter` of the meta-data is not updated yet, although this aspect contains an ID. To enable this, we have to implement two methods:

```
hasIds.NetworkProvenanceAspect <- function(aspect) {
    return(TRUE)
}

idProperty.NetworkProvenanceAspect <- function(aspect) {
    return("id")
}
```

The first one simply tells that the network provenance aspect has IDs, the second one returns the property, that holds the id. When we now update the meta-data, the `idCounter` is updated as well.

```
rcx <- updateMetaData(rcx, aspectClasses = aspectClasses)
rcx$metaData
```

```
## Meta-data:
##                name version idCounter elementCount consistencyGroup
## 1             nodes     1.0         9           10                1
## 2             edges     1.0         1            2                1
## 3 networkProvenance     1.0         3            3                1
```

Alternatively, we could specify the timestamp as id, and subsequently omit the `id` column in general.

```
idProperty.NetworkProvenanceAspect <- function(aspect) {
    return("time")
}

rcx <- updateMetaData(rcx, aspectClasses = aspectClasses)
rcx$metaData
```

```
## Meta-data:
##                 name version  idCounter elementCount consistencyGroup
## 1              nodes     1.0          9           10                1
## 2              edges     1.0          1            2                1
## 3 networkProvenance     1.0 1445437799            3                1
```

This might be useful in some cases, but for this example we stick to `id` as the dedicated column providing the IDs in our aspect.

```
idProperty.NetworkProvenanceAspect <- function(aspect) {
    return("id")
}
```

### 3.3 Aspect references

Additionally, we can provide a method to determine to which other aspects our custom aspect is referring, and by which other aspects are referred. In our case this would be the nodes aspect with its IDs. This later could be used in the validation.

```
refersTo.NetworkProvenanceAspect <- function(aspect) {
    nodes <- aspectClasses["nodes"]
    names(nodes) <- NULL
    result <- c(nodes = nodes)
    return(result)
}

refersTo(rcx$edges)
```

```
##        source        target
## "NodesAspect" "NodesAspect"
```

```
refersTo(rcx$networkProvenance)
```

```
##         nodes
## "NodesAspect"
```

```
referredBy(rcx)
```

```
## $NodesAspect
## [1] "EdgesAspect"
```

```
referredBy(rcx, aspectClasses)
```

```
## $NodesAspect
## [1] "EdgesAspect"             "NetworkProvenanceAspect"
```

#### 3.4 Convenience methods

To be consistent with the other aspects of the RCX package, we can provide a custom print method, that adds the aspect name before printing:

```
print.NetworkProvenanceAspect <- function(x, ...) {
    cat("Network provenance:\n")
    class(x) <- class(x)[!class(x) %in%
        "NetworkProvenanceAspect"]
    print(x, ...)
}
```

There are also further function, like `summary` and `countElements` that could be adjusted, but for this example they are not needed and work analogously.

```
summary.NetworkProvenanceAspect <- function(object, ...) {
    ...
}
countElements.NetworkProvenanceAspect <- function(x) {
    ...
}
```

### 4 Validation of the aspect

It is always a good idea to provide functions to validate the correctness of the data. Therefore, we implement the `validate` method for our aspect. What to evaluate in this method is up to the user, but the more checks and information provided, the more it helps other users to track down errors.

```
validate.NetworkProvenanceAspect = function(x, verbose=TRUE){
    if(verbose) cat("Checking Network Provenance Aspect:\n")

    test = all(! is.na(x$id))
    if(verbose) cat(paste0("- Column (id) doesn't contain any NA values...",
                           ifelse(test, "OK", "FAIL"),
                           "\n"))
    pass = test

    test = length(x$id) == length(unique(x$id))
    if(verbose) cat(paste0("- Column (id) contains only unique values...",
                           ifelse(test, "OK", "FAIL"),
                           "\n"))
    pass = pass & test


    if(verbose) cat(paste0(">> Network Provenance Aspect: ",
                           ifelse(test, "OK", "FAIL"),
                           "\n"))
    invisible(pass)
}
```

```r
validate(rcx, verbose = TRUE)
```

```
## Checking Nodes Aspect:
## - Is object of class "NodesAspect"...OK
## - All required columns present (id)...OK
## - Column (id) doesn't contain any NA values...OK
## - At least one node present...OK
## - Column (id) contains only unique values...OK
## - Column (id) only contains numeric values...OK
## - Column (id) only contains positive (>=0) values...OK
## - No merge artefacts present (i.e. column with old ids: oldId)...OK
## - Only allowed columns present (id, name, represents)...OK
## >> Nodes Aspect: OK
##
## Checking Edges Aspect:
## - Is object of class "EdgesAspect"...OK
## - All required columns present (id, source, target)...OK
## - Column (id) doesn't contain any NA values...OK
## - Column (id) contains only unique values...OK
## - Column (id) only contains numeric values...OK
## - Column (id) only contains positive (>=0) values...OK
## - Column (source) doesn't contain any NA values...OK
## - Column (source) only contains numeric values...OK
## - Column (source) only contains positive (>=0) values...OK
## - Column (target) doesn't contain any NA values...OK
## - Column (target) only contains numeric values...OK
## - Column (target) only contains positive (>=0) values...OK
## - No merge artefacts present (i.e. column with old ids: oldId)...OK
## - Only allowed columns present (id, source, target, name, interaction)...OK
## >> Edges Aspect: OK
##
## Checking Custom Aspects:
## Checking Network Provenance Aspect:
## - Column (id) doesn't contain any NA values...OK
## - Column (id) contains only unique values...OK
## >> Network Provenance Aspect: OK
## >> Cytoscape Table Column Aspect: OK
##
## Checking RCX:
## - Is object of class "RCX"...OK
## - nodes aspect is present...OK
## - Validate nodes aspect...OK
## - Validate edges aspect...OK
##   - Reference aspect (nodes) present and correct...OK
##   - All id references exist (EdgesAspect$source ids in NodesAspect$id)...OK
##   - All id references exist (EdgesAspect$target ids in NodesAspect$id)...OK
## >> RCX: OK
```

With this method, not only our custom aspect is evaluated solely, the method is also called when the whole RCX object is evaluated. Therefore, if validation for the aspect fails, also the validation of the RCX object fails.

### 5 Conversion to and from CX

#### 5.1 Convert to CX

Since we have our aspect data model already created in R, let's start with the conversion to CX in JSON format. To allow the aspect to be processed, we have to provide a method to take over this part. In the RCX package, the aspect class specific methods of `rcxToJson` convert the aspect to a JSON object.

```
rcxToJson.NetworkProvenanceAspect <- function(aspect, verbose = FALSE, ...) {
    if (verbose)
        cat("Convert network provenance to JSON...")

    ## rename id to @id
    colnames(aspect) <- gsub("id", "\\@id", colnames(aspect))

    ## convert to json
    json <- jsonlite::toJSON(aspect, pretty = TRUE)

    ## empty nodes are converted to 'nodes': {}, the simplest fix is just
    ## replacing it
    json <- gsub("\"nodes\": \\{\\},", "\"nodes\": \\[\\],", json)

    ## add the aspect name
    json <- paste0("{\"networkProvenance\":", json, "}")

    if (verbose)
        cat("done!\n")
    return(json)
}

cat(rcxToJson(rcx$networkProvenance))
```

```
## {"networkProvenance":[
##   {
##     "@id": 1,
##     "time": 1445437740,
##     "action": "created",
##     "nodes": [1, 2, 3, 4, 5, 6],
##     "source": "https://www.ndexbio.org/viewer/networks/66a902f5-2022-11e9-bb6a-0ac135e8ba[illegible]
##   },
##   {
##     "@id": 2,
##     "time": 1445437770,
##     "action": "merged",
##     "nodes": [7, 8, 9, 10],
##     "source": "https://www.ncbi.nlm.nih.gov/geo/query/acc.cgi?acc=GSE33075"
##   },
##   {
##     "@id": 3,
##     "time": 1445437799,
##     "action": "filtered",
##     "nodes": [],
##     "comment": "Some manual filtering was performed"
##   }
## ]}
```

The `rcxToJson` functions are used in the `toCX` and `writeCX` functions for converting and saving the RCX. So let's write the RCX to a file we can use later for reading.

```
tempCX <- tempfile(fileext = ".cx")
writeCX(rcx, tempCX)
```

```
## [1] "/tmp/Rtmpy2UPQs/filef953d4901d3.cx"
```

#### 5.2 Conversion from CX/JSON

Similarly to `rcxToJson` there exists a method for doing the reverse. The `readCX` function combines several steps, that can be performed individually:

1. `readJSON` which simply read the JSON from file
2. `parseJSON` which parsed the JSON text to JSON data (list of lists)
3. `processCX` takes the JSON data and calls for each aspect the `jsonToRCX`

The correct aspect can be accessed in the `jsonData` list, which then has to be processed and a created object of the aspect returned.

```
jsonToRCX.networkProvenance <- function(jsonData, verbose) {
    if (verbose)
        cat("Parsing network provenance...")
    data <- jsonData$networkProvenance

    ids <- sapply(
        data, function(d) {
            d$`@id`
        }
    )
    time <- sapply(
        data, function(d) {
            d$time
        }
    )
    action <- sapply(
        data, function(d) {
            d$action
        }
    )
    nodes <- sapply(
        data, function(d) {
            d$nodes
        }
    )
    source <- sapply(
        data, function(d) {
            d$source
        }
    )
    comment <- sapply(
        data, function(d) {
            d$comment
        }
    )

    if (verbose)
        cat("create aspect...")
    result <- createNetworkProvenance(
        id = ids, time = time, action = action, nodes = nodes, source = source,
        comment = comment
    )

    if (verbose)
```

```
        cat("done!\n")
    return(result)
}
rcxParsed <- readCX(tempCX, aspectClasses = aspectClasses)
```

However, the meta-data has to be updated again manually with the `aspectClasses` available:

```
rcxParsed <- updateMetaData(rcxParsed, aspectClasses = aspectClasses)
rcxParsed$metaData
```

```
## Meta-data:
##                name version idCounter elementCount consistencyGroup
## 1             nodes     1.0         9           10                1
## 2             edges     1.0         1            2                1
## 3 networkProvenance     1.0         3            3                1
```

So we have successfully converted our custom aspect between CX and RCX!

### 6 Session info

```
sessionInfo()
```

```
## R version 4.0.3 (2020-10-10)
## Platform: x86_64-pc-linux-gnu (64-bit)
## Running under: Ubuntu 20.04.3 LTS
##
## Matrix products: default
## BLAS:   /usr/lib/x86_64-linux-gnu/blas/libblas.so.3.9.0
## LAPACK: /usr/lib/x86_64-linux-gnu/lapack/liblapack.so.3.9.0
##
## locale:
##  [1] LC_CTYPE=en_US.UTF-8       LC_NUMERIC=C
##  [3] LC_TIME=de_DE.UTF-8        LC_COLLATE=en_US.UTF-8
##  [5] LC_MONETARY=de_DE.UTF-8    LC_MESSAGES=en_US.UTF-8
##  [7] LC_PAPER=de_DE.UTF-8       LC_NAME=C
##  [9] LC_ADDRESS=C               LC_TELEPHONE=C
## [11] LC_MEASUREMENT=de_DE.UTF-8 LC_IDENTIFICATION=C
##
## attached base packages:
## [1] stats     graphics  grDevices utils     datasets  methods   base
##
## other attached packages:
## [1] RCX_0.99.0       knitr_1.36       BiocStyle_2.18.1
##
## loaded via a namespace (and not attached):
##  [1] Rcpp_1.0.7          bookdown_0.24       digest_0.6.28
##  [4] plyr_1.8.6          jsonlite_1.7.2      formatR_1.11
##  [7] magrittr_2.0.1      evaluate_0.14       rlang_0.4.12
## [10] stringi_1.7.5       jquerylib_0.1.4     rmarkdown_2.11
## [13] tools_4.0.3         stringr_1.4.0       xfun_0.27
## [16] yaml_2.2.1          fastmap_1.1.0       compiler_4.0.3
## [19] BiocManager_1.30.16 htmltools_0.5.2
```
