## Supplementary material for "RCX – an R package adapting the Cytoscape Exchange format for biological networks": The RCX and CX Data Model

### Appendix: The RCX and CX Data Model

***Florian J. Auer***[1*]

[1]University Augsburg

[*]

**2021-10-26**

### Contents

1 CX data structure
- 1.1 Aspect dependencies

2 Data types

3 NDEx conventions
- 3.1 Handling of Identifiers
- 3.2 Citations

4 Meta aspects
- 4.1 metaData
- 4.2 status

5 Core aspects
- 5.1 nodes
- 5.2 edges
- 5.3 nodeAttributes
  - 5.3.1 NDEx conventions
- 5.4 edgeAttributes
- 5.5 networkAttributes
  - 5.5.1 NDEx conventions
- 5.6 cartesianLayout

6 Cytoscape aspects
- 6.1 cyGroups
- 6.2 CyVisualProperties
  - 6.2.1 CyVisualProperties (CX)
  - 6.2.2 CyVisualProperties (RCX)
- 6.3 cyHiddenAttributes
- 6.4 cyNetworkRelations
- 6.5 cySubNetworks
- 6.6 cyTableColum

7 Deprecated aspects

8 Session info

### 1 CX data structure

The CX data structure can be divided into three main classes:

- Meta aspects: necessary for data transmission
  - metaData
  - status
- Core aspects: essential for defining a network
  - nodes
  - edges
  - nodeAttributes
  - edgeAttributes
  - networkAttributes
  - cartesianLayout
- Cytoscape aspects: aspects needed to define the visual layout of a network in Cytoscape
  - cyGroups
  - cyVisualProperties
  - cyHiddenAttributes
  - cyNetworkRelations
  - cySubnetworks
  - cyTableColumn

#### 1.1 Aspect dependencies

Aspects with IDs:

- nodes
- edges
- cyGroups
- cySubnetworks

Aspects, that reference IDs:

- edges reference nodes by "source" and "target"
- nodeAttributes reference nodes by "propertyOf"
- edgeAttributes reference edges by "propertyOf"
- cartesianLayout references cySubnetworks by "view"
- cyGroups reference nodes by "nodes" and edges by "internalEdges" and "externalEdges"
- sub-aspect cyVisualProperty of cyVisualProperties references cySubnetworks by "appliesTo" and "view"
- cySubnetworks reference nodes by "nodes" and edges by "edges"

### 2 Data types

CX is a JSON based format, that means that JavaScript data types are used. Those will be mapped to and from R data types as follows:

| data type | example | R data type |
|---|---|---|
| boolean | “true” | logical |
| double | “2.3” | numeric |
| integer | “23” | integer |
| long | “123456” | integer |
| string | “blue” | character |
| list_of_boolean | [“true”,“false”] | list(logical) |
| list_of_double | [“2.3”,“4.5”] | list(numeric) |
| list_of_integer | [“23”,“45”] | list(integer) |
| list_of_long | [“23”,“45”] | list(integer) |
| list_of_string | [“Marco”,“Polo”] | list(character) |

### 3 NDEx conventions

#### 3.1 Handling of Identifiers

All identifiers should be either:

- <prefix>:<id> format
- a string without " : "
- a URI

#### 3.2 Citations

It is recomended to use the attribute “citation” on edges or nodes to store citations to specify literature references or other sources of information that are relevant to the network. Citations are primarily described by five dublin core terms. The “dc” prefix is implicitly interpreted as referencing dublin core in the context of the citations aspect.

The five primary dublin core terms defined for citations are:

- dc:title
  - http://dublincore.org/documents/2012/06/14/dcmi-terms/?v=terms#terms-title
- dc:contributor
  - http://dublincore.org/documents/2012/06/14/dcmi-terms/?v=terms#terms-contributor
- dc:identifier
  - http://dublincore.org/documents/2012/06/14/dcmi-terms/?v=terms#terms-identifier
  - Ideally citations make use of the “identifier” key to link to their source. This can be a URI, but might also use the “source:id” format, where source is usually “pmid” or “doi”.
- dc:type
  - http://dublincore.org/documents/2012/06/14/dcmi-terms/?v=terms#terms-type
- dc:description
  - http://dublincore.org/documents/2012/06/14/dcmi-terms/?v=elements#description
  - This should be a description of the resource, and it is preferable that information that can be expressed in more specific attributes should not be in this attribute. The “description” attribute could contain the title, authors, and/or journal reference, but in that case it would not be expected to be machine parsable.

### 4 Meta aspects

#### 4.1 metaData

As a transfer format,the owner (sender) is considered to have a complete picture of the network including all metadata and aspects associated with the network. When a sender has the choice of sending an element as either pre- or post-metadata. In RCX both are combined when converting a CX file. The content a the metaData aspect is specified as:

| RCX property | RCX specifics | CX property | CX options | description |
|---|---|---|---|---|
| name | | name | Required in pre- and post-metadata | The name of the aspect |
| version | default:“1.0” | version | Required in pre-metadata | version of this aspect schema (“1.1.0”, “2.0.0”, etc.) |
| idCounter | auto-generated | idCounter | Required if the aspect exports IDs | Integer (All Element IDs are integers; see node id, edge id, cytoscape groups) |
| elementCount | auto-generated | elementCount | Optional | number (integer) of elements in this aspect |
| consistencyGroup | default:1 | consistencyGroup | Optional | An integer identifier shared by aspects to indicate that they are mutually consistent |
| properties | | properties | Optional | An aspect-defined property list |
| - | *not supported* | checksum | Optional *(Deprecated)* | *NDEx CX implementation does not support this attribute. This attribute is ignored in NDEx* |

#### 4.2 status

A complete CX stream ends with a status aspect object. This aspect tells the recipient if the CX is successfully generated by the source.

**Note:** This aspect is auto-generated in RCX and cannot be changed!

| property | options | values | description |
|---|---|---|---|
| success | Required | “true” or “false” | Was the CX successfully generated by the source |
| error | Required | string or "" | error message or "" if sucess==“true” |

### 5 Core aspects

#### 5.1 nodes

| RCX property | CX property | CX options | values | description |
|---|---|---|---|---|
| id | @id | Required, Unique | integer | CX internal id, starts at 0, exported to metaData! |
| name | n | Optional | string | node name, eg. “EGFR”, “node 1” |
| represents | r | Optional | string | represents, eg. “HGNC:AKT1” |

**Note:** At least one node has to be present, therefore this aspect is mandatory!

#### 5.2 edges

| RCX property | CX property | CX options | values | description |
|---|---|---|---|---|
| id | @id | Required, Unique | integer | CX internal id, starts at 0, exported to metaData! |
| source | s | Required | integer | source, reference to CX internal node id |
| target | t | Required | integer | target, reference to CX internal node id |
| interaction | i | Optional | string | intercation, eg. “binds” |

#### 5.3 nodeAttributes

| RCX property | RCX specifics | CX property | options | values | description |
|---|---|---|---|---|---|
| propertyOf | | po | Required | integer | property of, reference to CX internal node id |
| name | | n | Required | string | attribute name, eg. “weight”, “name”, “selected” |
| value | | v | Required | string or list of strings | attribute value(s), eg. [“2”, “0.34”, “2.3”], “Node 6”, “true” |

| RCX property | RCX specifics | CX property | options | values | description |
|---|---|---|---|---|---|
| dataType | default:“string” | d | Optional | string | data type, default “string” |
| isList | default:FALSE | d | Optional | string | If set to TRUE, the CX data type will be modified |
| subnetworkId | default:NA | s | Optional | integer | reference to CX internal subnetwork id |

##### 5.3.1 NDEx conventions

| names | description |
|---|---|
| alias | alternative identifiers for the node. Same meaning as BioPAX “aliases” |
| relatedTo | identifiers denoting concepts related to the node. Same meaning as BioPAX “relatedTerms” |

#### 5.4 edgeAttributes

| RCX property | RCX specifics | CX property | options | values | description |
|---|---|---|---|---|---|
| propertyOf | | po | Required | integer | property of, reference to CX internal edge id |
| name | | n | Required | string | attribute name, eg. “weight”, “name”, “selected” |
| value | | v | Required | string or list of strings | attribute value(s), eg. [“2”, “0.34”, “2.3”], “Node 6”, “true” |
| dataType | default:“string” | d | Optional | string | data type, default “string” |
| isList | default:FALSE | d | Optional | boolean | If set to TRUE, the CX data type will be modified |
| subnetworkId | default:NA | s | Optional | integer | reference to CX internal subnetwork id |

#### 5.5 networkAttributes

| RCX property | RCX specifics | CX property | options | values | description |
|---|---|---|---|---|---|
| name | | n | Required | string | attribute name, eg. “dc:title” |
| value | | v | Required | string or list of strings | attribute value(s), eg. "Result of heat |
| dataType | default:“string” | d | Optional | string | data type, default “string” |
| isList | default:FALSE | d | Optional | boolean | If set to TRUE, the CX data type will be modified |
| subnetworkId | default:NA | s | Optional | integer | reference to CX internal subnetwork id |

##### 5.5.1 NDEx conventions

| names | description |
|---|---|
| name | the name of the network |
| description | a description of the network |
| version | NDEx will treat this attribute as the version of the network. Format is not controlled but best practice is to use string conforming to Semantic Versioning. |
| ndex:sourceFormat | used by NDEx to indicate format of an original file imported, can determine semantics as well. The NDEx UI will allow export options based on this value. Applications that alter a network such that it can no longer be exported in the format should remove the value. |

#### 5.6 cartesianLayout

Cartesian layout elements store coordinates of nodes.

| RCX property | CX property | options | values | description |
|---|---|---|---|---|
| node | node | Required | integer | reference to CX internal node id |
| x | x | Required | numeric | x coordinate |
| y | y | Required | numeric | y coordinate |
| z | z | Optional | numeric | z coordinate |
| view | view | Optional | integer | reference to CX internal reference to CX internal subnetwork id of type view |

### 6 Cytoscape aspects

See Cytoscape Styles on how styles are created in Cytoscape. The following aspects are based on how Cytoscape defines the visual representation of networks. For further information see Using Visual Properies in RCy3, which gives an overview of how to handle Visual properties in R automation.

#### 6.1 cyGroups

| RCX property | CX property | options | values | description |
|---|---|---|---|---|
| id | @id | Required, Unique | integer | CX internal id, starts at 0, exported to metaData! |
| n | n | Required | string | name of this group |
| nodes | nodes | Optional | list of integer | the nodes making up the group, reference to CX internal node id |
| externalEdges | external_edges | Optional | list of integer | the external edges making up the group, reference to CX internal edge id |
| internalEdges | internal_edges | Optional | list of integer | the internal edges making up the group, reference to CX internal edge id |
| collapsed | collapsed | Optional | boolean | indicate whether the group is displayed as a single node |

#### 6.2 CyVisualProperties

##### 6.2.1 CyVisualProperties (CX)

| CX property | options | values | description |
|---|---|---|---|
| properties_of | Required | string, dictionary | to indicate the element type these properties belong to, allowed values are: "network", "nodes:default", "edges:default", "nodes", "edges" |
| applies_to | Required | integer | the identifier of the element these properties apply to, reference to CX internal node id or edge id |
| view | Optional | integer | reference to CX internal subnetwork id of type view |
| properties | Optional | named list of strings | pairs of the actual properties, e.g. "NODE_BORDER_STROKE" : "SOLID", "NODE_BORDER_WIDTH" : "1.5", "NODE_FILL_COLOR" : "#FF3399" |
| dependencies | Required | named list of strings | pairs of the dependencies, e.g. "nodeCustomGraphicsSizeSync" : "true", "nodeSizeLocked" : "false" |
| mappings | Optional | named list of named list of strings, dictionary | e.g. "NODE_BORDER_WIDTH" : {"type" : "DISCRETE", "definition" : "COL=required,T=boolean,K=0=true,V=0=10.0"} |

The JSON object model for the cyVisualProperties aspect is not suitable to be proper represented in R data structures, therefore it is split up into the main aspect and several sub-aspects. The R structure looks as follows:

```
CyVisualProperties
├──network = CyVisualProperty
├──nodes = CyVisualProperty
├──edges = CyVisualProperty
├──defaultNodes = CyVisualProperty
└──defaultEdges = CyVisualProperty

CyVisualProperty
├──properties = CyVisualPropertyProperties
│    ├──name
│    └──value
├──dependencies = CyVisualPropertyDependencies
│    ├──name
│    └──value
├──mappings = CyVisualPropertyMappings
│    ├──name
│    ├──type
│    └──definition
├──appliesTo = <reference to subnetwork id>
└──view = <reference to subnetwork id>
```

##### 6.2.2 CyVisualProperties (RCX)

| RCX property | options | values | description |
|---|---|---|---|
| network | Optional | list of CyVisualProperty | represents "property_of"="network" of CyVisualProperties |
| nodes | Optional | list of CyVisualProperty | represents "property_of"="nodes:default" of CyVisualProperties |
| edges | Optional | list of CyVisualProperty | represents "property_of"="edges" of CyVisualProperties |
| defaultNodes | Optional | list of CyVisualProperty | represents "property_of"="nodes:default" of CyVisualProperties |
| defaultEdges | Optional | list of CyVisualProperty | represents "property_of"="edges:default" of CyVisualProperties |

###### 6.2.2.1 CyVisualProperty

| RCX property | RCX specifics | options | values | description |
|---|---|---|---|---|
| properties | default:NA | Optional | CyVisualPropertyProperties | represents "property_of"="network" of CyVisualProperties |
| depencdencies | default:NA | Optional | CyVisualPropertyDependencies | represents "property_of"="nodes:default" of CyVisualProperties |
| mappings | default:NA | Optional | CyVisualPropertyMappings | represents "property_of"="edges" of CyVisualProperties |
| appliesTo | default:NA | Optional | integer | reference to CX internal subnetwork id |
| view | default:NA | Optional | integer | reference to CX internal subnetwork id of type view |

###### 6.2.2.2 CyVisualPropertyProperties

| RCX property | options | values | description |
|---|---|---|---|
| name | Required | string | name of the property, e.g. "NODE_PAINT" or "EDGE_VISIBLE" |
| value | Required | string | value of the property, e.g. "#FF0000" or "true" |

##### 6.2.2.3 CyVisualPropertyDependencies

| RCX property | options | values | description |
| --- | --- | --- | --- |
| name | Required | string | name of the dependency, e.g. “nodeCustomGraphicsSizeSync” or “nodeSizeLocked” |
| value | Required | string | value of the dependency, e.g. “true” or “false” |

##### 6.2.2.4 CyVisualPropertyMappings

The JSON objects or the form `{"NODE_BORDER_WIDTH" : {"type" : "DISCRETE", "definition" : "COL=required,T=boolean,K=0=true,V=0=10.0"}}` are split up as follows:

| RCX property | options | values | description |
| --- | --- | --- | --- |
| name | Required | string | name of the property, e.g. “NODE_BORDER_WIDTH” |
| type | Required | string | value of the property, e.g. “DISCRETE” |
| definition | Required | string | value of the property, e.g. “COL=required,T=boolean,K=0=true,V=0=10.0” |

#### 6.3 cyHiddenAttributes

| RCX property | RCX specifics | CX property | options | values | description |
| --- | --- | --- | --- | --- | --- |
| name | | n | Required | string | name of this hidden attribute |
| value | | v | Required | string or list of strings | attribute value(s), eg. “layoutAlgorithm” |
| dataType | default:“string” | d | Optional | string | data type, default “string” |
| isList | default:FALSE | d | Optional | boolean | If set to TRUE, the CX data type will be modified |
| subnetworkId | default:NA | s | Optional | integer | reference to CX internal subnetwork id |

#### 6.4 cyNetworkRelations

| RCX property | RCX specifics | CX property | options | values | description |
|---|---|---|---|---|---|
| child | | c | Required | integer | child network, reference to CX internal subnetwork id |
| parent | default:NA | p | Optional | integer | parent network, reference to CX internal subnetwork id |
| name | | name | Optional | string | the name of the child network; if missing, default is reader-dependent |
| isView | default:FALSE | r | Optional | boolean | relationship type, default "subnetwork" |

#### 6.5 cySubNetworks

| RCX property | CX property | options | values | description |
|---|---|---|---|---|
| id | @id | Required, Unique | integer | CX internal id, starts at 0, exported to metaData! |
| nodes | nodes | Optional | list of integer or "all" | the nodes making up the group, reference to CX internal node id |
| edges | edges | Optional | list of integer or "all" | the edges making up the group, reference to CX internal edge id |

#### 6.6 cyTableColum

These elements are used to represent Cytoscape table column labels and types. Its main use is to disambiguate empty table columns.

| RCX property | RCX specifics | CX property | options | values | description |
|---|---|---|---|---|---|
| name | | n | Required | string | name of the column; usually the different attributes of nodes, edges and networks |

| RCX property | RCX specifics | CX property | options | values | description |
|---|---|---|---|---|---|
| appliesTo | “nodes”, “edges”, or “networks” | applies_to | Required | string: “node_table”, “edge_table”, or “network_table” | indicates the Cytoscape table this applies to |
| dataType | default:“string” | d | Optional | string | data type, default “string” |
| isList | default:FALSE | d | Optional | boolean | If set to TRUE, the CX data type will be modified |
| subnetworkId | default:NA | s | Optional | integer | reference to CX internal subnetwork id |

**Note:** Cytoscape does not currently support table columns for the root network, but this is option is included here for consistency

### 7 Deprecated aspects

| property | description |
|---|---|
| ndexStatus | NDEx server will no longer generate this aspect from the server side. Network last updated before the v2.4.0 release will still have this aspect. Users can use the “Get Network Summary” API function to get the status and other information of a network. |
| citations | We recommend using the attribute “citation” on edges or nodes to store citations. |
| nodeCitations | We recommend using the attribute “citation” on nodes to store citations. |
| edgeCitations | We recommend using the attribute “citation” on edges to store citations. |
| Supports | We recommend using the attribute “supports” on edges or nodes. |
| nodeSupports | We recommend using the attribute “supports” on nodes. |
| edgeSupports | We recommend using the attribute “supports” on edges. |
| functionTerms | No recommendations for alternative representation. |
| reiefiedEdges | No recommendations for alternative representation. |
| @context | We recommend using the attribute “@context” on networks. The value of this attribute is a serialized string from a JSON dictionary that has a prefix as its key and a URL as its value. @context maps terms to IRIs. Terms are case sensitive. |
| provenanceHistory | The server will just treat this as an opaque aspect. For recording the origin and significant events of this network, we recommend to use network attributes to store them. We recommend using prov:wasGeneratedBy and prov:wasDerivedFrom for the event and the list of contributing network URLs. |

**Note:** Starting from v2.4.0, NDEx server will no longer generate these aspects from the server side. Networks last updated before the v2.4.0 release will still have these aspects.

### 8 Session info

```
sessionInfo()
```

```
## R version 4.0.3 (2020-10-10)
## Platform: x86_64-pc-linux-gnu (64-bit)
## Running under: Ubuntu 20.04.3 LTS
##
## Matrix products: default
## BLAS:   /usr/lib/x86_64-linux-gnu/blas/libblas.so.3.9.0
## LAPACK: /usr/lib/x86_64-linux-gnu/lapack/liblapack.so.3.9.0
##
## locale:
##  [1] LC_CTYPE=en_US.UTF-8       LC_NUMERIC=C
##  [3] LC_TIME=de_DE.UTF-8        LC_COLLATE=en_US.UTF-8
##  [5] LC_MONETARY=de_DE.UTF-8    LC_MESSAGES=en_US.UTF-8
##  [7] LC_PAPER=de_DE.UTF-8       LC_NAME=C
##  [9] LC_ADDRESS=C               LC_TELEPHONE=C
## [11] LC_MEASUREMENT=de_DE.UTF-8 LC_IDENTIFICATION=C
##
## attached base packages:
## [1] stats     graphics  grDevices utils     datasets  methods   base
##
## other attached packages:
## [1] BiocStyle_2.18.1
##
## loaded via a namespace (and not attached):
##  [1] bookdown_0.24       digest_0.6.28       magrittr_2.0.1
##  [4] evaluate_0.14       rlang_0.4.12        stringi_1.7.5
##  [7] jquerylib_0.1.4     rmarkdown_2.11      tools_4.0.3
## [10] stringr_1.4.0       xfun_0.27           yaml_2.2.1
## [13] fastmap_1.1.0       compiler_4.0.3      BiocManager_1.30.16
## [16] htmltools_0.5.2     knitr_1.36
```
