## Supplementary material for "RCX – an R package adapting the Cytoscape Exchange format for biological networks": RCX Reference Manual

### Package ‘RCX’

October 27, 2021

**Type** Package

**Title** R package implementing the Cytoscape Exchange (CX) format

**Version** 0.99.0

**Description** Create, handle, validate, visualize and convert networks in the Cytoscape exchange (CX) format to standard data types and objects.
The package also provides conversion to and from objects of iGraph and graphNEL.
The CX format is also used by the NDEx platform, a online commons for biological networks, and the network visualization software Cytocape.

**License** MIT + file LICENSE

**Depends** R (>= 4.0)

**Imports** jsonlite,
plyr,
igraph

**RoxygenNote** 7.1.2

**Roxygen** list(markdown = TRUE)

**biocViews** Pathways, DataImport, Network

**Suggests** BiocStyle,
testthat,
knitr,
rmarkdown,
base64enc,
graph

**VignetteBuilder** knitr

**Encoding** UTF-8

**URL** `https://github.com/frankkramer-lab/RCX`

**BugReports** `https://github.com/frankkramer-lab/RCX/issues`

#### R topics documented:

**Index** **110**

---

`aspectClasses` *aspectClasses and subAspectClasses*

---

#### Description

`aspectClasses` and `subAspectClasses` contain the accession name and the classes of the corresponding (sub)aspect.

#### Usage

```
aspectClasses

subAspectClasses

updateAspectClasses(aspectClasses, aspect)
```

#### Arguments

| | |
|---|---|
| `aspectClasses` | named character; accession names and aspect classes |
| `aspect` | named character; new accession names and aspect classes |

#### Format

An object of class `character` of length 14.

An object of class `character` of length 4.

#### Details

`updateAspectClasses` adds aspect classes to the list and avoids duplicate accession names and aspect classes.

#### Value

named character; accession names and aspect classes

#### Examples

```
aspectClasses

aspectClasses = updateAspectClasses(
  aspectClasses,
  c(bla="BlaAspect")
)

aspectClasses

aspectClasses = updateAspectClasses(
  aspectClasses,
  c(bla="BlaAspect", blubb="BubbAspect")
)

aspectClasses
subAspectClasses
```

---

`CartesianLayout` *Cartesian layout*

---

#### Description

This function creates a cartesian layout aspect, that stores coordinates of nodes.

#### Usage

```
createCartesianLayout(node, x, y, z = NULL, view = NULL)
```

#### Arguments

| | |
|---|---|
| `node` | integer; reference to node ids |
| `x` | numeric; x coordinate |
| `y` | numeric; y coordinate |
| `z` | numeric (optional); z coordinate |
| `view` | integer (optional); reference to subnetwork id of type view (`CyNetworkRelations`) |

#### Details

The layout of networks can be influenced by setting the node position manually. While x an y coordinates are mandatory, the z coordinates are optional and can, for example, be used to define the vertical stacking order of overlapping nodes.

Similar to Cytoscape `https://cytoscape.org/`, it is possible to define different views of the same network. The views itself are definded in `CySubNetworks` and `CyNetworkRelations`, and only referenced by a unique subnetwork id.

#### Value

*CartesianLayoutAspect* object

#### See Also

updateCartesianLayout;

#### Examples

```
## a minimal example
cartesianLayout = createCartesianLayout(
  node=0,
  x=5.5,
  y=200.3
)

## defining several coordinates at once
cartesianLayout = createCartesianLayout(
  node=c(0, 1),
  x=c(5.5, 110.1),
  y=c(200.3, 210.2)
)

## with all parameters
cartesianLayout = createCartesianLayout(
  node=c(0, 1, 0),
  x=c(5.5, 110.1, 7.2),
  y=c(200.3, 210.2, 13.9),
  z=c(-1, 3.1, NA),
  view=c(NA, NA, 1476)
)
```

---

### Convert-Names-and-Classes

*Convert aspect class name to RCX accession*

---

#### Description

The aspects in an RCX object are accessed by a `name` and return the aspect as an object of `cls`. To simplify the conversion between those, these functions return the corresponding name.

#### Usage

```
aspectName2Class(name)

aspectClass2Name(cls)
```

##### Arguments

| | |
|---|---|
| `name` | character; name of the RCX accession of the Aspect |
| `cls` | character; name of the aspect class |

##### Details

The following accessions/classes are available within the standard RCX implementation:

**accession name <=> class name**

```
metaData  <=>  MetaDataAspect
nodes  <=>  NodesAspect
edges  <=>  EdgesAspect
nodeAttributes  <=>  NodeAttributesAspect
edgeAttributes  <=>  EdgeAttributesAspect
networkAttributes  <=>  NetworkAttributesAspect
cartesianLayout  <=>  CartesianLayoutAspect
cyGroups  <=>  CyGroupsAspect
cyVisualProperties  <=>  CyVisualPropertiesAspect
cyHiddenAttributes  <=>  CyHiddenAttributesAspect
cyNetworkRelations  <=>  CyNetworkRelationsAspect
cySubNetworks  <=>  CySubNetworksAspect
cyTableColumn  <=>  CyTableColumnAspect```
```

##### Value

accession or class name

##### Examples

```
aspectName2Class("nodes")
##[1] "NodesAspect"

aspectClass2Name("NodesAspect")
##[1] "nodes"

aspectClasses

subAspectClasses
```

---

#### `countElements` *Number of elements in aspect*

---

##### Description

This function returns the number of elements in an aspect.

#### Usage

```
countElements(x)

## Default S3 method:
countElements(x)

## S3 method for class 'RCX'
countElements(x)

## S3 method for class 'CyVisualPropertiesAspect'
countElements(x)

## S3 method for class 'MetaDataAspect'
countElements(x)
```

#### Arguments

| | |
|---|---|
| x | an object of one of the aspect classes (e.g. Nodes) or RCX class. |

#### Details

Uses method dispatch, so the default methods already returns the correct number for the most aspect classes. This way it is easier to extend the data model.

There are only two exceptions in the core and Cytoscape aspects: Meta-data and CyVisualProperties.

Meta-data is a meta-aspect and therefore not included in Meta-data, and so its return is NA.

CyVisualProperties is the only aspect with a complex data structure beneath. Therefore its number of elements is just the number of how many of the following properties are set: network, nodes, edges, defaultNodes or defaultEdges.

#### Value

integer; number of elements. For RCX objects all counts are returned in the vector named by the aspect class.

#### See Also

hasIds(), idProperty(), refersTo(), referredBy(), maxId()

#### Examples

```
nodes = createNodes(name = c("ĈDK1","CDK2","CDK3"))
edges = createEdges(source = c(0,0), target = c(1,2))
rcx = createRCX(nodes = nodes, edges = edges)

countElements(nodes)

countElements(rcx)
```

custom-print *Print functions for RCX and aspect classes*

#### Description

These functions attempt to print RCX and aspect objects in a more readable form.

#### Usage

```
## S3 method for class 'MetaDataAspect'
print(x, ...)

## S3 method for class 'NodesAspect'
print(x, ...)

## S3 method for class 'EdgesAspect'
print(x, ...)

## S3 method for class 'NodeAttributesAspect'
print(x, ...)

## S3 method for class 'EdgeAttributesAspect'
print(x, ...)

## S3 method for class 'NetworkAttributesAspect'
print(x, ...)

## S3 method for class 'CartesianLayoutAspect'
print(x, ...)

## S3 method for class 'CyGroupsAspect'
print(x, ...)

## S3 method for class 'CyVisualPropertyProperties'
print(x, ...)

## S3 method for class 'CyVisualPropertyDependencies'
print(x, ...)

## S3 method for class 'CyVisualPropertyMappings'
print(x, ...)

## S3 method for class 'CyVisualProperty'
print(x, fields = c("all"), ...)

## S3 method for class 'CyVisualPropertiesAspect'
print(x, propertyOf = "all", fields = "all", ...)
```

```
## S3 method for class 'CyHiddenAttributesAspect'
print(x, ...)

## S3 method for class 'CyNetworkRelationsAspect'
print(x, ...)

## S3 method for class 'CySubNetworksAspect'
print(x, ...)

## S3 method for class 'CyTableColumnAspect'
print(x, ...)

## S3 method for class 'RCX'
print(x, inofficial = TRUE, ...)
```

#### Arguments

| | |
|---|---|
| x | aspect or RCX object |
| ... | further arguments passed to or from other methods. See `base::print()` |
| fields | character; Which fields should be shown, one of properties, dependencies, mappings or `all` |
| propertyOf | character; Which properties should be shown, one of network, nodes, edges, nodes:default, edges:default or `all` |
| inofficial | logical; if `FALSE` only the official aspects are printed |

#### Value

prints the object and returns it invisibly (invisible)

#### See Also

summary

#### Examples

```
rcx = createRCX(createNodes())
print(rcx)
```

---

CyGroups *Cytoscape Groups*

---

#### Description

This function is used to create Cytoscape "groups" aspects.

#### Usage

```
createCyGroups(
  id = NULL,
  name,
  nodes = NULL,
  externalEdges = NULL,
  internalEdges = NULL,
  collapsed = NULL
)
```

#### Arguments

| | |
|---|---|
| `id` | integer (optional); Cytoscape group ids |
| `name` | character; names of the groups |
| `nodes` | list of integers (optional); reference to node ids |
| `externalEdges` | list of integers (optional); the external edges making up the group; reference to edge ids |
| `internalEdges` | list of integers (optional); the internal edges making up the group; reference to edge ids |
| `collapsed` | logical (optional); whether the group is displayed as a single node |

#### Details

Cytoscape contributes aspects that organize subnetworks, attribute tables, and visual attributes for use by its own layout and analysis tools. Furthermore are the aspects used in web-based visualizations like within the NDEx platform.

Cytoscape groups allow to group a set of `Nodes` and corresponding internal and external `Edges` together, and represent a group as a single node in the visualization. A group is defined by its unique id, which must be an (positive) integer, which serves as reference to other aspects. If no ids are provided, they are created automatically.

#### Value

*CyGroupsAspect* object

#### See Also

`updateCyGroups`;

#### Examples

```
## a minimal example
cyGroups = createCyGroups(
  name = "Group One",
  nodes = list(c(1,2,3)),
  internalEdges = list(c(0,1))
)
```

```
## defining several groups at once
cyGroups = createCyGroups(
  name = c("Group One", "Group Two"),
  nodes = list(c(1,2,3), 0),
  internalEdges = list(c(0,1),NA)
)

## with all parameters
cyGroups = createCyGroups(
  id = c(0,1),
  name = c("Group One", "Group Two"),
  nodes = list(c(1,2,3), 0),
  internalEdges = list(c(0,1),NA),
  externalEdges = list(NA,c(1,3)),
  collapsed = c(TRUE,NA)
)
```

---

`CyHiddenAttributes` *Cytoscape hidden attributes*

---

#### Description

This function is used to create Cytoscape hidden attributes aspects.

#### Usage

```
createCyHiddenAttributes(
  name,
  value,
  dataType = NULL,
  isList = NULL,
  subnetworkId = NULL
)
```

#### Arguments

| | |
|---|---|
| `name` | character; key of the attribute |
| `value` | character or list of character; value of the attribute |
| `dataType` | character (optional); data type of the attribute |
| `isList` | logical (optional); a value should be considered as list |
| `subnetworkId` | integer (optional); refers to the IDs of a subnetwork aspect, but left blank (or `NA`) if root-network |

#### Details

Cytoscape contributes aspects that organize subnetworks, attribute tables, and visual attributes for use by its own layout and analysis tools. Furthermore are the aspects used in web-based visualizations like within the NDEx platform.

Besides network attributes, networks may have additional describing attributes originated from and used by Cytoscape. They are also defined in a key-value like manner, with the name of the attribute as key. The same attribute can also be defined for different subnetworks with different values. The values itself may differ in their data types, therefore it is necessary to provide the values as a list of the single values instead of a vector.

With *isList* it can be set, if a value should be considered as a list. This is of minor significance while working solely with RCX objects, unless it will be transformed to JSON. For some attributes it might be necessary that the values are encoded as lists, even if they contain only one element (or even zero elements). To force an element to be encoded correctly, this parameter can be used, for example: name="A", value=a, isList=T will be encoded in JSON as A=["a"].

#### Value

*CyHiddenAttributesAspect* object

#### See Also

`updateCyHiddenAttributes`;

#### Examples

```
## a minimal example
hiddenAttributes = createCyHiddenAttributes(
  name="A",
  value="a"
)

## defining several properties at once
hiddenAttributes = createCyHiddenAttributes(
  name=c("A", "B"),
  value=c("a","b")
)

## with characters and numbers mixed
hiddenAttributes = createCyHiddenAttributes(
  name=c("A","B"),
  value=list("a",3.14)
)

## force the number to be characters
hiddenAttributes = createCyHiddenAttributes(
  name=c("A","B"),
  value=list("a",3.14),
  dataType=c("character","character")
)
```

```
## with a list as input for one value
hiddenAttributes = createCyHiddenAttributes(
  name=c("A","B"),
  value=list(c("a1","a2"),
             "b")
)

## force "B" to be a list as well
hiddenAttributes = createCyHiddenAttributes(
  name=c("A","B"),
  value=list(c("a1","a2"),
             "b"),
  isList=c(TRUE,TRUE)
)

## with a subnetwork
hiddenAttributes = createCyHiddenAttributes(
  name=c("A","A"),
  value=c("a","a with subnetwork"),
  subnetworkId=c(NA,1)
)

## with all parameters
hiddenAttributes = createCyHiddenAttributes(
  name=c("A","A","B","B"),
  value=list(c("a1","a2"),
             "a with subnetwork",
             "b",
             "b with subnetwork"),
  isList=c(TRUE,FALSE,TRUE,FALSE),
  subnetworkId=c(NA,1,NA,1)
)
```

---

`CyNetworkRelations` *Cytoscape network relations*

---

#### Description

This function is used to create Cytoscape network relations aspects.

#### Usage

```
createCyNetworkRelations(child, parent = NULL, name = NULL, isView = FALSE)
```

#### Arguments

| | |
|---|---|
| `child` | integer; reference to subnetwork id |
| `parent` | integer (optional); reference to subnetwork id, but left blank (or `NA`) for root-network |
| `name` | character (optional); name of the subnetwork or view |
| `isView` | logical (optional); `TRUE` for views, else the network defines a subnetwork |

#### Details

Cytoscape contributes aspects that organize subnetworks, attribute tables, and visual attributes for use by its own layout and analysis tools. Furthermore are the aspects used in web-based visualizations like within the NDEx platform.

Cytoscape network relations define the relationship between the main network, subnetworks and views and also a name can be assigned to the relationship. Both, subnetworks and views are defined as subnetworks aspect, but their type is defined here by the *isView* property. The parent of a subnetwork or view can be an other subnetwork or the root network.

#### Value

*CyNetworkRelationsAspect* object

#### See Also

`updateCyNetworkRelations`;

#### Examples

```
## a minimal example
cyNetworkRelations = createCyNetworkRelations(
  child = 1
)

## with all parameters
cyNetworkRelations = createCyNetworkRelations(
  child = c(1,2),
  parent = c(NA,1),
  name = c("Network A",
           "View A"),
  isView = c(FALSE, TRUE)
)
```

---

`CySubNetworks` *Cytoscape subnetworks*

---

#### Description

This function is used to create Cytoscape subnetwork aspects.

#### Usage

```
createCySubNetworks(id, nodes = NULL, edges = NULL)
```

##### Arguments

| | |
|---|---|
| `id` | integer; subnetwork IDs |
| `nodes` | integer; reference to node id OR character "all" to refer to all nodes |
| `edges` | integer; reference to edge id OR character "all" to refer to all edges |

##### Details

Cytoscape contributes aspects that organize subnetworks, attribute tables, and visual attributes for use by its own layout and analysis tools. Furthermore are the aspects used in web-based visualizations like within the NDEx platform.

A group is defined by its unique *id*, which must be an (positive) integer, which serves as reference to other aspects. If no IDs are provided, they are created automatically.

Nodes and edges are referred by the IDs of the corresponding aspect. Unlike other aspects referring those IDs, the Cytoscape subnetwork aspect allows to refer to all nodes and edges using the keyword `all`.

The relationship between (sub-)networks and views, and also the type (subnetwork or view) is defined in `CyNetworkRelations`.

##### Value

*CySubNetworksAspect* object

##### See Also

`updateCySubNetworks`;

##### Examples

```
## a minimal example
cySubNetworks = createCySubNetworks(
  nodes = "all",
  edges = "all"
)

## defining several subnetworks at once
cySubNetworks = createCySubNetworks(
  nodes = list("all",
               c(1,2,3)),
  edges = list("all",
               c(0,2))
)

## with all parameters
cySubNetworks = createCySubNetworks(
  id = c(0,1),
  nodes = list("all",
               c(1,2,3)),
  edges = list("all",
               c(0,2))
```

```
)
```

---

CyTableColumn *Cytoscape table column properties*

---

#### Description

This function is used to create Cytoscape table column aspects.

#### Usage

```
createCyTableColumn(
  appliesTo,
  name,
  dataType = NULL,
  isList = NULL,
  subnetworkId = NULL
)
```

#### Arguments

| | |
|---|---|
| `appliesTo` | character; indicates whether this applies to "nodes", "edges" or "networks" table columns |
| `name` | character; key of the attribute |
| `dataType` | character (optional); data type of the attribute |
| `isList` | logical (optional); a value should be considered as list |
| `subnetworkId` | integer (optional); reference to subnetwork id, but left blank (or `NA`) if root-network |

#### Details

Cytoscape contributes aspects that organize subnetworks, attribute tables, and visual attributes for use by its own layout and analysis tools. Furthermore are the aspects used in web-based visualizations like within the NDEx platform.

These elements are used to represent Cytoscape table column labels and types. Its main use is to disambiguate empty table columns. The same attribute can also be defined for different subnetworks with different values. Cytoscape does not currently support table columns for the root network, but this is option is included here for consistency.

With *isList* it can be set, if a value should be considered as a list. This is of minor significance while working solely with RCX objects, unless it will be transformed to JSON.

#### Value

*CyTableColumnAspect* object

#### See Also

updateCyTableColumn; CyNetworkRelations

#### Examples

```
## a minimal example
tableColumn = createCyTableColumn(
  appliesTo="nodes",
  name="weight"
)

## defining several properties at once
tableColumn = createCyTableColumn(
  appliesTo=c("nodes","edges"),
  name=c("weight","weight")
)

## with all parameters
tableColumn = createCyTableColumn(
  appliesTo=c("nodes","edges","networks"),
  name=c("weight","weight","collapsed"),
  dataType=c("numeric","numeric","logical"),
  isList=c(FALSE,FALSE,TRUE),
  subnetworkId=c(NA,NA,1)
)
```

---

CyVisualProperties *Cytoscape visual properties (aspect)*

---

#### Description

This function is used to create Cytoscape visual properties aspects, that consists of CyVisualProperty objects for networks, nodes, edges, and default nodes and edges.

#### Usage

```
createCyVisualProperties(
  network = NULL,
  nodes = NULL,
  edges = NULL,
  defaultNodes = NULL,
  defaultEdges = NULL
)
```

#### Arguments

| | |
|---|---|
| network | CyVisualProperty object (optional); the visual properties of networks |
| nodes | CyVisualProperty object (optional); the visual properties of nodes |

| | |
|---|---|
| edges | CyVisualProperty object (optional); the visual properties of edges |
| defaultNodes | CyVisualProperty object (optional); the default visual properties of nodes |
| defaultEdges | CyVisualProperty object (optional); the default visual properties of edges |

#### Details

Cytoscape contributes aspects that organize subnetworks, attribute tables, and visual attributes for use by its own layout and analysis tools. Furthermore are the aspects used in web-based visualizations like within the NDEx platform.

The visual properties aspect is the only aspect (CyVisualProperties) with a complex structure. It is composed of several sub-property classes and consists of CyVisualProperty objects, that belong to, or more precisely describe one of the following network elements: *network*, *nodes*, *edges*, *defaultNodes* or *defaultEdges*.

A single visual property (i.e. CyVisualProperty object) organizes the information as *properties*, *dependencies* and *mappings*, as well as the single values *appliesTo* and *view*, that define the subnetwork or view to which the IDs apply.

Properties are CyVisualPropertyProperties objects, that hold information like `"NODE_FILL_COLOR" : "#26CCC9"` or `"NODE_LABEL_TRANSPARENCY" : "255"` in a key-value like manner.

Dependencies are CyVisualPropertyDependencies objects, that hold information about dependencies between visual properties. Currently there are only three dependencies supported:

- Lock Node with and height: `nodeSizeLocked = "false"`
- Fit Custom Graphics to node: `nodeCustomGraphicsSizeSync = "true"`
- Edge color to arrows: `arrowColorMatchesEdge = "false"`

Mappings are CyVisualPropertyMappings objects, that hold information as a triplet consisting of name, type and definition, like `"NODE_FILL_COLOR" : "DISCRETE" : "COL=molecule_type,T=string,K=0=miRNA,V=0=#F` "NODE_FILL_COLOR" : "CONTINUOUS" : "COL=gal1RGexp,T=double... or `"NODE_LABEL" : "PASSTHROUGH" : "COL=COMMON,T=string"`.

For further information about Cytoscape visual properties see the `Styles` topic of the official Cytoscape documentation: http://manual.cytoscape.org/en/stable/Styles.html

##### Structure of Cytoscape Visual Properties:

```
CyVisualProperties
|---network = CyVisualProperty
|---nodes = CyVisualProperty
|---edges = CyVisualProperty
|---defaultNodes = CyVisualProperty
|---defaultEdges = CyVisualProperty

CyVisualProperty
|---properties = CyVisualPropertyProperties
|    |--name
|    |--value
|---dependencies = CyVisualPropertyDependencies
|    |--name
```

```
|   |--value
|---mappings = CyVisualPropertyMappings
|   |--name
|   |--type
|   |--definition
|---appliesTo = <reference to subnetwork id>
|---view = <reference to subnetwork id>
```

#### Value

*CyVisualPropertiesAspect* object

#### See Also

updateCyVisualProperties, updateCyVisualProperty, getCyVisualProperty

#### Examples

```
## Prepare used properties
## Visual property: Properties
vpPropertyP1 = createCyVisualPropertyProperties(c(NODE_BORDER_STROKE="SOLID"))

## Visual property: Dependencies
vpPropertyD1 = createCyVisualPropertyDependencies(c(nodeSizeLocked="false"))

## Visual property: Mappings
vpPropertyM1 = createCyVisualPropertyMappings(c(NODE_FILL_COLOR="CONTINUOUS"),
                                               "COL=directed,T=boolean,K=0=true,V=0=ARROW")

## Create visual property object
vpProperty1 = createCyVisualProperty(properties=vpPropertyP1,
                                     dependencies=vpPropertyD1,
                                     mappings=vpPropertyM1)

## Create a visual properties aspect
## (using the same visual property object for simplicity)
createCyVisualProperties(network=vpProperty1,
                         nodes=vpProperty1,
                         edges=vpProperty1,
                         defaultNodes=vpProperty1,
                         defaultEdges=vpProperty1)
```

---

CyVisualProperty *Cytoscape visual property (object used in CyVisualProperties aspect)*

---

#### Description

This function is used to create Cytoscape visual property objects, that define networks, nodes, edges, and default nodes and edges in a CyVisualProperties aspect.

##### Usage

```
createCyVisualProperty(
  properties = NULL,
  dependencies = NULL,
  mappings = NULL,
  appliesTo = NULL,
  view = NULL
)
```

##### Arguments

| | |
|---|---|
| `properties` | a single or a list of `CyVisualPropertyProperties` object (optional); |
| `dependencies` | a single or a list of`CyVisualPropertyDependencies` object (optional); |
| `mappings` | a single or a list of`CyVisualPropertyMappings` object (optional); |
| `appliesTo` | integer (optional); might refer to the IDs of a subnetwork aspect, but CX documantation is unclear |
| `view` | integer (optional); might refer to the IDs of a subnetwork aspect that is a view, but CX documantation is unclear |

##### Details

Cytoscape contributes aspects that organize subnetworks, attribute tables, and visual attributes for use by its own layout and analysis tools. Furthermore are the aspects used in web-based visualizations like within the NDEx platform.

The visual properties aspect is the only aspect (`CyVisualProperties`) with a complex structure. It is composed of several sub-property classes and consists of `CyVisualProperty` objects, that belong to, or more precisely describe one of the following network elements: *network*, *nodes*, *edges*, *defaultNodes* or *defaultEdges*.

A single visual property (i.e. `CyVisualProperty` object) organizes the information as *properties*, *dependencies* and *mappings*, as well as the single values *appliesTo* and *view*, that define the subnetwork or view to which the IDs apply.

Properties are `CyVisualPropertyProperties` objects, that hold information like `"NODE_FILL_COLOR" : "#26CCC9"` or `"NODE_LABEL_TRANSPARENCY" : "255"` in a key-value like manner.

Dependencies are `CyVisualPropertyDependencies` objects, that hold information about dependencies between visual properties. Currently there are only three dependencies supported:

- Lock Node with and height: `nodeSizeLocked = "false"`
- Fit Custom Graphics to node: `nodeCustomGraphicsSizeSync = "true"`
- Edge color to arrows: `arrowColorMatchesEdge = "false"`

Mappings are `CyVisualPropertyMappings` objects, that hold information as a triplet consisting of name, type and definition, like `"NODE_FILL_COLOR" : "DISCRETE" : "COL=molecule_type,T=string,K=0=miRNA,V=0=#F` "NODE_FILL_COLOR" : "CONTINUOUS" : "COL=gal1RGexp,T=double... or `"NODE_LABEL" : "PASSTHROUGH" : "COL=COMMON,T=string"`.

For further information about Cytoscape visual properties see the `Styles` topic of the official Cytoscape documentation: http://manual.cytoscape.org/en/stable/Styles.html

**Structure of Cytoscape Visual Properties:**

```
CyVisualProperties
|---network = CyVisualProperty
|---nodes = CyVisualProperty
|---edges = CyVisualProperty
|---defaultNodes = CyVisualProperty
|---defaultEdges = CyVisualProperty

CyVisualProperty
|---properties = CyVisualPropertyProperties
|    |--name
|    |--value
|---dependencies = CyVisualPropertyDependencies
|    |--name
|    |--value
|---mappings = CyVisualPropertyMappings
|    |--name
|    |--type
|    |--definition
|---appliesTo = <reference to subnetwork id>
|---view = <reference to subnetwork id>
```

#### Value

CyVisualProperty object

#### See Also

updateCyVisualProperty, updateCyVisualProperties

#### Examples

```
## Prepare used properties
## Visual property: Properties
vpPropertyNamedValue = c(NODE_BORDER_STROKE="SOLID",
                         NODE_BORDER_WIDTH="1.5")
vpPropertyP = createCyVisualPropertyProperties(vpPropertyNamedValue)

## Visual property: Dependencies
vpDependencyNamedValue = c(nodeSizeLocked="false",
                           arrowColorMatchesEdge="true")
vpPropertyD = createCyVisualPropertyDependencies(vpDependencyNamedValue)

## Visual property: Mappings
vpMappingNamedType = c(NODE_FILL_COLOR="CONTINUOUS",
                       EDGE_TARGET_ARROW_SHAPE="DISCRETE")
vpMappingDefinition = c("COL=gal1RGexp,T=double,...",
                        "COL=directed,T=boolean,K=0=true,V=0=ARROW")
vpPropertyM = createCyVisualPropertyMappings(vpMappingNamedType,
                                             vpMappingDefinition)
```

```
## Create visual property object
createCyVisualProperty(properties=vpPropertyP,
                       dependencies=vpPropertyD,
                       mappings=vpPropertyM)

## Create visual property object with different subnetworks
createCyVisualProperty(properties=list(vpPropertyP,
                                       vpPropertyP),
                       dependencies=list(vpPropertyD,
                                         NA),
                       mappings=list(NA,
                                     vpPropertyM),
                       appliesTo = c(NA,
                                     1),
                       view = c(1,
                                NA))
```

---

CyVisualPropertyDependencies

*Create a object for dependency of Cytoscape Visual Properties (object used in CyVisualProperty)*

---

#### Description

This function is used to create aspects for mappings in Cytoscape visual properties. Networks, nodes, edges, and default nodes and edges mappings are realized as `CyVisualProperty` objects, that each consist of properties (`CyVisualPropertyProperties` objects), dependencies (**this here**) and mappings (`CyVisualPropertyMappings` objects).

#### Usage

```
createCyVisualPropertyDependencies(value, name = NULL)
```

#### Arguments

| | |
|---|---|
| `value` | character or named character; value of the dependencies |
| `name` | character (optional); name of the dependencies |

#### Details

Cytoscape contributes aspects that organize subnetworks, attribute tables, and visual attributes for use by its own layout and analysis tools. Furthermore are the aspects used in web-based visualizations like within the NDEx platform.

The visual properties aspect is the only aspect (`CyVisualProperties`) with a complex structure. It is composed of several sub-property classes and consists of `CyVisualProperty` objects, that belong to, or more precisely describe one of the following network elements: *network*, *nodes*, *edges*, *defaultNodes* or *defaultEdges*.

A single visual property (i.e. `CyVisualProperty` object) organizes the information as *properties*, *dependencies* and *mappings*, as well as the single values *appliesTo* and *view*, that define the subnetwork or view to which the IDs apply.

Properties are `CyVisualPropertyProperties` objects, that hold information like `"NODE_FILL_COLOR" : "#26CCC9"` or `"NODE_LABEL_TRANSPARENCY" : "255"` in a key-value like manner.

Dependencies are `CyVisualPropertyDependencies` objects, that hold information about dependencies between visual properties. Currently there are only three dependencies supported:

- Lock Node with and height: `nodeSizeLocked = "false"`
- Fit Custom Graphics to node: `nodeCustomGraphicsSizeSync = "true"`
- Edge color to arrows: `arrowColorMatchesEdge = "false"`

Mappings are `CyVisualPropertyMappings` objects, that hold information as a triplet consisting of name, type and definition, like `"NODE_FILL_COLOR" : "DISCRETE" : "COL=molecule_type,T=string,K=0=miRNA,V=0=#F` "NODE_FILL_COLOR" : "CONTINUOUS" : "COL=gal1RGexp,T=double... or `"NODE_LABEL" : "PASSTHROUGH" : "COL=COMMON,T=string"`.

For further information about Cytoscape visual properties see the `Styles` topic of the official Cytoscape documentation: http://manual.cytoscape.org/en/stable/Styles.html

#### Value

CyVisualPropertyDependencies object

#### Note

If *name* is not provided, the *names(value)* is used instead to infer the names.

#### See Also

`updateCyVisualProperty`, `updateCyVisualProperties`

#### Examples

```
## Using a named vector
vpDependencyNamedValue = c(nodeSizeLocked="false",
                           arrowColorMatchesEdge="true")
createCyVisualPropertyDependencies(vpDependencyNamedValue)

## Using two separate vectors
vpDependencyName = c("nodeSizeLocked",
                     "arrowColorMatchesEdge")
vpDependencyValue = c("false",
                      "true")
createCyVisualPropertyDependencies(vpDependencyValue,
                                   vpDependencyName)

# Result for either:
#                     name value
# 1         nodeSizeLocked false
# 2 arrowColorMatchesEdge  true
```

#### CyVisualPropertyMappings

*Create an object for mappings of Cytoscape Visual Properties (object used in CyVisualProperty)*

##### Description

This function is used to create objects for mappings in Cytoscape visual properties. Networks, nodes, edges, and default nodes and edges mappings are realized as `CyVisualProperty` objects, that each consist of properties (`CyVisualPropertyProperties` objects), dependencies (`CyVisualPropertyDependencies` objects) and mappings (**this here**).

##### Usage

```
createCyVisualPropertyMappings(type, definition, name = NULL)
```

##### Arguments

| | |
|---|---|
| `type` | character or named character; value of the mappings |
| `definition` | character; definitions of the mappings |
| `name` | character (optional); names of the mappings |

##### Details

Cytoscape contributes aspects that organize subnetworks, attribute tables, and visual attributes for use by its own layout and analysis tools. Furthermore are the aspects used in web-based visualizations like within the NDEx platform.

The visual properties aspect is the only aspect (`CyVisualProperties`) with a complex structure. It is composed of several sub-property classes and consists of `CyVisualProperty` objects, that belong to, or more precisely describe one of the following network elements: *network*, *nodes*, *edges*, *defaultNodes* or *defaultEdges*.

A single visual property (i.e. `CyVisualProperty` object) organizes the information as *properties*, *dependencies* and *mappings*, as well as the single values *appliesTo* and *view*, that define the subnetwork or view to which the IDs apply.

Properties are `CyVisualPropertyProperties` objects, that hold information like `"NODE_FILL_COLOR" : "#26CCC9"` or `"NODE_LABEL_TRANSPARENCY" : "255"` in a key-value like manner.

Dependencies are `CyVisualPropertyDependencies` objects, that hold information about dependencies between visual properties. Currently there are only three dependencies supported:

- Lock Node with and height: `nodeSizeLocked = "false"`
- Fit Custom Graphics to node: `nodeCustomGraphicsSizeSync = "true"`
- Edge color to arrows: `arrowColorMatchesEdge = "false"`

Mappings are `CyVisualPropertyMappings` objects, that hold information as a triplet consisting of name, type and definition, like `"NODE_FILL_COLOR" : "DISCRETE" : "COL=molecule_type,T=string,K=0=miRNA,V=0=#F` "NODE_FILL_COLOR" : "CONTINUOUS" : "COL=gal1RGexp,T=double... or `"NODE_LABEL" : "PASSTHROUGH" : "COL=COMMON,T=string"`.

For further information about Cytoscape visual properties see the `Styles` topic of the official Cytoscape documentation: http://manual.cytoscape.org/en/stable/Styles.html

**Structure of Cytoscape Visual Properties:**

```
CyVisualProperties
|---network = CyVisualProperty
|---nodes = CyVisualProperty
|---edges = CyVisualProperty
|---defaultNodes = CyVisualProperty
|---defaultEdges = CyVisualProperty

CyVisualProperty
|---properties = CyVisualPropertyProperties
|   |--name
|   |--value
|---dependencies = CyVisualPropertyDependencies
|   |--name
|   |--value
|---mappings = CyVisualPropertyMappings
|   |--name
|   |--type
|   |--definition
|---appliesTo = <reference to subnetwork id>
|---view = <reference to subnetwork id>
```

#### Value

CyVisualPropertyMappings object

#### Note

If *name* is not provided, the *names(type)* is used instead to infer the names.

#### See Also

`updateCyVisualProperty`, `updateCyVisualProperties`

#### Examples

```
## Using a named vector
vpMappingNamedType = c(NODE_FILL_COLOR="CONTINUOUS",
                       EDGE_TARGET_ARROW_SHAPE="DISCRETE")
vpMappingDefinition = c("COL=gal1RGexp,T=double,...",
                        "COL=directed,T=boolean,K=0=true,V=0=ARROW")
createCyVisualPropertyMappings(vpMappingNamedType,
```

```
                                  vpMappingDefinition)

## Using three separate vectors
vpMappingName = c("NODE_FILL_COLOR",
                  "EDGE_TARGET_ARROW_SHAPE")
vpMappingType = c("CONTINUOUS",
                  "DISCRETE")
createCyVisualPropertyMappings(vpMappingType,
                               vpMappingDefinition,
                               vpMappingName)

# Result for either:
#                      name       type                              definition
# 1         NODE_FILL_COLOR CONTINUOUS                COL=gal1RGexp,T=double,...
# 2 EDGE_TARGET_ARROW_SHAPE   DISCRETE COL=directed,T=boolean,K=0=true,V=0=ARROW
```

---

CyVisualPropertyProperties

*Create a object for properties of Cytoscape Visual Properties (object used in CyVisualProperty)*

---

#### Description

This function is used to create aspects for mappings in Cytoscape visual properties. Networks, nodes, edges, and default nodes and edges mappings are realized as `CyVisualProperty` objects, that each consist of properties (**this here**), dependencies (`CyVisualPropertyDependencies` objects) and mappings (`CyVisualPropertyMappings` objects).

#### Usage

```
createCyVisualPropertyProperties(value, name = NULL)
```

#### Arguments

| | |
|---|---|
| `value` | character or named character; value of the property |
| `name` | character (optional); name of the property |

#### Details

Cytoscape contributes aspects that organize subnetworks, attribute tables, and visual attributes for use by its own layout and analysis tools. Furthermore are the aspects used in web-based visualizations like within the NDEx platform.

The visual properties aspect is the only aspect (`CyVisualProperties`) with a complex structure. It is composed of several sub-property classes and consists of `CyVisualProperty` objects, that belong to, or more precisely describe one of the following network elements: *network*, *nodes*, *edges*, *defaultNodes* or *defaultEdges*.

A single visual property (i.e. `CyVisualProperty` object) organizes the information as *properties*, *dependencies* and *mappings*, as well as the single values *appliesTo* and *view*, that define the subnetwork or view to which the IDs apply.

Properties are `CyVisualPropertyProperties` objects, that hold information like `"NODE_FILL_COLOR" : "#26CCC9"` or `"NODE_LABEL_TRANSPARENCY" : "255"` in a key-value like manner.

Dependencies are `CyVisualPropertyDependencies` objects, that hold information about dependencies between visual properties. Currently there are only three dependencies supported:

- Lock Node with and height: `nodeSizeLocked = "false"`
- Fit Custom Graphics to node: `nodeCustomGraphicsSizeSync = "true"`
- Edge color to arrows: `arrowColorMatchesEdge = "false"`

Mappings are `CyVisualPropertyMappings` objects, that hold information as a triplet consisting of name, type and definition, like `"NODE_FILL_COLOR" : "DISCRETE" : "COL=molecule_type,T=string,K=0=miRNA,V=0=#F` "NODE_FILL_COLOR" : "CONTINUOUS" : "COL=gal1RGexp,T=double... or `"NODE_LABEL" : "PASSTHROUGH" : "COL=COMMON,T=string"`.

For further information about Cytoscape visual properties see the `Styles` topic of the official Cytoscape documentation: http://manual.cytoscape.org/en/stable/Styles.html

**Structure of Cytoscape Visual Properties:**

```
CyVisualProperties
|---network = CyVisualProperty
|---nodes = CyVisualProperty
|---edges = CyVisualProperty
|---defaultNodes = CyVisualProperty
|---defaultEdges = CyVisualProperty

CyVisualProperty
|---properties = CyVisualPropertyProperties
|    |--name
|    |--value
|---dependencies = CyVisualPropertyDependencies
|    |--name
|    |--value
|---mappings = CyVisualPropertyMappings
|    |--name
|    |--type
|    |--definition
|---appliesTo = <reference to subnetwork id>
|---view = <reference to subnetwork id>
```

#### Value

CyVisualPropertyProperties object

#### Note

If *name* is not provided, the *names(value)* is used instead to infer the names.

#### See Also

updateCyVisualProperty, updateCyVisualProperties

#### Examples

```
## Using a named vector
vpPropertyNamedValue = c(NODE_BORDER_STROKE="SOLID",
                         NODE_BORDER_WIDTH="1.5")
createCyVisualPropertyProperties(vpPropertyNamedValue)

## Using two separate vectors
vpPropertyName = c("NODE_BORDER_STROKE",
                   "NODE_BORDER_WIDTH")
vpPropertyValue = c("SOLID",
                    "1.5")
createCyVisualPropertyProperties(vpPropertyValue,
                                 vpPropertyName)

# Result for either:
#                  name value
# 1 NODE_BORDER_STROKE SOLID
# 2  NODE_BORDER_WIDTH   1.5
```

---

### EdgeAttributes *Edge attributes*

---

#### Description

This function creates an aspect for additional attributes of edges.

#### Usage

```
createEdgeAttributes(
  propertyOf,
  name,
  value,
  dataType = NULL,
  isList = NULL,
  subnetworkId = NULL
)
```

#### Arguments

| | |
|---|---|
| propertyOf | integer; reference to edge ids |
| name | character; key of the attribute |
| value | character; value of the attribute |
| dataType | character (optional); data type of the attribute |
| isList | logical (optional); a value should be considered as list |
| subnetworkId | integer (optional); reference to subnetwork id |

#### Details

Edges may have additional attributes besides a name and a representation. Those additional attributes reference a edge by its id and are defined in a key-value like manner, with the name of the attribute as key. The same attribute can also be defined for different subnetworks with different values. The values itself may also differ in their data types, therefore it is necessary to provide the values as a list of the single values instead of a vector.

With *isList* it can be set, if a value should be considered as a list. This is of minor significance while working solely with RCX objects, unless it will be transformed to JSON. For some attributes it might be necessary that the values are encoded as lists, even if they contain only one element (or even zero elements). To force an element to be encoded correctly, this parameter can be used, for example: name="A", value=a, isList=T will be encoded in JSON as A=["a"].

#### Value

*EdgeAttributesAspect* object

#### Note

The *propertyOf* parameter references the edge ids to which the attributes belong to. When adding an EdgeAttributesAspect object to an RCX object, those ids must be present in the `Edges` aspect, otherwise an error is raised.

#### See Also

`updateEdgeAttributes`

#### Examples

```
## a minimal example
edgeAttributes = createEdgeAttributes(
  propertyOf=1,
  name="A",
  value="a"
)

## defining several properties at once
edgeAttributes = createEdgeAttributes(
  propertyOf=c(1,1),
  name=c("A", "B"),
  value=c("a","b")
)

## with characters and numbers mixed
edgeAttributes = createEdgeAttributes(
  propertyOf=c(1,1),
  name=c("A","B"),
  value=list("a",3.14)
)

## force the number to be characters
```

```
edgeAttributes = createEdgeAttributes(
  propertyOf=c(1,1),
  name=c("A","B"),
  value=list("a",3.14),
  dataType=c("character","character")
)

## with a list as input for one value
edgeAttributes = createEdgeAttributes(
  propertyOf=c(1,1),
  name=c("A","B"),
  value=list(c("a1","a2"),
             "b")
)

## force "B" to be a list as well
edgeAttributes = createEdgeAttributes(
  propertyOf=c(1,1),
  name=c("A","B"),
  value=list(c("a1","a2"),
             "b"),
  isList=c(TRUE,TRUE)
)

## with a subnetwork
edgeAttributes = createEdgeAttributes(
  propertyOf=c(1,1),
  name=c("A","A"),
  value=c("a","a with subnetwork"),
  subnetworkId=c(NA,1)
)

## with all parameters
edgeAttributes = createEdgeAttributes(
  propertyOf=c(1,1,1,1),
  name=c("A","A","B","B"),
  value=list(c("a1","a2"),
             "a with subnetwork",
             "b",
             "b with subnetwork"),
  isList=c(TRUE,FALSE,TRUE,FALSE),
  subnetworkId=c(NA,1,NA,1)
)
```

---

Edges *Edges*

---

#### Description

This function creates edges between nodes in networks.

#### Usage

```
createEdges(id = NULL, source, target, interaction = NULL)
```

#### Arguments

| | |
|---|---|
| id | integer (optional); edge IDs |
| source | integer; reference to node id |
| target | integer; reference to node id |
| interaction | character (optional); type of interaction, eg. "binds" or "activates" |

#### Details

Edges are represented by *EdgesAspect* objects. Edges connect two nodes, which means that *source* and *target* must reference the IDs of nodes in a Nodes object. On creation, the IDs don't matter yet, but at least while adding the *EdgesAspect* object to an RCX-object, the *IDs* must be present in the nodes aspect of the RCX-object.

Similar to nodes, an edge also has a unique *id*, which must be an (positive) integer, which serves as reference to other aspects. If no IDs are provided, those are assigned automatically. Optionally, edges can have an interaction attribute to define the type of interaction between the nodes.

#### Value

*EdgesAspect* object

#### See Also

updateEdges for adding a *EdgesAspect* object to an *EdgesAspect* or *RCX* object

#### Examples

```
## create some simple edges
edges1 = createEdges(source=1, target=2)

## create edges with more information
edges2 = createEdges(id=c(3,2,4),
                     source=c(0,0,1),
                     target=c(1,2,2),
                     interaction=c("activates","inhibits", NA))
```

---

`getCyVisualProperty` *Get a Cytoscape visual property (object used in CyVisualProperties aspect) by appliesTo and view*

---

##### Description

This function helps filtering `CyVisualProperty` objects by appliesTo and view attributes (i.e. a unique combination of both). If nothing matches the searched pattern `NULL` is returned.

##### Usage

```
getCyVisualProperty(cyVisualProperty, appliesTo = NA, view = NA)
```

##### Arguments

| | |
|---|---|
| `cyVisualProperty` | `CyVisualProperty` object |
| `appliesTo` | integer (optional); value of appliesTo to filter for |
| `view` | integer (optional); value of view to filter for |

##### Details

Cytoscape contributes aspects that organize subnetworks, attribute tables, and visual attributes for use by its own layout and analysis tools. Furthermore are the aspects used in web-based visualizations like within the NDEx platform.

The visual properties aspect is the only aspect (`CyVisualProperties`) with a complex structure. It is composed of several sub-property classes and consists of `CyVisualProperty` objects, that belong to, or more precisely describe one of the following network elements: *network*, *nodes*, *edges*, *defaultNodes* or *defaultEdges*.

A single visual property (i.e. `CyVisualProperty` object) organizes the information as *properties*, *dependencies* and *mappings*, as well as the single values *appliesTo* and *view*, that define the subnetwork or view to which the IDs apply.

Properties are `CyVisualPropertyProperties` objects, that hold information like `"NODE_FILL_COLOR" : "#26CCC9"` or `"NODE_LABEL_TRANSPARENCY" : "255"` in a key-value like manner.

Dependencies are `CyVisualPropertyDependencies` objects, that hold information about dependencies between visual properties. Currently there are only three dependencies supported:

- Lock Node with and height: `nodeSizeLocked = "false"`
- Fit Custom Graphics to node: `nodeCustomGraphicsSizeSync = "true"`
- Edge color to arrows: `arrowColorMatchesEdge = "false"`

Mappings are `CyVisualPropertyMappings` objects, that hold information as a triplet consisting of name, type and definition, like `"NODE_FILL_COLOR" : "DISCRETE" : "COL=molecule_type,T=string,K=0=miRNA,V=0=#F` "NODE_FILL_COLOR" : "CONTINUOUS" : "COL=gal1RGexp,T=double... or `"NODE_LABEL" : "PASSTHROUGH" : "COL=COMMON,T=string"`.

For further information about Cytoscape visual properties see the `Styles` topic of the official Cytoscape documentation: http://manual.cytoscape.org/en/stable/Styles.html

**Structure of Cytoscape Visual Properties:**

```
CyVisualProperties
|---network = CyVisualProperty
|---nodes = CyVisualProperty
|---edges = CyVisualProperty
|---defaultNodes = CyVisualProperty
|---defaultEdges = CyVisualProperty

CyVisualProperty
|---properties = CyVisualPropertyProperties
|    |--name
|    |--value
|---dependencies = CyVisualPropertyDependencies
|    |--name
|    |--value
|---mappings = CyVisualPropertyMappings
|    |--name
|    |--type
|    |--definition
|---appliesTo = <reference to subnetwork id>
|---view = <reference to subnetwork id>
```

#### Value

CyVisualProperty object containing only one element, or NULL

#### See Also

updateCyVisualProperty, updateCyVisualProperties

#### Examples

```
## Visual property: Properties
vpPropertyP1 = createCyVisualPropertyProperties(c(NODE_BORDER_STROKE="SOLID"))

## Visual property: Dependencies
vpPropertyD1 = createCyVisualPropertyDependencies(c(nodeSizeLocked="false"))

## Visual property: Mappings
vpPropertyM1 = createCyVisualPropertyMappings(c(NODE_FILL_COLOR="CONTINUOUS"),
                                              "COL=directed,T=boolean,K=0=true,V=0=ARROW")

## Create visual property object
vpProperty = createCyVisualProperty(properties=list(vpPropertyP1,
                                                    vpPropertyP1,
                                                    vpPropertyP1),
                                    dependencies=list(vpPropertyD1,
                                                      vpPropertyD1,
                                                      NA),
                                    mappings=list(vpPropertyM1,
```

```
                                        NA,
                                        vpPropertyM1),
                          appliesTo = c(NA,
                                        NA,
                                        1)),
                          view = c(NA,
                                   1,
                                   1))

## Get VP for no subnetwork an no view
getCyVisualProperty(vpProperty)

getCyVisualProperty(vpProperty,
                    appliesTo = 1,
                    view = 1)
```

---

graphNEL *Convert an RCX object from and to an graphNEL object*

---

#### Description

Convert an RCX object to an graphNEL object

#### Usage

```
toGraphNEL(rcx, directed = FALSE)

fromGraphNEL(
  graphNEL,
  nodeId = "id",
  nodeName = "nodeName",
  nodeIgnore = c("name"),
  edgeId = "id",
  edgeInteraction = "edgeInteraction",
  edgeIgnore = c(),
  suppressWarning = FALSE
)
```

#### Arguments

| | |
|---|---|
| rcx | RCX object |
| directed | logical; whether the graph is directed |
| graphNEL | graphNEL object |
| nodeId | character; igraph attribute name used for node ids |
| nodeName | character; igraph attribute name used for node names |
| nodeIgnore | character; igraph attribute names that should be ignored |
| edgeId | character; igraph attribute name used for edge ids |

edgeInteraction
: character; igraph attribute name used for edge interaction

edgeIgnore
: character; igraph attribute names that should be ignored

suppressWarning
: logical; whether to suppress a warning message, if the validation of the RCX object fails

#### Details

In the graphNEL object the attributes are not separated from the graph like in RCX. Therefore, for converting an RCX object to an graphNEL object, and back, some adjustments in the naming of the attributes have to be made.

For nodes the name can be present in the nodes aspect, as name in the nodeAttributes aspect. Also name is used in graphNEL for naming the vertices. To avoid collisions in the conversion, the nodes name is saved in graphNEL as nodeName, while the nodeAttributes property name is saved as "attribute...name". These names are also used for the conversion back to RCX, but here the name used in the nodes aspect can be changed by the nodeName parameter.

Similar to the node name, if "represents" is present as property in nodeAttributes its name is changed to "attribute...represents".

The conversion of edges works analogously: If "interaction" is present as property in edgeAttributes its name is changed to "attribute...interaction".

Nodes and edges must have IDs in the RCX, but not in the graphNEL object. To define an vertex or edge attribute to be used as ID, the parameters nodeId and edgeId can be used to define ether an attribute name (default:"id") or set it to NULL to generate ID automatically.

The attributes also may have a special data type assigned. The data type then is saved by adding "...dataType" to the attribute name.

The cartesian layout is also stored in the graphNEL object. To make those graph vertex attributes distinguishable from nodeAttributes they are named "cartesianLayout...x", "cartesianLayout...y" and "cartesianLayout...z".

In the RCX attributes it is also possible to define a subnetwork, to which an attribute applies. Those attributes are added with "...123" added to its name, where "123" is the subnetwork id. The subnetwork id itself are added as graph graph attributes, and are named subnetwork...123...nodes" and "subnetwork...123...edges", where "123" is the subnetwork id.

Altogether, the conventions look as follows: "[attribute...]<name>[...<subnetwork>][...dataType]"

#### Value

graphNEL or RCX object

#### See Also

Igraph, igraph::as_graphnel()

#### Examples

```
## Read from a CX file
## reading the provided example network of the package
cxFile <- system.file(
 "extdata",
 "Imatinib-Inhibition-of-BCR-ABL-66a902f5-2022-11e9-bb6a-0ac135e8bacf.cx",
 package = "RCX"
)

rcx = readCX(cxFile)

## graphNEL can handle multi-edges, but only if the graph is directed and the
## source and target start and end not between the same nodes.
## Unfortunaltelly this is the case in our sample network.
## A quick fix is simply switching the direction of source and target
## for the multi-edges:
dubEdges = duplicated(rcx$edges[c("source","target")])

s = rcx$edges$source
rcx$edges$source[dubEdges] = rcx$edges$target[dubEdges]
rcx$edges$target[dubEdges] = s[dubEdges]

## convert the network to graphNEL
gNel = toGraphNEL(rcx, directed = TRUE)

## convert it back
rcxFromGraphNel = fromGraphNEL(gNel)
```

---

`hasIds` *IDs of an aspect*

---

#### Description

This function checks, if an aspect has IDs that may be referenced by other aspects.

By default aspects don't have IDs, so only the implemented classes have IDs. Aspects with IDs will be considered in the meta-data aspect to determine properties like: *idCounter* and *elementCount*.

#### Usage

```
hasIds(aspect)

## Default S3 method:
hasIds(aspect)

## S3 method for class 'NodesAspect'
hasIds(aspect)

## S3 method for class 'EdgesAspect'
```

```
hasIds(aspect)

## S3 method for class 'CyGroupsAspect'
hasIds(aspect)

## S3 method for class 'CySubNetworksAspect'
hasIds(aspect)
```

##### Arguments

| | |
|---|---|
| aspect | an object of one of the aspect classes (e.g. NodesAspect, EdgesAspect, etc.) |

##### Details

Uses method dispatch, so the default return is *FALSE* and only aspect classes with IDs are implemented. This way it is easier to extend the data model.

##### Value

logical

##### See Also

idProperty(), refersTo(), referredBy(), maxId()

##### Examples

```
edges = createEdges(source = c(0,0), target = c(1,2))
hasIds(edges)
```

---

#### idProperty *Name of the property of an aspect that is an ID*

---

##### Description

This function returns the name of the property, if an aspect uses IDs for its elements. As example, the aspect *NodesAspect* has the property *id* that represents the IDs of the aspect.

##### Usage

```
idProperty(aspect)

## Default S3 method:
idProperty(aspect)

## S3 method for class 'NodesAspect'
idProperty(aspect)
```

```
## S3 method for class 'EdgesAspect'
idProperty(aspect)

## S3 method for class 'CyGroupsAspect'
idProperty(aspect)

## S3 method for class 'CySubNetworksAspect'
idProperty(aspect)
```

#### Arguments

| | |
|---|---|
| aspect | an object of one of the aspect classes (e.g. NodesAspect, EdgesAspect, etc.) |

#### Details

By default aspects don't have IDs, so only the implemented classes have IDs. Aspects with IDs will be considered in the meta-data aspect to determine properties like: *idCounter* and *elementCount*.

Uses method dispatch, so the default return is *NULL* and only aspect classes with IDs are implemented. This way it is easier to extend the data model.

#### Value

character; Name of the ID property or *NULL*

#### See Also

`hasIds()`, `refersTo()`, `referredBy()`, `maxId()`

#### Examples

```
edges = createEdges(source = c(0,0), target = c(1,2))
idProperty(edges)
```

---

### Igraph *Convert an RCX object from and to an igraph object*

---

#### Description

Convert an RCX object to an igraph object

#### Usage

```
toIgraph(rcx, directed = FALSE)

fromIgraph(
  ig,
  nodeId = "id",
  nodeName = "nodeName",
```

```
  nodeIgnore = c("name"),
  edgeId = "id",
  edgeInteraction = "edgeInteraction",
  edgeIgnore = c(),
  suppressWarning = FALSE
)
```

#### Arguments

| | |
|---|---|
| rcx | RCX object |
| directed | logical; whether the graph is directed |
| ig | igraph object |
| nodeId | character; igraph attribute name used for node ids |
| nodeName | character; igraph attribute name used for node names |
| nodeIgnore | character; igraph attribute names that should be ignored |
| edgeId | character; igraph attribute name used for edge ids |
| edgeInteraction | character; igraph attribute name used for edge interaction |
| edgeIgnore | character; igraph attribute names that should be ignored |
| suppressWarning | logical; whether to suppress a warning message, if the validation of the RCX object fails |

#### Details

In the igraph object the attributes are not separated from the graph like in RCX. Therefore, for converting an RCX object to an igraph object, and back, some adjustments in the naming of the attributes have to be made.

For nodes the `name` can be present in the nodes aspect, as name in the nodeAttributes aspect. Also `name` is used in igraph for naming the vertices. To avoid collisions in the conversion, the nodes name is saved in igraph as `nodeName`, while the nodeAttributes property `name` is saved as `"attribute...name"`. These names are also used for the conversion back to RCX, but here the `name` used in the nodes aspect can be changed by the `nodeName` parameter.

Similar to the node name, if `"represents"` is present as property in nodeAttributes its name is changed to `"attribute...represents"`.

The conversion of edges works analogously: If `"interaction"` is present as property in edgeAttributes its name is changed to `"attribute...interaction"`.

Nodes and edges must have IDs in the RCX, but not in the igraph object. To define an vertex or edge attribute to be used as ID, the parameters `nodeId` and `edgeId` can be used to define ether an attribute name (default:"id") or set it to `NULL` to generate ID automatically.

The attributes also may have a special data type assigned. The data type then is saved by adding `"...dataType"` to the attribute name.

The cartesian layout is also stored in the igraph object. To make those igraph vertex attributes distinguishable from nodeAttributes they are named `"cartesianLayout...x"`, `"cartesianLayout...y"` and `"cartesianLayout...z"`.

In the RCX attributes it is also possible to define a subnetwork, to which an attribute applies. Those attributes are added with "...123" added to its name, where "123" is the subnetwork id. The subnetwork id itself are added as igraph graph attributes, and are named subnetwork...123...nodes" and "subnetwork...123...edges", where "123" is the subnetwork id.

Altogether, the conventions look as follows: "[attribute...]<name>[...<subnetwork>][...dataType]"

#### Value

igraph or RCX object

#### See Also

graphNEL

#### Examples

```
## Read from a CX file
## reading the provided example network of the package
cxFile <- system.file(
 "extdata",
 "Imatinib-Inhibition-of-BCR-ABL-66a902f5-2022-11e9-bb6a-0ac135e8bacf.cx",
 package = "RCX"
)

rcx = readCX(cxFile)

## convert the network to igraph
ig = toIgraph(rcx)

## convert it back
rcxFromIg = fromIgraph(ig)
```

---

jsonToRCX *Convert parsed JSON aspects to RCX*

---

#### Description

Functions to handle parsed JSON for the different aspects.

#### Usage

```
jsonToRCX(jsonData, verbose)

## Default S3 method:
jsonToRCX(jsonData, verbose)

## S3 method for class 'status'
jsonToRCX(jsonData, verbose)
```

```
## S3 method for class 'numberVerification'
jsonToRCX(jsonData, verbose)

## S3 method for class 'metaData'
jsonToRCX(jsonData, verbose)

## S3 method for class 'nodes'
jsonToRCX(jsonData, verbose)

## S3 method for class 'edges'
jsonToRCX(jsonData, verbose)

## S3 method for class 'nodeAttributes'
jsonToRCX(jsonData, verbose)

## S3 method for class 'edgeAttributes'
jsonToRCX(jsonData, verbose)

## S3 method for class 'networkAttributes'
jsonToRCX(jsonData, verbose)

## S3 method for class 'cartesianLayout'
jsonToRCX(jsonData, verbose)

## S3 method for class 'cyGroups'
jsonToRCX(jsonData, verbose)

## S3 method for class 'cyHiddenAttributes'
jsonToRCX(jsonData, verbose)

## S3 method for class 'cyNetworkRelations'
jsonToRCX(jsonData, verbose)

## S3 method for class 'cySubNetworks'
jsonToRCX(jsonData, verbose)

## S3 method for class 'cyTableColumn'
jsonToRCX(jsonData, verbose)

## S3 method for class 'cyVisualProperties'
jsonToRCX(jsonData, verbose)
```

#### Arguments

| | |
|---|---|
| jsonData | nested list from parsed JSON |
| verbose | logical; whether to print what is happening |

#### Details

These functions will be used in `processCX` to process the JSON data for every aspect. Each aspect is accessible in the CX-JSON by a particular accession name (i.e. its aspect name; see NDEx documentation: `https://home.ndexbio.org/data-model/`). This name is used as class to handle different aspects by method dispatch. This simplifies the extension of RCX for non-standard or self-defined aspects.

The CX-JSON is parsed to R data types using the jsonlite package as follows:

`jsonlite::fromJSON(cx,simplifyVector = FALSE)`

This results in a list of lists (of lists...) to avoid automatic data type conversions, which affect the correctness and usability of the data. Simplified JSON data for example NodeAttributes would be coerced into a data.frame, therefore the `value` column looses the format for data types other than `string`.

The *jsonData* will be a list with only one element named by the aspect: jsonData$<accessionName>

To access the parsed data for example nodes, this can be done by `jsonData$nodes`. The single aspects are then created using the corresponding **create** functions and combined to an RCX object using the corresponding **update** functions.

#### Value

created aspect or `NULL`

#### See Also

rcxToJson, toCX, readCX, writeCX

#### Examples

```
nodesJD = list(nodes=list(list("@id"=6, name="EGFR"),
                          list("@id"=7, name="CDK3")))
class(nodesJD) = c("nodes", class(nodesJD))

jsonToRCX(nodesJD, verbose=TRUE)
```

---

`maxId` *Highest ID of an aspect*

---

#### Description

This function returns the highest id used in an aspect, that has ids. As example, the aspect *NodesAspect* has the property *id* that must be a unique positive integer.

#### Usage

```
maxId(x)

## Default S3 method:
maxId(x)

## S3 method for class 'RCX'
maxId(x)
```

#### Arguments

| | |
|---|---|
| `x` | an object of one of the aspect classes (e.g. NodesAspect, EdgesAspect, etc.) or RCX class. |

#### Details

Uses method dispatch, so the default return is *NULL* and only aspect classes that have ids are implemented. This way it is easier to extend the data model.

#### Value

integer; Highest id. For RCX objects all highest ids are returned in the vector named by the aspect class.

#### See Also

`hasIds()`, `idProperty()`, `refersTo()`, `referredBy()`, `maxId()`

#### Examples

```
nodes = createNodes(name = c("ĈDK1","CDK2","CDK3"))
maxId(nodes)
```

---

`Meta-data` *Update RCX meta-data*

---

#### Description

The meta-data aspect contains meta-data about the aspects in the RCX object. It can be generated automatically based on the aspects present in a RCX object:

- for *version* and *consistencyGroup* default values are used
- *idCounter* is inferred with `hasIds` and `maxId` of an aspect
- *elementCount* is inferred from `countElements`
- *properties* is left out by default

##### Usage

```
updateMetaData(
  x,
  version = NULL,
  consistencyGroup = NULL,
  properties = NULL,
  aspectClasses = NULL
)

## S3 method for class 'RCX'
updateMetaData(
  x,
  version = NULL,
  consistencyGroup = NULL,
  properties = NULL,
  aspectClasses = NULL
)

## Default S3 method:
updateMetaData(
  x,
  version = NULL,
  consistencyGroup = NULL,
  properties = NULL,
  aspectClasses = NULL
)
```

##### Arguments

| | |
|---|---|
| `x` | RCX object or an aspect of a RCX; its class must be one of the standard RCX aspect classes |
| `version` | named character (optional); version of the aspect (default:"1.0") |
| `consistencyGroup` | named numerical (optional); consistency group of the aspect (default:1) |
| `properties` | named list (optional); properties that need to be fetched or updated independently of aspect data |
| `aspectClasses` | named character; accession names and aspect classes aspectClasses |

##### Details

If *version*, *consistencyGroup* or *properties* should have a different value, they can be set using a named vector (or named list for *properties*), where the name must be an accession name of that aspect in the `RCX-object` (e.g. `nodes` or `cyVisualProperties`).

Besides being a named list by aspect accession name, *properties* must also contain the single key-value pairs as a further named list. To remove all key-value pairs for one aspect, an empty list can be provided instead of a list with key-value pairs. To simplify adding of properties to a single aspect, there is the `updateMetaDataProperties` function available.

#### Value

MetaDataAspect object or RCX object

#### Note

The meta-data will always be updated automatically, when an aspect is added to or changed in the RCX object.

#### See Also

`updateMetaDataProperties`

#### Examples

```
## prepare RCX object:
nodes = createNodes(name = c("a","b","c","d","e","f"))
edges = createEdges(source=c(1,2,0,0,0,2),
                    target=c(2,3,1,2,5,4))
rcx = createRCX(nodes, edges)
cySubNetworks = createCySubNetworks(
  id = c(1,2),
  nodes = list("all", c(1,2,3)),
  edges = list("all", c(0,2))
)
rcx = updateCySubNetworks(rcx, cySubNetworks)

## update meta-data manually
rcx = updateMetaData(rcx)

## update meta-data with some values
rcx = updateMetaData(rcx,
                     version=c(edges="2.0"),
                     consistencyGroup=c(nodes=3),
                     properties=list(cySubNetworks=list(some="value",
                                                        another="VALUE"),
                                     edges=list(some="edge",
                                                another="EDGE")))

## remove all properties for edges
rcx = updateMetaData(rcx, properties=list(edges=list()))
```

---

`NetworkAttributes` *Network attributes*

---

#### Description

This function creates an aspect for attributes of a network.

#### Usage

```
createNetworkAttributes(
  name,
  value,
  dataType = NULL,
  isList = NULL,
  subnetworkId = NULL
)
```

#### Arguments

| | |
|---|---|
| name | character; key of the attribute |
| value | character; value of the attribute |
| dataType | character (optional); data type of the attribute |
| isList | logical (optional); a value should be considered as list |
| subnetworkId | integer (optional); reference to subnetwork id |

#### Details

Networks may have describing attributes, that are defined in a key-value like manner, with the name of the attribute as key. The same attribute can also be defined for different subnetworks with different values. The values itself may differ in their data types, therefore it is necessary to provide the values as a list of the single values instead of a vector.

With *isList* it can be set, if a value should be considered as a list. This is of minor significance while working solely with RCX objects, unless it will be transformed to JSON. For some attributes it might be necessary that the values are encoded as lists, even if they contain only one element (or even zero elements). To force an element to be encoded correctly, this parameter can be used, for example: name="A", value=a, isList=T will be encoded in JSON as A=["a"].

#### Value

NetworkAttributesAspect object

#### See Also

updateNetworkAttributes; NodeAttributes, EdgeAttributes

#### Examples

```
## a minimal example
networkAttributes = createNetworkAttributes(
  name="A",
  value="a"
)

## defining several properties at once
networkAttributes = createNetworkAttributes(
  name=c("A", "B"),
```

```
  value=c("a","b")
)

## with characters and numbers mixed
networkAttributes = createNetworkAttributes(
  name=c("A","B"),
  value=list("a",3.14)
)

## force the number to be characters
networkAttributes = createNetworkAttributes(
  name=c("A","B"),
  value=list("a",3.14),
  dataType=c("character","character")
)

## with a list as input for one value
networkAttributes = createNetworkAttributes(
  name=c("A","B"),
  value=list(c("a1","a2"),
             "b")
)

## force "B" to be a list as well
networkAttributes = createNetworkAttributes(
  name=c("A","B"),
  value=list(c("a1","a2"),
             "b"),
  isList=c(TRUE,TRUE)
)

## with a subnetwork
networkAttributes = createNetworkAttributes(
  name=c("A","A"),
  value=c("a","a with subnetwork"),
  subnetworkId=c(NA,1)
)

## with all parameters
networkAttributes = createNetworkAttributes(
  name=c("A","A","B","B"),
  value=list(c("a1","a2"),
             "a with subnetwork",
             "b",
             "b with subnetwork"),
  isList=c(TRUE,FALSE,TRUE,FALSE),
  subnetworkId=c(NA,1,NA,1)
)
```

---

#### `NodeAttributes` *Node attributes*

---

#### Description

This function creates an aspect for additional attributes of nodes.

#### Usage

```
createNodeAttributes(
  propertyOf,
  name,
  value,
  dataType = NULL,
  isList = NULL,
  subnetworkId = NULL
)
```

#### Arguments

| | |
|---|---|
| `propertyOf` | integer; reference to node ids |
| `name` | character; key of the attribute |
| `value` | character; value of the attribute |
| `dataType` | character (optional); data type of the attribute |
| `isList` | logical (optional); a value should be considered as list |
| `subnetworkId` | integer (optional); reference to subnetwork id |

#### Details

Nodes may have additional attributes besides a name and a representation. Those additional attributes reference a node by its id and are defined in a key-value like manner, with the name of the attribute as key. The same attribute can also be defined for different subnetworks with different values. The values itself may also differ in their data types, therefore it is necessary to provide the values as a list of the single values instead of a vector.

With *isList* it can be set, if a value should be considered as a list. This is of minor significance while working solely with RCX objects, unless it will be transformed to JSON. For some attributes it might be necessary that the values are encoded as lists, even if they contain only one element (or even zero elements). To force an element to be encoded correctly, this parameter can be used, for example: name="A", value=a, isList=T will be encoded in JSON as A=["a"].

#### Value

*NodeAttributesAspect* object

#### Note

The *propertyOf* parameter references the node ids to which the attributes belong to. When adding an NodeAttributesAspect object to an RCX object, those ids must be present in the `Nodes` aspect, otherwise an error is raised.

#### See Also

updateNodeAttributes, EdgeAttributes, NetworkAttributes

#### Examples

```
## a minimal example
nodeAttributes = createNodeAttributes(
  propertyOf=1,
  name="A",
  value="a"
)

## defining several properties at once
nodeAttributes = createNodeAttributes(
  propertyOf=c(1,1),
  name=c("A", "B"),
  value=c("a","b")
)

## with characters and numbers mixed
nodeAttributes = createNodeAttributes(
  propertyOf=c(1,1),
  name=c("A","B"),
  value=list("a",3.14)
)

## force the number to be characters
nodeAttributes = createNodeAttributes(
  propertyOf=c(1,1),
  name=c("A","B"),
  value=list("a",3.14),
  dataType=c("string","string")
)

## with a list as input for one value
nodeAttributes = createNodeAttributes(
  propertyOf=c(1,1),
  name=c("A","B"),
  value=list(c("a1","a2"),
             "b")
)

## force "B" to be a list as well
nodeAttributes = createNodeAttributes(
  propertyOf=c(1,1),
  name=c("A","B"),
  value=list(c("a1","a2"),
             "b"),
  isList=c(TRUE,TRUE)
)

## with a subnetwork
```

```
nodeAttributes = createNodeAttributes(
  propertyOf=c(1,1),
  name=c("A","A"),
  value=c("a","a with subnetwork"),
  subnetworkId=c(NA,1)
)

## with all parameters
nodeAttributes = createNodeAttributes(
  propertyOf=c(1,1,1,1,1,1),
  name=c("A","A","b","d","i","l"),
  value=list(c("a1","a2"),
             "a with subnetwork",
             TRUE,
             3.14,
             314,
             314),
  dataType=c("string","string","boolean","double","integer","long"),
  isList=c(TRUE,FALSE,FALSE,FALSE,FALSE,FALSE),
  subnetworkId=c(NA,1,NA,NA,NA,NA)
)
```

---

Nodes *Nodes*

---

#### Description

This function creates nodes for networks.

#### Usage

```
createNodes(id = NULL, name = NULL, represents = NULL)
```

#### Arguments

| | |
|---|---|
| `id` | integer (optional); node IDs |
| `name` | character (optional); names of the nodes |
| `represents` | character (optional); representation, e.g. a link to another database |

#### Details

Nodes are represented by *NodesAspect* objects. A single node is defined by its unique *id*, which must be an (positive) integer, which serves as reference to other aspects. Optionally, nodes can have a name and a represents attribute. If no IDs are provided, but either names or representations (or both) IDs are assigned automatically. To be valid, a nodes aspect must contain at least one node. However, if no parameters are set (i.e. *id*, *name* and *represents* = `NULL`) there is still one node created with neither name nor representation, just an ID. The *NodesAspect* is the only mandatory aspect for an `RCX-object`.

##### Value

*NodesAspect* object

##### See Also

updateNodes, RCX-object

##### Examples

```
## a minimal example
nodes = createNodes()

## ids will be generated
nodes = createNodes(name = c("a","b","c"))

## with all parameters
nodes = createNodes(id=c(1, 2, 3),
                    name=c("CDK1", "CDK2", "CDK3"),
                    represents=c("HGNC:CDK1",
                                 "Uniprot:P24941",
                                 "Ensembl:ENSG00000250506"))
```

---

RCX *R package implementing the Cytoscape Exchange (CX) format*

---

##### Description

Create, handle, validate, visualize and convert networks in the Cytoscape exchange (CX) format to standard data types and objects.

##### Details

The CX format is also used by the NDEx platform, a online commons for biological networks, and the network visualization software Cytocape.

`browseVignettes("RCy3")`

##### Author(s)

Florian Auer <>

RCX-object *Create an RCX object from aspects*

#### Description

An RCX object consists of several aspects, but at least one node in the nodes aspect. The network can either created by creating every single aspect first and the create the network with all aspects present, or by creating the aspect only with the nodes and adding the remaining aspects one by one.

#### Usage

```
createRCX(
  nodes,
  edges,
  nodeAttributes,
  edgeAttributes,
  networkAttributes,
  cartesianLayout,
  cyGroups,
  cyVisualProperties,
  cyHiddenAttributes,
  cyNetworkRelations,
  cySubNetworks,
  cyTableColumn,
  checkReferences = TRUE
)
```

#### Arguments

| | |
|---|---|
| nodes | Nodes aspect; |
| edges | Edges aspect (optional); |
| nodeAttributes | NodeAttributes aspect (optional); |
| edgeAttributes | EdgeAttributes aspect (optional); |
| networkAttributes | NetworkAttributes aspect (optional); |
| cartesianLayout | CartesianLayout aspect (optional); |
| cyGroups | CyGroups aspect (optional); |
| cyVisualProperties | CyVisualProperties aspect (optional); |
| cyHiddenAttributes | CyHiddenAttributes aspect (optional); |
| cyNetworkRelations | CyNetworkRelations aspect (optional); |

cySubNetworks CySubNetworks aspect (optional);

cyTableColumn CyTableColumn aspect (optional);

checkReferences
logical; whether to check if references to other aspects are present in the RCX object

#### Details

vignette("01. RCX -an R package implementing the Cytoscape Exchange (CX) format",package = "RCX") vignette("02. Creating RCX from scratch",package = "RCX") vignette("Appendix: The RCX and CX Data Model",package = "RCX")

#### Value

RCX object

#### Examples

```
## minimal example
rcx = createRCX(createNodes())

## create by aspect
nodes = createNodes(name = c("a","b","c"))
edges = createEdges(source=c(0,0), target=c(1,2))

nodeAttributes = createNodeAttributes(
  propertyOf=c(1,1),
  name=c("A","B"),
  value=c("a","b")
)

edgeAttributes = createEdgeAttributes(
  propertyOf=c(0,0),
  name=c("A", "B"),
  value=c("a","b")
)

networkAttributes = createNetworkAttributes(
  name=c("A","B"),
  value=list("a",3.14)
)

cartesianLayout = createCartesianLayout(
  node=c(0, 1),
  x=c(5.5, 110.1),
  y=c(200.3, 210.2)
)

cyGroups = createCyGroups(
  name = c("Group One", "Group Two"),
  nodes = list(c(0,1), 0)
```

```
)

vpPropertyP = createCyVisualPropertyProperties(c(NODE_BORDER_STROKE="SOLID"))
vpPropertyD = createCyVisualPropertyDependencies(c(nodeSizeLocked="false"))
vpPropertyM = createCyVisualPropertyMappings(c(NODE_FILL_COLOR="CONTINUOUS"),
                                            "COL=directed,T=boolean,K=0=true,V=0=ARROW")
vpProperty = createCyVisualProperty(properties=vpPropertyP,
                                    dependencies=vpPropertyD,
                                    mappings=vpPropertyM)

cyVisualProperties = createCyVisualProperties(nodes=vpProperty)

cyHiddenAttributes = createCyHiddenAttributes(
  name=c("A","B"),
  value=list(c("a1","a2"), "b")
)

cyNetworkRelations = createCyNetworkRelations(
  child = c(0,1),
  name = c("Network A", NA)
)

cySubNetworks = createCySubNetworks(
  nodes = list("all", c(0,1,2)),
  edges = list("all", c(0,1))
)

cyTableColumn = createCyTableColumn(
  appliesTo=c("nodes","edges","networks"),
  name=c("weight","weight","collapsed"),
  dataType=c("double","double","boolean")
)

rcx = createRCX(nodes, edges,
                nodeAttributes, edgeAttributes,
                networkAttributes,
                cartesianLayout,
                cyGroups,
                cyVisualProperties,
                cyHiddenAttributes,
                cyNetworkRelations,
                cySubNetworks,
                cyTableColumn)

## create all at once
rcx = createRCX(
  createNodes(name = c("a","b","c")),
  createEdges(source=c(0,0), target=c(1,2)),
  createNodeAttributes(
    propertyOf=c(1,1),
    name=c("A","B"),
    value=c("a","b")
  ),
```

```
    createEdgeAttributes(
      propertyOf=c(0,0),
      name=c("A", "B"),
      value=c("a","b")
    ),
    networkAttributes = createNetworkAttributes(
      name=c("A","B"),
      value=list("a",3.14)
    ),
    cartesianLayout = createCartesianLayout(
      node=c(0, 1),
      x=c(5.5, 110.1),
      y=c(200.3, 210.2)
    ),
    createCyGroups(
      name = c("Group One", "Group Two"),
      nodes = list(c(0,1), 0)
    ),
    createCyVisualProperties(
      nodes=createCyVisualProperty(
        properties=createCyVisualPropertyProperties(
          c(NODE_BORDER_STROKE="SOLID")
        ),
        dependencies=createCyVisualPropertyDependencies(
          c(nodeSizeLocked="false")
        ),
        mappings=createCyVisualPropertyMappings(
          c(NODE_FILL_COLOR="CONTINUOUS"),
          "COL=directed,T=boolean,K=0=true,V=0=ARROW")
      )
    ),
    createCyHiddenAttributes(
      name=c("A","B"),
      value=list(c("a1","a2"), "b")
    ),
    createCyNetworkRelations(
      child = c(0,1),
      name = c("Network A", NA)
    ),
    createCySubNetworks(
      nodes = list("all", c(0,1,2)),
      edges = list("all", c(0,1))
    ),
    createCyTableColumn(
      appliesTo=c("nodes","edges","networks"),
      name=c("weight","weight","collapsed"),
      dataType=c("double","double","boolean")
    )
  )
```

#### rcxToJson — *Convert RCX aspects to JSON*

#### Description

Functions for converting the different aspects to JSON following the CX data structure definition (see NDEx documentation: https://home.ndexbio.org/data-model/).

#### Usage

```
rcxToJson(aspect, verbose = FALSE, ...)

## Default S3 method:
rcxToJson(aspect, verbose = FALSE, ...)

## S3 method for class 'MetaDataAspect'
rcxToJson(aspect, verbose = FALSE, ...)

## S3 method for class 'NodesAspect'
rcxToJson(aspect, verbose = FALSE, ...)

## S3 method for class 'EdgesAspect'
rcxToJson(aspect, verbose = FALSE, ...)

## S3 method for class 'NodeAttributesAspect'
rcxToJson(aspect, verbose = FALSE, ...)

## S3 method for class 'EdgeAttributesAspect'
rcxToJson(aspect, verbose = FALSE, ...)

## S3 method for class 'NetworkAttributesAspect'
rcxToJson(aspect, verbose = FALSE, ...)

## S3 method for class 'CartesianLayoutAspect'
rcxToJson(aspect, verbose = FALSE, ...)

## S3 method for class 'CyGroupsAspect'
rcxToJson(aspect, verbose = FALSE, ...)

## S3 method for class 'CyHiddenAttributesAspect'
rcxToJson(aspect, verbose = FALSE, ...)

## S3 method for class 'CyNetworkRelationsAspect'
rcxToJson(aspect, verbose = FALSE, ...)

## S3 method for class 'CySubNetworksAspect'
rcxToJson(aspect, verbose = FALSE, ...)

## S3 method for class 'CyTableColumnAspect'
rcxToJson(aspect, verbose = FALSE, ...)

## S3 method for class 'CyVisualPropertiesAspect'
```

```
rcxToJson(aspect, verbose = FALSE, ...)

## S3 method for class 'CyVisualProperty'
rcxToJson(aspect, verbose = FALSE, propertyOf = "", ...)

## S3 method for class 'CyVisualPropertyProperties'
rcxToJson(aspect, verbose = FALSE, ...)

## S3 method for class 'CyVisualPropertyDependencies'
rcxToJson(aspect, verbose = FALSE, ...)

## S3 method for class 'CyVisualPropertyMappings'
rcxToJson(aspect, verbose = FALSE, ...)
```

#### Arguments

| | |
|---|---|
| aspect | aspects of an RCX object |
| verbose | logical; whether to print what is happening |
| ... | additional parameters, that might needed for extending |
| propertyOf | character; provide propertyOf (only necessary for CyVisualProperty) |

#### Details

For converting RCX objects to JSON, each aspect is processed by a generic function for its aspect class. Those functions return a character only containing the JSON of this aspect, which is then combined by toCX to be a valid CX data structure.

To support the conversion for non-standard or own-defined aspects, generic functions for those aspect classes have to be implemented.

#### Value

character; JSON of an aspect

#### See Also

toCX, writeCX, jsonToRCX, readCX

#### Examples

```
nodes = createNodes(name = c("a","b","c","d","e","f"))
rcxToJson(nodes)
```

`readCX` *Read CX from file, parse the JSON and convert it to an RCX object*

#### Description

The `readCX` function combines three sub-task:

- read the JSON from file
- parse the JSON
- process the contained aspects to create an RCX object

#### Usage

```
readCX(file, verbose = FALSE, aspectClasses = NULL)

readJSON(file, verbose = FALSE)

parseJSON(json, verbose = FALSE)

processCX(aspectList, verbose = FALSE, aspectClasses = NULL)
```

#### Arguments

| | |
|---|---|
| `file` | character; the name of the file which the data are to be read from |
| `verbose` | logical; whether to print what is happening |
| `aspectClasses` | named character; accession names and aspect classes aspectClasses |
| `json` | character; raw JSON data |
| `aspectList` | list; list containing the aspect data (parsed JSON) |

#### Details

If any errors occur during this process, the single steps can be performed individually. This also allows to skip certain steps, for example if the JSON data is already availabe as text, there is no need to save it as file and read it again.

**Read the JSON from file:**

The `readJSON` function only read the content of a text file and returns it as a simple character vector.

**Parse the JSON:**

The `parseJSON` function uses the jsonlite package, to parse JSON text:

`jsonlite::fromJSON(cx,simplifyVector = FALSE)`

The result is a list containing the aspect data as elements. If, for some reason, the JSON is not valid, the jsonlite package raises an error.

**Process the contained aspects to create an RCX object:**

With the processCX function, the single elements from the previous list will be processed with the jsonToRCX functions, which creating objects for the single aspects. The standard CX aspects are processed by generic functions named by the aspect names of the CX data structure, e.g. jsonToRCX.nodeAttributes for the samely named CX aspect the corresponding NodeAttributesAspect in RCX (see also vignette("02. The RCX and CX Data Model") or NDEx documentation: https://home.ndexbio.org/data-model/).

The CX network may contain additional aspects besides the officially defined ones. This includes self defined or deprecated aspects, that sill can be found in the networks at the NDEx platform. By default, those aspects are simply omitted. In those cases, the setting *verbose* to TRUE is a good idea to see, which aspects cannot be processed this package.

Those not processable aspects can be handled individually, but it is advisable to extend the jsonToRCX functions by implementing own versions for those aspects. Additionally, the **update** functions have to be implemented to add the newly generated aspect objects to RCX object (see e.g. updateNodes or updateEdges). Therefore, the function also have to be named "update<aspect-name>, where aspect-name is the capitalized version of the name used in the CX. (see also vignette("03. Extending the RCX Data Model")

#### Value

RCX object

#### Functions

- readJSON: Reads the CX/JSON from file and returns the content as text
- parseJSON: Parses the JSON text and returns a list with the aspect data
- processCX: Processes the list of aspect data and creates an RCX

#### See Also

jsonToRCX, writeCX

#### Examples

```
cxFile = system.file(
  "extdata",
  "Imatinib-Inhibition-of-BCR-ABL-66a902f5-2022-11e9-bb6a-0ac135e8bacf.cx",
  package = "RCX"
)

rcx = readCX(cxFile)

## OR:

json = readJSON(cxFile)
aspectList = parseJSON(json)
rcx = processCX(aspectList)
```

#### referredBy — *List the aspects that are refered by an other aspect*

##### Description

This function returns a list of all aspects with all present aspects, that refer to it. As example, the aspect *NodesAspect* is refered by the property *source* and *target* of the *EdgesAspect* aspect.

##### Usage

```
referredBy(rcx, aspectClasses = NULL)
```

##### Arguments

| | |
|---|---|
| `rcx` | an object of one of the aspect classes (e.g. NodesAspect, EdgesAspect, etc.) |
| `aspectClasses` | named character; accession names and aspect classes aspectClasses |

##### Value

named list; Aspect class names with names of aspect classes, that refer to them.

##### Note

Uses `hasIds()` and `refersTo()` to determine the referring aspects.

##### See Also

`hasIds()`, `idProperty()`, `refersTo()`, `maxId()`

##### Examples

```
nodes = createNodes(name = c("ĈDK1","CDK2","CDK3"))
edges = createEdges(source = c(0,0), target = c(1,2))
rcx = createRCX(nodes = nodes, edges = edges)

referredBy(rcx)
```

---

refersTo *Name of the property of an aspect that is an ID*

---

#### Description

This function returns the name of the property and the aspect class it refers to. As example, the aspect *EdgesAspect* has the property *source* that refers to the *ids* of the *NodesAspect* aspect.

#### Usage

```
refersTo(aspect)

## Default S3 method:
refersTo(aspect)

## S3 method for class 'EdgesAspect'
refersTo(aspect)

## S3 method for class 'NodeAttributesAspect'
refersTo(aspect)

## S3 method for class 'EdgeAttributesAspect'
refersTo(aspect)

## S3 method for class 'CartesianLayoutAspect'
refersTo(aspect)

## S3 method for class 'CyGroupsAspect'
refersTo(aspect)

## S3 method for class 'CyVisualPropertiesAspect'
refersTo(aspect)

## S3 method for class 'CySubNetworksAspect'
refersTo(aspect)
```

#### Arguments

| | |
|---|---|
| aspect | an object of one of the aspect classes (e.g. NodesAspect, EdgesAspect, etc.) |

#### Details

Uses method dispatch, so the default return is *NULL* and only aspect classes that refer to other aspects are implemented. This way it is easier to extend the data model.

#### Value

named list; Name of the refering property and aspect class name.

##### Methods (by class)

- `default`: of default returns *NULL*
- `EdgesAspect`: of EdgesAspect refers to id by *source* and *target*
- `NodeAttributesAspect`: of NodeAttributesAspect refers to id by *propertyOf* and to id by *subnetworkId*
- `EdgeAttributesAspect`: of EdgeAttributesAspect refers to id by *propertyOf* and to id by *subnetworkId*
- `CartesianLayoutAspect`: of CartesianLayoutAspect refers to id by *node* and to id by *view*
- `CyGroupsAspect`: of CyGroupsAspect refers to id by *nodes* and to id by *externalEdges* and *internalEdges*
- `CyVisualPropertiesAspect`: of CyVisualPropertiesAspect refers to id by *appliesTo* of the sub-aspects
- `CySubNetworksAspect`: of refers to id by *nodes* and to id by *edges*

##### See Also

`hasIds()`, `idProperty()`, `referredBy()`, `maxId()`

##### Examples

```
edges = createEdges(source = c(0,0), target = c(1,2))
refersTo(edges)
```

---

`summary` *RCX and aspect summary*

---

##### Description

`summary` is a generic function used to produce result summaries of the RCX object. The function invokes particular methods which depend on the class of the first argument.

##### Usage

```
## S3 method for class 'RCX'
summary(object, ...)

## S3 method for class 'MetaDataAspect'
summary(object, ...)

## S3 method for class 'NodesAspect'
summary(object, ...)

## S3 method for class 'EdgesAspect'
summary(object, ...)
```

```
## S3 method for class 'NodeAttributesAspect'
summary(object, ...)

## S3 method for class 'EdgeAttributesAspect'
summary(object, ...)

## S3 method for class 'NetworkAttributesAspect'
summary(object, ...)

## S3 method for class 'CartesianLayoutAspect'
summary(object, ...)

## S3 method for class 'CyGroupsAspect'
summary(object, ...)

## S3 method for class 'CyHiddenAttributesAspect'
summary(object, ...)

## S3 method for class 'CyNetworkRelationsAspect'
summary(object, ...)

## S3 method for class 'CySubNetworksAspect'
summary(object, ...)

## S3 method for class 'CyTableColumnAspect'
summary(object, ...)

## S3 method for class 'CyVisualPropertiesAspect'
summary(object, ...)

## S3 method for class 'CyVisualProperty'
summary(object, ...)

## S3 method for class 'AspectIdColumn'
summary(object, ...)

## S3 method for class 'AspectRefColumn'
summary(object, ...)

## S3 method for class 'AspectReqRefColumn'
summary(object, ...)

## S3 method for class 'AspectValueColumn'
summary(object, ...)

## S3 method for class 'AspectAttributeColumn'
summary(object, ...)
```

```
## S3 method for class 'AspectListLengthColumn'
summary(object, ...)
```

#### Arguments

| | |
|---|---|
| object | an object; RCX object or aspect (or column of data.frame) |
| ... | additional arguments affecting the summary produced. |

#### Details

The form of the returned summary depends on the class of its argument, therefore it is possible to summarize RCX objects and their single aspects.

To enhance readability of the summary, some additional classes have `summary` functions, that are used to show for example ids of an aspect, required and optional references to ids of aspects, or the number of elements in lists.

#### Value

object summary as list

#### Methods (by class)

- `AspectIdColumn`: Summarize an id property
- `AspectRefColumn`: Summarize an optional property, that references the ids of an other aspect
- `AspectReqRefColumn`: Summarize a required property, that references the ids of an other aspect
- `AspectValueColumn`: Summarize the occurrences of the different elements in the property
- `AspectAttributeColumn`: Summarize the different attributes in the property
- `AspectListLengthColumn`: The property is a list of vectors, so summarize the length of the vectors

#### Examples

```
rcx = createRCX(
  nodes = createNodes(name = c("a","b","c")),
  edges = createEdges(source=1, target=2)
)

summary(rcx)
```

| toCX | *Convert an RCX object to CX (JSON)* |
|---|---|

#### Description

This function converts an RCX object to JSON in a valid CX data structure (see NDEx documentation: https://home.ndexbio.org/data-model/).

#### Usage

```
toCX(rcx, verbose = FALSE, pretty = FALSE)
```

#### Arguments

| | |
|---|---|
| `rcx` | RCX object |
| `verbose` | logical; whether to print what is happening |
| `pretty` | logical; adds indentation whitespace to JSON output. Can be TRUE/FALSE or a number specifying the number of spaces to indent. See `jsonlite::prettify()` |

#### Details

The single aspects of the RCX object are processed by generic functions of rcxToJson for each aspect class. Therefore, not only the single aspects are converted to JSON, but also necessary additional aspects are added, so the resulting CX is accepted by the NDEx platform (https://ndexbio.org/):

- *numberVerication* shows the supported maximal number
- *status* is needed at the end to show, that no errors have occurred while creation

If the RCX object contains additional aspects besides the officially defined ones, the corresponding rcxToJson functions for those aspect classes have to be implemented in order to include them in the resulting CX.

#### Value

CX (JSON) text

#### See Also

toCX, rcxToJson, readCX, writeCX

#### Examples

```
rcx = createRCX(
  nodes = createNodes(
    name = LETTERS[seq_len(10)]
  ),
  edges = createEdges(
    source=c(1,2),
    target = c(2,3)
  )
)

json = toCX(rcx, pretty=TRUE)
```

---

updateCartesianLayout    *Update Cartesian Layouts*

---

#### Description

This functions add a cartesian layout in the form of a CartesianLayout object to an other CartesianLayout or an RCX object.

#### Usage

```
updateCartesianLayout(
  x,
  cartesianLayout,
  replace = TRUE,
  stopOnDuplicates = FALSE,
  ...
)

## S3 method for class 'CartesianLayoutAspect'
updateCartesianLayout(
  x,
  cartesianLayout,
  replace = TRUE,
  stopOnDuplicates = FALSE,
  ...
)

## S3 method for class 'RCX'
updateCartesianLayout(
  x,
  cartesianLayout,
  replace = TRUE,
  stopOnDuplicates = FALSE,
  checkReferences = TRUE,
  ...
)
```

#### Arguments

- `x` — RCX or `CartesianLayout` object; (to which the new layout will be added)
- `cartesianLayout` — `CartesianLayout` object; (the layout, that will be added)
- `replace` — logical; if existing values are updated (or ignored)
- `stopOnDuplicates` — logical; whether to stop, if duplicates in `nodes` (and `view` if present) column are found
- `...` — additional parameters
- `checkReferences` — logical; whether to check if references to other aspects are present in the RCX object

#### Details

Networks, or more precisely its nodes may have a cartesian layout, that is represented as `CartesianLayout` object. `CartesianLayout` objects can be added to an RCX or an other `CartesianLayout` object.

In the case, that a `CartesianLayout` object is added to an other, or the RCX object already contains a `CartesianLayout` object, some attributes might be present in both. By default, the properties are updated with the values of the latest one. This can prevented by setting the *replace* parameter to `FALSE`, in that case only new properties are added and the existing properties remain untouched.

Furthermore, if duplicated properties are considered as a preventable mistake, an error can be raised by setting *stopOnDuplicates* to `TRUE`. This forces the function to stop and raise an error, if duplicated properties are present.

#### Value

*CartesianLayoutAspect* or RCX object with added layout

#### Examples

```
## For CartesianLayoutAspects:
## prepare some aspects:
cartesianLayout = createCartesianLayout(
  node=c(0, 1),
  x=c(5.5, 110.1),
  y=c(200.3, 210.2),
  z=c(-1, 3.1),
)

## node 0 is updated, new view is added
cartesianLayout2 = createCartesianLayout(
  node=c(0, 0),
  x=c(5.7, 7.2),
  y=c(98, 13.9),
  view=c(NA, 1476)
)
```

```
## Simply update with new values
cartesianLayout3 = updateCartesianLayout(cartesianLayout, cartesianLayout2)

## Ignore already present keys
cartesianLayout3 = updateCartesianLayout(cartesianLayout, cartesianLayout2,
                                         replace=FALSE)

## Raise an error if duplicate keys are present
try(updateCartesianLayout(cartesianLayout, cartesianLayout2,
                          stopOnDuplicates=TRUE))
## =>ERROR:
## Provided IDs (node, view) countain duplicates!

## For RCX:
## prepare RCX object:
nodes = createNodes(name = c("a","b"))
edges = createEdges(source = 0, target = 1)
cySubNetworks = createCySubNetworks(
  id = 1476,
  nodes = "all",
  edges = "all"
)
rcx = createRCX(nodes,
                edges = edges,
                cySubNetworks=cySubNetworks)

## add the network attributes
rcx = updateCartesianLayout(rcx, cartesianLayout)

## add additional network attributes and update existing
rcx = updateCartesianLayout(rcx, cartesianLayout2)
```

---

`updateCyGroups` *Update Cytoscape Groups*

---

#### Description

This functions add Cytoscape groups in the form of a `CyGroups` object to an RCX or an other `CyGroups` object.

#### Usage

```
updateCyGroups(x, cyGroups, stopOnDuplicates = FALSE, keepOldIds = TRUE, ...)

## S3 method for class 'CyGroupsAspect'
updateCyGroups(x, cyGroups, stopOnDuplicates = FALSE, keepOldIds = TRUE, ...)

## S3 method for class 'RCX'
updateCyGroups(
```

```
  x,
  cyGroups,
  stopOnDuplicates = FALSE,
  keepOldIds = TRUE,
  checkReferences = TRUE,
  ...
)
```

#### Arguments

| | |
|---|---|
| x | RCX or CyGroups object; (to which the new Cytoscape groups will be added) |
| cyGroups | CyGroups object; (the new aspect, that will be added) |
| stopOnDuplicates | logical; whether to stop, if duplicates in id column are found, or re-assign ids instead. |
| keepOldIds | logical; if ids are re-assigned, the original ids are kept in the column *oldId* |
| ... | additional parameters |
| checkReferences | logical; whether to check if references to other aspects are present in the RCX object |

#### Details

Cytoscape groups allow to group a set of nodes and corresponding internal and external edges together, and represent a group as a single node in the visualization. CyGroups objects can be added to an RCX or an other CyGroups object. The *nodes*, *internalEdges* and *externalEdges* parameters reference the node or edge IDs that belong to a group. When adding an CyGroups object to an RCX object, those IDs must be present in the Nodes or Edges aspect respectively, otherwise an error is raised.

When two groups should be added to each other some conflicts may rise, since the aspects might use the same IDs. If the aspects do not share any IDs, the two aspects are simply combined. Otherwise, the IDs of the new groups are re-assinged continuing with the next available ID (i.e. maxId(cyGroupsAspect) + 1 and maxId(rcx$cyGroups) + 1, respectively).

To keep track of the changes, it is possible to keep the old IDs of the newly added nodes in the automatically added column *oldId*. This can be omitted by setting *keepOldIds* to FALSE. Otherwise, if a re-assignment of the IDs is not desired, this can be prevented by setting *stopOnDuplicates* to TRUE. This forces the function to stop and raise an error, if duplicated IDs are present.

#### Value

CyGroups or RCX object with added Cytoscape groups

#### See Also

CyGroups;

#### Examples

```
## For CyGroupsAspects:
## prepare some aspects:
cyGroups1 = createCyGroups(
  name = c("Group One", "Group Two"),
  nodes = list(c(1,2,3), 0),
  internalEdges = list(c(0,1),NA),
  externalEdges = list(NA,c(2,3)),
  collapsed = c(TRUE,NA)
)

cyGroups2 = createCyGroups(
  name = "Group Three",
  nodes = list(c(4,5)),
  externalEdges = list(c(4,5))
)

## group ids will be kept
cyGroups3 = updateCyGroups(cyGroups1, cyGroups2)

## old group ids will be omitted
cyGroups3 = updateCyGroups(cyGroups1, cyGroups2,
                           keepOldIds=FALSE)

## Raise an error if duplicate keys are present
try(updateCyGroups(cyGroups1, cyGroups2,
                   stopOnDuplicates=TRUE))
## =>ERROR:
## Elements of "id" (in updateCyGroups) must not contain duplicates!

## For RCX
## prepare RCX object:
nodes = createNodes(name = c("a","b","c","d","e","f"))
edges = createEdges(source=c(1,2,0,0,0,2),
                    target=c(2,3,1,2,5,4))
rcx = createRCX(nodes, edges)

## add the group
rcx = updateCyGroups(rcx, cyGroups1)

## add an additional group
rcx = updateCyGroups(rcx, cyGroups2)

## create a group with a not existing node...
cyGroups3 = createCyGroups(
  name = "Group Three",
  nodes = list(9)
)

## ...and try to add them
try(updateCyGroups(rcx, cyGroups3))
## =>ERROR:
```

```
## Provided IDs of "additionalGroups$nodes" (in updateCyGroups)
## don't exist in "rcx$nodes$id"

## create a group with a not existing edge...
cyGroups4 = createCyGroups(
  name = "Group Four",
  nodes = list(c(1,2)),
  internalEdges = list(c(9))
)

## ...and try to add them
try(updateCyGroups(rcx, cyGroups4))
## =>ERROR:
## Provided IDs of "additionalGroups$internalEdges" (in updateCyGroups)
## don't exist in "rcx$edges$id"
```

---

updateCyHiddenAttributes

*Update Cytoscape hidden attributes*

---

#### Description

This functions add hidden attributes in the form of a CyHiddenAttributes object to an other CyHiddenAttributes or an RCX object.

#### Usage

```
updateCyHiddenAttributes(
  x,
  hiddenAttributes,
  replace = TRUE,
  stopOnDuplicates = FALSE,
  ...
)

## S3 method for class 'CyHiddenAttributesAspect'
updateCyHiddenAttributes(
  x,
  hiddenAttributes,
  replace = TRUE,
  stopOnDuplicates = FALSE,
  ...
)

## S3 method for class 'RCX'
updateCyHiddenAttributes(
  x,
  hiddenAttributes,
```

```
  replace = TRUE,
  stopOnDuplicates = FALSE,
  checkReferences = TRUE,
  ...
)
```

#### Arguments

- `x` — RCX or `CyHiddenAttributes` object; (to which the new hidden attributes will be added)
- `hiddenAttributes` — `CyHiddenAttributes` object; (the new aspect, that will be added)
- `replace` — logical; if existing values are updated (or ignored)
- `stopOnDuplicates` — logical; whether to stop, if duplicates in `name` (and `subnetworkId` if present) column are found
- `...` — additional parameters
- `checkReferences` — logical; whether to check if references to other aspects are present in the RCX object

#### Details

Cytoscape subnetworks allow to group a set of nodes and corresponding edges together, and network relations define the relations between those networks. `CyHiddenAttributes` objects can be added to an RCX or an other `CyHiddenAttributes` object.

In the case, that a `CyHiddenAttributes` object is added to an other, or the RCX object already contains a `CyHiddenAttributes` object, some attributes might be present in both. By default, the attributes are updated with the values of the latest one. This can prevented by setting the *replace* parameter to `FALSE`, in that case only new attributes are added and the existing attributes remain untouched.

Furthermore, if duplicated attributes are considered as a preventable mistake, an error can be raised by setting *stopOnDuplicates* to `TRUE`. This forces the function to stop and raise an error, if duplicated attributes are present.

#### Value

`CyHiddenAttributes` or RCX object with added hidden attributes

#### Examples

```
## For CyHiddenAttributesAspects:
## prepare some aspects:
hiddenAttributes1 = createCyHiddenAttributes(
  name=c("A","A","B","B"),
  value=list(c("a1","a2"),
             "a with subnetwork",
             "b",
```

```
              "b with subnetwork"),
  isList=c(TRUE,FALSE,TRUE,FALSE),
  subnetworkId=c(NA,1,NA,1)
)

## A is updated, C is new
hiddenAttributes2 = createCyHiddenAttributes(
  name=c("A","A","C"),
  value=list("new a",
              "new a with subnetwork",
              c(1,2)),
  subnetworkId=c(NA,1,NA)
)

## Simply update with new values
hiddenAttributes3 = updateCyHiddenAttributes(hiddenAttributes1,
                                              hiddenAttributes2)

## Ignore already present keys
hiddenAttributes3 = updateCyHiddenAttributes(hiddenAttributes1,
                                              hiddenAttributes2,
                                              replace=FALSE)

## Raise an error if duplicate keys are present
try(updateCyHiddenAttributes(hiddenAttributes1, hiddenAttributes2,
                              stopOnDuplicates=TRUE))
## =>ERROR:
## Elements of "name" and "subnetworkId" (in updateCyHiddenAttributes)
## must not contain duplicates!

## For RCX
## prepare RCX object:
nodes = createNodes(name = c("a","b","c","d","e","f"))
edges = createEdges(source=c(1,2,0,0,0,2),
                    target=c(2,3,1,2,5,4))
rcx = createRCX(nodes, edges)
cySubNetworks = createCySubNetworks(
  id = c(1,2),
  nodes = list("all", c(1,2,3)),
  edges = list("all", c(0,2))
)
rcx = updateCySubNetworks(rcx, cySubNetworks)

## add a network relation
rcx = updateCyHiddenAttributes(rcx, hiddenAttributes1)

## add an additional relation (update with new values)
rcx = updateCyHiddenAttributes(rcx, hiddenAttributes2)

## create a relation with a not existing subnetwork...
hiddenAttributes3 = createCyHiddenAttributes(
  name="X",
  value="new x",
```

```
  subnetworkId=9
)

## ...and try to add them
try(updateCyHiddenAttributes(rcx, hiddenAttributes3))
## =>ERROR:
## Provided IDs of "additionalAttributes$subnetworkId" (in updateCyHiddenAttributes)
## don't exist in "rcx$cySubNetworks$id"
```

---

updateCyNetworkRelations
*Update Cytoscape network relations*

---

#### Description

This functions add network relations in the form of a CyNetworkRelations object to an other CyNetworkRelations or an RCX object.

#### Usage

```
updateCyNetworkRelations(
  x,
  cyNetworkRelations,
  replace = TRUE,
  stopOnDuplicates = FALSE,
  ...
)

## S3 method for class 'CyNetworkRelationsAspect'
updateCyNetworkRelations(
  x,
  cyNetworkRelations,
  replace = TRUE,
  stopOnDuplicates = FALSE,
  ...
)

## S3 method for class 'RCX'
updateCyNetworkRelations(
  x,
  cyNetworkRelations,
  replace = TRUE,
  stopOnDuplicates = FALSE,
  checkReferences = TRUE,
  ...
)
```

#### Arguments

- `x` — RCX or `CySubNetworks` object; (to which the new network relations will be added)
- `cyNetworkRelations` — `CySubNetworks` object; (the network relations, that will be added)
- `replace` — logical; if existing values are updated (or ignored)
- `stopOnDuplicates` — logical; whether to stop, if duplicates in the *child* column are found
- `...` — additional parameters
- `checkReferences` — logical; whether to check if references to other aspects are present in the RCX object

#### Details

Cytoscape subnetworks allow to group a set of nodes and corresponding edges together, and network relations define the relations between those networks. `CyNetworkRelations` objects can be added to an RCX or an other `CyNetworkRelations` object.

When network relations are added to a `CyNetworkRelations` or a RCX object some conflicts may rise, since the aspects might use the same child IDs. If the aspects do not share any child IDs, the two aspects are simply combined, otherwise, the properties of the child are updated. If that is not wanted, the updating can be prevented by setting *replace* to `FALSE`. Furthermore, if duplicated properties are considered as a preventable mistake, an error can be raised by setting *stopOnDuplicates* to `TRUE`. This forces the function to stop and raise an error, if duplicated child IDs are present.

#### Value

`CyNetworkRelations` or RCX object with added network relations

#### Examples

```
## For CyNetworkRelationsAspects:
## prepare some aspects:
cyNetworkRelations1 = createCyNetworkRelations(
  child = c(1,2),
  parent = c(NA,1),
  name = c("Network A",
           "View A"),
  isView = c(FALSE, TRUE)
)

cyNetworkRelations2 = createCyNetworkRelations(
  child = 2,
  name = "View B",
  isView = TRUE
)

## update the duplicated child
cyNetworkRelations3 = updateCyNetworkRelations(cyNetworkRelations1,
```

```
                                              cyNetworkRelations2)

## keep old child values
cyNetworkRelations3 = updateCyNetworkRelations(cyNetworkRelations1,
                                              cyNetworkRelations2,
                                              replace=FALSE)

## Raise an error if duplicate keys are present
try(updateCyNetworkRelations(cyNetworkRelations1,
                             cyNetworkRelations2,
                             stopOnDuplicates=TRUE))
## =>ERROR:
## Elements of "child" (in updateCyNetworkRelations)
## must not contain duplicates!

## For RCX
## prepare RCX object:
nodes = createNodes(name = c("a","b","c","d","e","f"))
edges = createEdges(source=c(1,2,0,0,0,2),
                    target=c(2,3,1,2,5,4))
rcx = createRCX(nodes, edges)
cySubNetworks = createCySubNetworks(
  id = c(1,2),
  nodes = list("all", c(1,2,3)),
  edges = list("all", c(0,2))
)
rcx = updateCySubNetworks(rcx, cySubNetworks)

## add a network relation
rcx = updateCyNetworkRelations(rcx, cyNetworkRelations1)

## add an additional relation (View A is replaced by B)
rcx = updateCyNetworkRelations(rcx, cyNetworkRelations2)

## create a relation with a not existing subnetwork...
cyNetworkRelations3 = createCyNetworkRelations(
  child = 9
)

## ...and try to add them
try(updateCyNetworkRelations(rcx, cyNetworkRelations3))
## =>ERROR:
## Provided IDs of "additionalNetworkRelations$child" (in addCyNetworkRelations)
## don't exist in "rcx$cySubNetworks$id"

## create a relation with a not existing parent subnetwork...
cyNetworkRelations4 = createCyNetworkRelations(
  child = 1,
  parent = 9
)

## ...and try to add them
try(updateCyNetworkRelations(rcx, cyNetworkRelations4))
```

```
## =>ERROR:
## Provided IDs of "additionalNetworkRelations$parent" (in addCyNetworkRelations)
## don't exist in "rcx$cySubNetworks$id"
```

updateCySubNetworks *Update Cytoscape subnetworks*

#### Description

This functions add subnetworks in the form of a CySubNetworks object to an other CySubNetworks or an RCX object.

#### Usage

```
updateCySubNetworks(
  x,
  cySubNetworks,
  stopOnDuplicates = FALSE,
  keepOldIds = TRUE,
  ...
)

## S3 method for class 'CySubNetworksAspect'
updateCySubNetworks(
  x,
  cySubNetworks,
  stopOnDuplicates = FALSE,
  keepOldIds = TRUE,
  ...
)

## S3 method for class 'RCX'
updateCySubNetworks(
  x,
  cySubNetworks,
  stopOnDuplicates = FALSE,
  keepOldIds = TRUE,
  checkReferences = TRUE,
  ...
)
```

#### Arguments

| | |
|---|---|
| x | RCX or CySubNetworks object; (to which the new subnetworks will be added) |
| cySubNetworks | CySubNetworks object; (the subnetwork, that will be added) |
| stopOnDuplicates | logical; whether to stop, if duplicates in *id* column are found, or re-assign ids instead. |

| | |
|---|---|
| keepOldIds | logical; if ids are re-assigned, the original ids are kept in the column *oldId* |
| ... | additional parameters |
| checkReferences | logical; whether to check if references to other aspects are present in the RCX object |

#### Details

Cytoscape subnetworks allow to group a set of nodes and corresponding edges together. `CySubNetworks` objects can be added to an RCX or an other `CySubNetworks` object. The *nodes* and *edges* parameters reference the node or edge IDs that belong to a subnetwork. When adding an `CySubNetworks` object to an RCX object, those IDs must be present in the `Nodes` or `Edges` aspect respectively, otherwise an error is raised. Unlike other aspects referring those IDs, the Cytoscape subnetwork aspect allows to refer to all nodes and edges using the keyword `all`.

When subnetworks should be added to a `CySubNetworks` or a RCX object some conflicts may rise, since the aspects might use the same IDs. If the aspects do not share any IDs, the two aspects are simply combined. Otherwise, the IDs of the new subnetworks are re-assinged continuing with the next available ID (i.e. `maxId`(cySubNetworks) + 1 and `maxId`(rcx$cySubNetworks) + 1, respectively).

To keep track of the changes, it is possible to keep the old IDs of the newly added nodes in the automatically added column *oldId*. This can be omitted by setting *keepOldIds* to `FALSE`. Otherwise, if a re-assignment of the IDs is not desired, this can be prevented by setting *stopOnDuplicates* to `TRUE`. This forces the function to stop and raise an error, if duplicated IDs are present.

#### Value

`CySubNetworks` or RCX object with added subnetworks

#### See Also

`CyNetworkRelations`;

#### Examples

```
## For CySubNetworksAspects:
## prepare some aspects:
cySubNetworks1 = createCySubNetworks(
  id = c(0,1),
  nodes = list("all",
               c(1,2,3)),
  edges = list("all",
               c(0,2))
)

cySubNetworks2 = createCySubNetworks(
  nodes = c(0,3),
  edges = c(1)
)

## subnetwork ids will be kept
```

```
cySubNetworks3 = updateCySubNetworks(cySubNetworks1, cySubNetworks2)

## old subnetwork ids will be omitted
cySubNetworks3 = updateCySubNetworks(cySubNetworks1, cySubNetworks2,
                                     keepOldIds=FALSE)

## Raise an error if duplicate keys are present
try(updateCySubNetworks(cySubNetworks1, cySubNetworks2,
                        stopOnDuplicates=TRUE))
## =>ERROR:
## Elements of "id" (in updateCySubNetworks) must not contain duplicates!

## For RCX
## prepare RCX object:
nodes = createNodes(name = c("a","b","c","d","e","f"))
edges = createEdges(source=c(1,2,0,0,0,2),
                    target=c(2,3,1,2,5,4))
rcx = createRCX(nodes, edges)

## add the subnetwork
rcx = updateCySubNetworks(rcx, cySubNetworks1)

## add additional subnetwork
rcx = updateCySubNetworks(rcx, cySubNetworks2)

## create a subnetwork with a not existing node...
cySubNetworks3 = createCySubNetworks(
  nodes = list(9)
)

## ...and try to add them
try(updateCySubNetworks(rcx, cySubNetworks3))
## =>ERROR:
## Provided IDs of "additionalSubNetworks$nodes" (in addCySubNetworks)
## don't exist in "rcx$nodes$id"

## create a group with a not existing edge...
cySubNetworks4 = createCySubNetworks(
  nodes = c(0,1),
  edges = 9
)

## ...and try to add them
try(updateCySubNetworks(rcx, cySubNetworks4))
## =>ERROR:
## Provided IDs of "additionalSubNetworks$edges" (in addCySubNetworks)
## don't exist in "rcx$edges$id"
```

---

#### updateCyTableColumn *Update Cytoscape table column properties*

---

#### Description

This functions add hidden attributes in the form of a CyTableColumn object to an other CyTableColumn or an RCX object.

#### Usage

```
updateCyTableColumn(
  x,
  cyTableColumns,
  replace = TRUE,
  stopOnDuplicates = FALSE,
  ...
)

## S3 method for class 'CyTableColumnAspect'
updateCyTableColumn(
  x,
  cyTableColumns,
  replace = TRUE,
  stopOnDuplicates = FALSE,
  ...
)

## S3 method for class 'RCX'
updateCyTableColumn(
  x,
  cyTableColumns,
  replace = TRUE,
  stopOnDuplicates = FALSE,
  checkReferences = TRUE,
  ...
)
```

#### Arguments

| | |
|---|---|
| x | RCX or CyTableColumn object; (to which the new table column properties will be added) |
| cyTableColumns | CyTableColumn object; (the new aspect, that will be added) |
| replace | logical; if existing values are updated (or ignored) |
| stopOnDuplicates | logical; whether to stop, if duplicates in *appliesTo* and *name*‘ (and *subnetworkId* if present) column are found |
| ... | additional parameters |
| checkReferences | logical; whether to check if references to other aspects are present in the RCX object |

#### Details

In the case, that a CyTableColumn object is added to an other, or the RCX object already contains a CyTableColumn object, some properties might be present in both. By default, the properties are updated with the values of the latest one. This can prevented by setting the *replace* parameter to FALSE, in that case only new attributes are added and the existing attributes remain untouched.

Furthermore, if duplicated properties are considered as a preventable mistake, an error can be raised by setting *stopOnDuplicates* to TRUE. This forces the function to stop and raise an error, if duplicated properties are present.

Cytoscape does not currently support table columns for the root network, but this is option is included here for consistency.

#### Value

CyTableColumn or RCX object with added hidden attributes

#### See Also

CySubNetworks

#### Examples

```
## For CyTableColumnssAspects:
## prepare some aspects:
tableColumn1 = createCyTableColumn(
  appliesTo=c("nodes","edges","networks"),
  name=c("weight","weight","collapsed"),
  dataType=c("numeric","double","logical"),
  isList=c(FALSE,FALSE,TRUE),
  subnetworkId=c(NA,NA,1)
)

## nodes is updated, networks is new
tableColumn2 = createCyTableColumn(
  appliesTo=c("nodes","networks"),
  name=c("weight","collapsed"),
  dataType=c("double","character")
)

## Simply update with new values
tableColumn3 = updateCyTableColumn(tableColumn1, tableColumn2)

## Ignore already present keys
tableColumn3 = updateCyTableColumn(tableColumn1, tableColumn2,
                                  replace=FALSE)

## Raise an error if duplicate keys are present
try(updateCyTableColumn(tableColumn1, tableColumn2,
                       stopOnDuplicates=TRUE))
## =>ERROR:
## Elements of "appliesTo", "name" and "subnetworkId" (in updateCyTableColumn)
```

```
## must not contain duplicates!

## For RCX
## prepare RCX object:
nodes = createNodes(name = c("a","b","c","d","e","f"))
edges = createEdges(source=c(1,2,0,0,0,2),
                    target=c(2,3,1,2,5,4))
rcx = createRCX(nodes, edges)
cySubNetworks = createCySubNetworks(
  id = c(1,2),
  nodes = list("all", c(1,2,3)),
  edges = list("all", c(0,2))
)
rcx = updateCySubNetworks(rcx, cySubNetworks)

## add a table column property
rcx = updateCyTableColumn(rcx, tableColumn1)

## add an additional property (update with new values)
rcx = updateCyTableColumn(rcx, tableColumn2)

## create a prpperty with a not existing subnetwork...
tableColumn3 = createCyTableColumn(
  appliesTo="nodes",
  name="weight",
  subnetworkId=9
)

## ...and try to add them
try(updateCyTableColumn(rcx, tableColumn3))
## =>ERROR:
## Provided IDs of "additionalColumns$subnetworkId" (in addCyTableColumn)
## don't exist in "rcx$cySubNetworks$id"
```

---

updateCyVisualProperties *Update Cytoscape Visual Properties (aspect)*

---

#### Description

This function is used to add Cytoscape visual properties aspects to each other or to an RCX object. In a `CyVisualProperties` aspect, `CyVisualProperty` objects define networks, nodes, edges, and default nodes and edges.

#### Usage

```
updateCyVisualProperties(
  x,
  cyVisualProperties,
  replace = TRUE,
```

```
  stopOnDuplicates = FALSE,
  ...
)

## S3 method for class 'CyVisualPropertiesAspect'
updateCyVisualProperties(
  x,
  cyVisualProperties,
  replace = TRUE,
  stopOnDuplicates = FALSE,
  ...
)

## S3 method for class 'RCX'
updateCyVisualProperties(
  x,
  cyVisualProperties,
  replace = TRUE,
  stopOnDuplicates = FALSE,
  checkReferences = TRUE,
  ...
)
```

#### Arguments

| | |
|---|---|
| x | RCX or CyVisualProperties object; (to which it will be added) |
| cyVisualProperties | CyVisualProperties object; (that will be added) |
| replace | logical; if existing values are updated (or ignored) |
| stopOnDuplicates | logical; whether to stop, if duplicates in name (and subnetworkId if present) column are found |
| ... | additional parameters |
| checkReferences | logical; whether to check if references to other aspects are present in the RCX object |

#### Details

##### Structure of Cytoscape Visual Property:

```
CyVisualProperty
|---properties = CyVisualPropertyProperties
|   |--name
|   |--value
|---dependencies = CyVisualPropertyDependencies
|   |--name
|   |--value
```

```
|---mappings = CyVisualPropertyMappings
|    |--name
|    |--type
|    |--definition
|---appliesTo = <reference to subnetwork id>
|---view = <reference to subnetwork id>
```

CyVisualProperties aspects consist of CyVisualProperty objects for each entry: networks, nodes, edges, and default nodes and edges. Two CyVisualProperties aspects are merged by adding its entries individually.

CyVisualProperty objects differ in the sub-networks and views (CySubNetworks) they apply to, subsequently properties, dependencies and mappings are merged based on the uniqueness in those two.

Properties, dependencies and mappings (i.e. CyVisualPropertyProperties, CyVisualPropertyDependencies and CyVisualPropertyMappings objects) are unique in name. By default, the duplicate attributes are updated with the values of the latest one. This can prevented by setting the *replace* parameter to FALSE, in that case only new attributes are added and the existing attributes remain untouched. Furthermore, if duplicated attributes are considered as a preventable mistake, an error can be raised by setting *stopOnDuplicates* to TRUE. This forces the function to stop and raise an error, if duplicated attributes are present.

#### Value

CyVisualProperties or RCX object with added Cytoscape visual properties

#### See Also

updateCyVisualProperty, getCyVisualProperty

#### Examples

```
## Prepare used properties
## Visual property: Properties
vpPropertyP1 = createCyVisualPropertyProperties(c(NODE_BORDER_STROKE="SOLID"))
vpPropertyP2 = createCyVisualPropertyProperties(c(NODE_BORDER_WIDTH="1.5"))
vpPropertyP3 = createCyVisualPropertyProperties(c(NODE_BORDER_WIDTH="999"))

## Visual property: Dependencies
vpPropertyD1 = createCyVisualPropertyDependencies(c(nodeSizeLocked="false"))
vpPropertyD2 = createCyVisualPropertyDependencies(c(arrowColorMatchesEdge="true"))
vpPropertyD3 = createCyVisualPropertyDependencies(c(arrowColorMatchesEdge="false"))

## Visual property: Mappings
vpPropertyM1 = createCyVisualPropertyMappings(c(NODE_FILL_COLOR="CONTINUOUS"),
                                              "COL=directed,T=boolean,K=0=true,V=0=ARROW")
vpPropertyM2 = createCyVisualPropertyMappings(c(EDGE_TARGET_ARROW_SHAPE="DISCRETE"),
                                                "TRIANGLE")
vpPropertyM3 = createCyVisualPropertyMappings(c(EDGE_TARGET_ARROW_SHAPE="DISCRETE"),
                                                "NONE")

## Create visual property object
```

```
vpProperty1 = createCyVisualProperty(properties=list(vpPropertyP1,
                                                     vpPropertyP1),
                                     dependencies=list(vpPropertyD1,
                                                       NA),
                                     mappings=list(vpPropertyM1,
                                                   NA),
                                     appliesTo = c(NA,
                                                   1),
                                     view = c(NA,
                                              1))
vpProperty2 = createCyVisualProperty(properties=vpPropertyP2,
                                     dependencies=vpPropertyD2,
                                     mappings=vpPropertyM2)
vpProperty3 = createCyVisualProperty(properties=vpPropertyP3,
                                     dependencies=vpPropertyD3,
                                     mappings=vpPropertyM3)

## Create a visual properties aspect
## (using the same visual property object for simplicity)
visProp1 = createCyVisualProperties(network=vpProperty1,
                                    nodes=vpProperty1,
                                    edges=vpProperty1,
                                    defaultNodes=vpProperty1,
                                    defaultEdges=vpProperty1)

visProp2 = createCyVisualProperties(network=vpProperty2,
                                    nodes=vpProperty2,
                                    edges=vpProperty2,
                                    defaultNodes=vpProperty2,
                                    defaultEdges=vpProperty2)

visProp3 = createCyVisualProperties(network=vpProperty3,
                                    nodes=vpProperty3,
                                    edges=vpProperty3,
                                    defaultNodes=vpProperty3,
                                    defaultEdges=vpProperty3)

## Adding a different visual property (Properties, Dependencies, Mappings)
## (e.g. "NODE_BORDER_WIDTH", which is not present before)
visProp4 = updateCyVisualProperties(visProp1, visProp2)

## Update a existing visual property
visProp5 = updateCyVisualProperties(visProp4, visProp3)

## Raise an error if duplicate keys are present
try(updateCyVisualProperties(visProp4, visProp3,
                             stopOnDuplicates=TRUE))
## =>ERROR:
##   Elements of name (in VisualProperties$network$properties<appliesTo=NA,view=NA>)
##   must not contain duplicates!

## For RCX
## prepare RCX object:
```

```
nodes = createNodes(name = c("a","b","c","d","e","f"))
edges = createEdges(source=c(1,2,0,0,0,2),
                    target=c(2,3,1,2,5,4))
rcx = createRCX(nodes, edges)
cySubNetworks = createCySubNetworks(
  id = c(1,2),
  nodes = list("all", c(1,2,3)),
  edges = list("all", c(0,2))
)
rcx = updateCySubNetworks(rcx, cySubNetworks)

## Adding visual properties to an RCX object
rcx = updateCyVisualProperties(rcx, visProp1)

## Adding a different visual property (Properties, Dependencies, Mappings)
## (e.g. "NODE_BORDER_WIDTH", which is not present before)
rcx = updateCyVisualProperties(rcx, visProp2)

## Update a existing visual property
rcx = updateCyVisualProperties(rcx, visProp3)

## Raise an error if duplicate keys are present
try(updateCyVisualProperties(rcx, visProp3,
                             stopOnDuplicates=TRUE))
## =>ERROR:
## Elements of "name" (in VisualProperties$network$properties<appliesTo=NA,view=NA>)
## must not contain duplicates!
```

---

updateCyVisualProperty *Update Cytoscape Visual Property objects and sub-objects (used in CyVisualProperties aspect)*

---

#### Description

This function is used to add Cytoscape visual property objects (CyVisualProperty) and its sub-objects (CyVisualPropertyProperties, CyVisualPropertyDependencies and CyVisualPropertyMappings) to each other. Cytoscape visual property objects define networks, nodes, edges, and default nodes and edges in a CyVisualProperties aspect.

#### Usage

```
updateCyVisualProperty(
  cyVisualProperty,
  additionalProperty,
  replace = TRUE,
  stopOnDuplicates = FALSE,
  .log = c()
)
```

```
## S3 method for class 'CyVisualPropertyProperties'
updateCyVisualProperty(
  cyVisualProperty,
  additionalProperty,
  replace = TRUE,
  stopOnDuplicates = FALSE,
  .log = c()
)

## S3 method for class 'CyVisualPropertyDependencies'
updateCyVisualProperty(
  cyVisualProperty,
  additionalProperty,
  replace = TRUE,
  stopOnDuplicates = FALSE,
  .log = c()
)

## S3 method for class 'CyVisualPropertyMappings'
updateCyVisualProperty(
  cyVisualProperty,
  additionalProperty,
  replace = TRUE,
  stopOnDuplicates = FALSE,
  .log = c()
)

## S3 method for class 'CyVisualProperty'
updateCyVisualProperty(
  cyVisualProperty,
  additionalProperty,
  replace = TRUE,
  stopOnDuplicates = FALSE,
  .log = c()
)
```

#### Arguments

`cyVisualProperty`
: object; (to which it will be added)

`additionalProperty`
: object; (that will be added)

`replace`
: logical; if existing values are updated (or ignored)

`stopOnDuplicates`
: logical; whether to stop, if duplicates in `name` (and `subnetworkId` if present) column are found

`.log`
: character (optional); name of the calling function used in error logging

#### Details

**Structure of Cytoscape Visual Property:**

```
CyVisualProperty
|---properties = CyVisualPropertyProperties
|   |--name
|   |--value
|---dependencies = CyVisualPropertyDependencies
|   |--name
|   |--value
|---mappings = CyVisualPropertyMappings
|   |--name
|   |--type
|   |--definition
|---appliesTo = <reference to subnetwork id>
|---view = <reference to subnetwork id>
```

CyVisualProperty objects differ in the sub-networks and views (CySubNetworks) they apply to, subsequently properties, dependencies and mappings are merged based on the uniqueness in those two.

Properties, dependencies and mappings (i.e. CyVisualPropertyProperties, CyVisualPropertyDependencies and CyVisualPropertyMappings objects) are unique in name. By default, the duplicate attributes are updated with the values of the latest one. This can prevented by setting the *replace* parameter to FALSE, in that case only new attributes are added and the existing attributes remain untouched. Furthermore, if duplicated attributes are considered as a preventable mistake, an error can be raised by setting *stopOnDuplicates* to TRUE. This forces the function to stop and raise an error, if duplicated attributes are present.

#### Value

CyVisualProperty, CyVisualPropertyProperties, CyVisualPropertyDependencies or CyVisualPropertyMappings objects

#### See Also

getCyVisualProperty, updateCyVisualProperties

#### Examples

```
## Prepare used properties
## Visual property: Properties
vpPropertyP1 = createCyVisualPropertyProperties(c(NODE_BORDER_STROKE="SOLID"))
vpPropertyP2 = createCyVisualPropertyProperties(c(NODE_BORDER_WIDTH="1.5"))
vpPropertyP3 = createCyVisualPropertyProperties(c(NODE_BORDER_WIDTH="999"))

## Add two properties:
vpPropertyP4 = updateCyVisualProperty(vpPropertyP1,vpPropertyP2)
vpPropertyP4 = updateCyVisualProperty(vpPropertyP4,vpPropertyP3)

## Visual property: Dependencies
```

```
vpPropertyD1 = createCyVisualPropertyDependencies(c(nodeSizeLocked="false"))
vpPropertyD2 = createCyVisualPropertyDependencies(c(arrowColorMatchesEdge="true"))
vpPropertyD3 = createCyVisualPropertyDependencies(c(arrowColorMatchesEdge="false"))

## Add two dependencies:
vpPropertyD4 = updateCyVisualProperty(vpPropertyD1,vpPropertyD2)
vpPropertyD4 = updateCyVisualProperty(vpPropertyD4,vpPropertyD3)

## Visual property: Mappings
vpPropertyM1 = createCyVisualPropertyMappings(c(NODE_FILL_COLOR="CONTINUOUS"),
                                            "COL=directed,T=boolean,K=0=true,V=0=ARROW")
vpPropertyM2 = createCyVisualPropertyMappings(c(EDGE_TARGET_ARROW_SHAPE="DISCRETE"),
                                              "TRIANGLE")
vpPropertyM3 = createCyVisualPropertyMappings(c(EDGE_TARGET_ARROW_SHAPE="DISCRETE"),
                                              "NONE")

## Add two mappings:
vpPropertyM4 = updateCyVisualProperty(vpPropertyM1,vpPropertyM2)
vpPropertyM4 = updateCyVisualProperty(vpPropertyM4,vpPropertyM3)

## Create visual property object
vpProperty1 = createCyVisualProperty(properties=list(vpPropertyP1,
                                                     vpPropertyP1,
                                                     vpPropertyP1),
                                     dependencies=list(vpPropertyD1,
                                                       vpPropertyD1,
                                                       NA),
                                     mappings=list(vpPropertyM1,
                                                   NA,
                                                   vpPropertyM1),
                                     appliesTo = c(NA,
                                                   NA,
                                                   1),
                                     view = c(NA,
                                              1,
                                              NA))
vpProperty2 = createCyVisualProperty(properties=vpPropertyP2,
                                     dependencies=vpPropertyD2,
                                     mappings=vpPropertyM2)
vpProperty3 = createCyVisualProperty(properties=vpPropertyP3,
                                     dependencies=vpPropertyD3,
                                     mappings=vpPropertyM3)

## add two visual property objects
vpProperty4 = updateCyVisualProperty(vpProperty1, vpProperty2)

## update values
updateCyVisualProperty(vpProperty4, vpProperty3)

## keep old values
updateCyVisualProperty(vpProperty4, vpProperty3,
                       replace = FALSE)
```

```
## keep old values
try(updateCyVisualProperty(vpProperty4, vpProperty3,
                           stopOnDuplicates = TRUE))
## =>ERROR:
## Elements of name (in properties<appliesTo=NA,view=NA>) must not contain duplicates!
```

---

updateEdgeAttributes *Update edge attributes*

---

#### Description

This functions add edge attributes in the form of a EdgeAttributes object to an RCX or an other EdgeAttributes object.

#### Usage

```
updateEdgeAttributes(
  x,
  edgeAttributes,
  replace = TRUE,
  stopOnDuplicates = FALSE,
  ...
)

## S3 method for class 'EdgeAttributesAspect'
updateEdgeAttributes(
  x,
  edgeAttributes,
  replace = TRUE,
  stopOnDuplicates = FALSE,
  ...
)

## S3 method for class 'RCX'
updateEdgeAttributes(
  x,
  edgeAttributes,
  replace = TRUE,
  stopOnDuplicates = FALSE,
  checkReferences = TRUE,
  ...
)
```

#### Arguments

| | |
|---|---|
| x | RCX or EdgeAttributes object; (to which the new edge attributes will be added) |

| | |
|---|---|
| edgeAttributes | EdgeAttributes object; (the new aspect, that will be added) |
| replace | logical; if existing values are updated (or ignored) |
| stopOnDuplicates | logical; whether to stop, if duplicates in *propertyOf* and *name* (and *subnetworkId* if present) column are found |
| ... | additional parameters |
| checkReferences | logical; whether to check if references to other aspects are present in the RCX object |

#### Details

Edges may have additional attributes besides a name and a representation, and are represented as EdgeAttributes objects. EdgeAttributes objects can be added to an RCX or an other EdgeAttributes object. The *propertyOf* parameter references the Edges ids to which the attributes belong to. When adding an EdgeAttributes object to an RCX object, those ids must be present in the Edges aspect, otherwise an error is raised.

In the case, that a EdgeAttributes object is added to an other, or the RCX object already contains a EdgeAttributes object, some attributes might be present in both. By default, the attributes are updated with the values of the latest one. This can prevented by setting the *replace* parameter to *FALSE*, in that case only new attributes are added and the existing attributes remain untouched.

Furthermore, if duplicated attributes are considered as a preventable mistake, an error can be raised by setting *stopOnDuplicates* to TRUE. This forces the function to stop and raise an error, if duplicated attributes are present.

#### Value

EdgeAttributes or RCX object with added node attributes

#### See Also

NodeAttributes, NetworkAttributes

#### Examples

```
## For EdgeAttributesAspects:
## prepare some aspects:
edgeAttributes = createEdgeAttributes(
  propertyOf=c(0,0,0,0),
  name=c("A","A","B","B"),
  value=list(c("a1","a2"),
             "a with subnetwork",
             "b",
             "b with subnetwork"),
  isList=c(TRUE,FALSE,TRUE,FALSE),
  subnetworkId=c(NA,1,NA,1)
)

## A is updated, C is new
```

```
edgeAttributes2 = createEdgeAttributes(
  propertyOf=c(0,0,0),
  name=c("A","A","C"),
  value=list("new a",
             "new a with subnetwork",
             c(1,2)),
  subnetworkId=c(NA,1,NA)
)

## Simply update with new values
edgeAttributes3 = updateEdgeAttributes(edgeAttributes, edgeAttributes2)

## Ignore already present keys
edgeAttributes3 = updateEdgeAttributes(edgeAttributes, edgeAttributes2,
                                       replace=FALSE)

## Raise an error if duplicate keys are present
try(updateEdgeAttributes(edgeAttributes, edgeAttributes2,
                         stopOnDuplicates=TRUE))
## =>ERROR:
## Elements of "propertyOf", "name" and "subnetworkId" (in updateEdgeAttributes)
## must not contain duplicates!

## For RCX
## prepare RCX object:
nodes = createNodes(name = c("a","b","c","d","e","f"))
edges = createEdges(source=c(1,2,0,0,0,2),
                    target=c(2,3,1,2,5,4))
rcx = createRCX(nodes, edges)
cySubNetworks = createCySubNetworks(
  id = c(1,2),
  nodes = list("all", c(1,2,3)),
  edges = list("all", c(0,2))
)
rcx = updateCySubNetworks(rcx, cySubNetworks)

## add the edge attributes
rcx = updateEdgeAttributes(rcx, edgeAttributes)

## add additional edge attributes and update existing
rcx = updateEdgeAttributes(rcx, edgeAttributes2)

## create edge attributes for a not existing edge...
edgeAttributes3 = createEdgeAttributes(propertyOf=9,
                                       name="A",
                                       value="a")
## ...and try to add them
try(updateEdgeAttributes(rcx, edgeAttributes3))
## =>ERROR:
## Provided IDs of "additionalAttributes$propertyOf" (in updateEdgeAttributes)
## don't exist in "rcx$edges$id"
```

updateEdges *Update edges*

#### Description

This functions add edges in the form of a `Edges` object to an other `Edges` or an *RCX* object.

#### Usage

```
updateEdges(x, edges, stopOnDuplicates = FALSE, keepOldIds = TRUE, ...)

## S3 method for class 'EdgesAspect'
updateEdges(x, edges, stopOnDuplicates = FALSE, keepOldIds = TRUE, ...)

## S3 method for class 'RCX'
updateEdges(
  x,
  edges,
  stopOnDuplicates = FALSE,
  keepOldIds = TRUE,
  checkReferences = TRUE,
  ...
)
```

#### Arguments

| | |
|---|---|
| `x` | `RCX-object` or `Edges` object; (to which the new `Edges` will be added) |
| `edges` | `Edges` object; (the `Edges`, that will be added) |
| `stopOnDuplicates` | logical (optional); whether to stop, if duplicates in *id* column are found, or re-assign ids instead. |
| `keepOldIds` | logical (optional); if ids are re-assigned, the original ids are kept in the column *oldId* |
| `...` | additional parameters |
| `checkReferences` | logical; whether to check if references to other aspects are present in the RCX object |

#### Details

When edges should be added to a `Edges` or a `RCX-object` object some conflicts may rise, since the aspects might use the same IDs. If the aspects do not share any IDs, the two aspects are simply combined. Otherwise, the IDs of the new edges are re-assinged continuing with the next available ID (i.e. `maxId`(edgesAspect) + 1 and `maxId`(rcx$edges) + 1, respectively).

To keep track of the changes, it is possible to keep the old IDs of the newly added edges in the automatically added column *oldId*. This can be omitted by setting *keepOldIds* to `FALSE`. Otherwise, if a re-assignment of the IDs is not desired, this can be prevented by setting *stopOnDuplicates* to TRUE. This forces the function to stop and raise an error, if duplicated IDs are present.

#### Value

Edges or RCX with added edges

#### Examples

```
## create some edges
edges1 = createEdges(source=c(1,1,0), target=c(2,0,1))
edges2 = createEdges(id=c(3,2,4),
                     source=c(0,0,1),
                     target=c(1,2,2),
                     interaction=c("activates","inhibits", NA))

## simply add the edges and keep old ids
edges3 = updateEdges(edges1, edges2)

## add the edges
edges4 = updateEdges(edges1, edges2, keepOldIds=FALSE)

## force an error because of duplicated ids
try(updateEdges(edges1, edges2, stopOnDuplicates=TRUE))
## =>Error:
## Elements of "id" (in updateEdges) must not contain duplicates!

## Prepare an RCX object
rcx = createRCX(createNodes(name = c("EGFR","AKT1","WNT")))

## add edges to the RCX object
rcx = updateEdges(rcx, edges1)

## add new edges and don't keep old ids
rcx = updateEdges(rcx, edges2, keepOldIds=FALSE)

## force an error because of duplicated ids
try(updateEdges(rcx, edges2, stopOnDuplicates=TRUE))
## =>Error:
## Elements of "id" (in updateEdges) must not contain duplicates!
```

---

### updateMetaDataProperties

*Update meta-data properties*

---

#### Description

The Meta-data aspect contains meta-data about the aspects in the RCX-object. Properties that need to be fetched or updated independently of aspect data are added with this function.

#### Usage

```
updateMetaDataProperties(rcx, aspectName, property)
```

#### Arguments

| | |
|---|---|
| rcx | RCX object; |
| aspectName | character; name of the aspect as displayed in Meta-data (e.g. "nodes") |
| property | named list; property as key-value pairs (empty list to remove all) |

#### Value

RCX object with updated Meta-data aspect

#### Examples

```
## prepare RCX object:
nodes = createNodes(name = c("a","b","c","d","e","f"))
edges = createEdges(source=c(1,2,0,0,0,2),
                    target=c(2,3,1,2,5,4))
rcx = createRCX(nodes, edges)
cySubNetworks = createCySubNetworks(
  id = c(1,2),
  nodes = list("all", c(1,2,3)),
  edges = list("all", c(0,2))
)
rcx = updateCySubNetworks(rcx, cySubNetworks)

## add properties for edges
updateMetaDataProperties(rcx,
                         "edges",
                         list(some="value",
                              another="VALUE"))

## remove properties for edges
updateMetaDataProperties(rcx,
                         "edges",
                         list())
```

---

### updateNetworkAttributes

*Update network attributes*

---

#### Description

This functions add network attributes in the form of a NetworkAttributes object to an RCX or an other NetworkAttributes object.

#### Usage

```
updateNetworkAttributes(
  x,
  networkAttributes,
  replace = TRUE,
  stopOnDuplicates = FALSE,
  ...
)

## S3 method for class 'NetworkAttributesAspect'
updateNetworkAttributes(
  x,
  networkAttributes,
  replace = TRUE,
  stopOnDuplicates = FALSE,
  ...
)

## S3 method for class 'RCX'
updateNetworkAttributes(
  x,
  networkAttributes,
  replace = TRUE,
  stopOnDuplicates = FALSE,
  checkReferences = TRUE,
  ...
)
```

#### Arguments

`x` RCX object; (to which the new network attributes will be added)

`networkAttributes` `NetworkAttributes` object; (the new aspect, that will be added)

`replace` logical; if existing values are updated (or ignored)

`stopOnDuplicates` logical; whether to stop, if duplicates in *name* (and *subnetworkId* if present) column are found

`...` additional parameters

`checkReferences` logical; whether to check if references to other aspects are present in the RCX object

#### Details

Networks may have attributes, that are represented as `NetworkAttributes` objects. `NetworkAttributes` objects can be added to an RCX or an other `NetworkAttributes` object.

In the case, that a NetworkAttributes object is added to an other, or the RCX object already contains a NetworkAttributes object, some attributes might be present in both. By default, the attributes are updated with the values of the latest one. This can prevented by setting the *replace* parameter to FALSE, in that case only new attributes are added and the existing attributes remain untouched.

Furthermore, if duplicated attributes are considered as a preventable mistake, an error can be raised by setting *stopOnDuplicates* to TRUE. This forces the function to stop and raise an error, if duplicated attributes are present.

#### Value

NetworkAttributes or RCX object with added network attributes

#### See Also

NetworkAttributes; NodeAttributes, EdgeAttributes

#### Examples

```
## For NetworkAttributesAspects:
## prepare some aspects:
networkAttributes1 = createNetworkAttributes(
  name=c("A","A","B","B"),
  value=list(c("a1","a2"),
             "a with subnetwork",
             "b",
             "b with subnetwork"),
  isList=c(TRUE,FALSE,TRUE,FALSE),
  subnetworkId=c(NA,1,NA,1)
)

## A is updated, C is new
networkAttributes2 = createNetworkAttributes(
  name=c("A","A","C"),
  value=list("new a",
             "new a with subnetwork",
             c(1,2)),
  subnetworkId=c(NA,1,NA)
)

## Simply update with new values
networkAttributes3 = updateNetworkAttributes(networkAttributes1, networkAttributes2)

## Ignore already present keys
networkAttributes3 = updateNetworkAttributes(networkAttributes1, networkAttributes2,
                                             replace=FALSE)

## Raise an error if duplicate keys are present
try(updateNetworkAttributes(networkAttributes1, networkAttributes2,
                            stopOnDuplicates=TRUE))
## =>ERROR:
## Provided IDs (name, subnetworkId) countain duplicates!
```

```
## For RCX
## prepare RCX object:
nodes = createNodes(name = c("a","b","c","d","e","f"))
edges = createEdges(source=c(1,2,0,0,0,2),
                    target=c(2,3,1,2,5,4))
rcx = createRCX(nodes, edges)
cySubNetworks = createCySubNetworks(
  id = c(1,2),
  nodes = list("all", c(1,2,3)),
  edges = list("all", c(0,2))
)
rcx = updateCySubNetworks(rcx, cySubNetworks)

## add the network attributes
rcx = updateNetworkAttributes(rcx, networkAttributes1)

## add additional network attributes and update existing
rcx = updateNetworkAttributes(rcx, networkAttributes2)

## create a relation with a not existing subnetwork...
networkAttributes3 = createNetworkAttributes(
  name="X",
  value="new x",
  subnetworkId=9
)

## ...and try to add them
try(updateNetworkAttributes(rcx, networkAttributes3))
## =>ERROR:
## NetworkAttributesAspect$subnetworkId IDs don't exist in CySubNetworksAspect
```

---

#### updateNodeAttributes *Update node attributes*

---

##### Description

This functions add node attributes in the form of a NodeAttributes object to an RCX or an other NodeAttributes object.

##### Usage

```
updateNodeAttributes(
  x,
  nodeAttributes,
  replace = TRUE,
  stopOnDuplicates = FALSE,
  ...
)
```

```
## S3 method for class 'NodeAttributesAspect'
updateNodeAttributes(
  x,
  nodeAttributes,
  replace = TRUE,
  stopOnDuplicates = FALSE,
  ...
)

## S3 method for class 'RCX'
updateNodeAttributes(
  x,
  nodeAttributes,
  replace = TRUE,
  stopOnDuplicates = FALSE,
  checkReferences = TRUE,
  ...
)
```

##### Arguments

| | |
|---|---|
| `x` | RCX or `NodeAttributes` object; (to which the new node attributes will be added) |
| `nodeAttributes` | `NodeAttributes` object; (the new aspect, that will be added) |
| `replace` | logical; if existing values are updated (or ignored) |
| `stopOnDuplicates` | logical; whether to stop, if duplicates in *propertyOf* and *name* (and *subnetworkId* if present) columns are found |
| `...` | additional parameters |
| `checkReferences` | logical; whether to check if references to other aspects are present in the RCX object |

##### Details

Nodes may have additional attributes besides a name and a representation, and are represented as `NodeAttributes` objects. `NodeAttributes` objects can be added to an RCX object or an other `NodeAttributes` object. The *propertyOf* parameter references the node IDs to which the attributes belong to. When adding an `NodeAttributes` object to an RCX object, those IDs must be present in the `Nodes` aspect, otherwise an error is raised.

In the case, that a `NodeAttributes` object is added to an other, or the RCX object already contains a `NodeAttributes` object, some attributes might be present in both. By default, the attributes are updated with the values of the latest one. This can prevented setting the *replace* parameter to `FALSE`, in that case only new attributes are added and the existing attributes remain untouched.

Furthermore, if duplicated attributes are considered as a preventable mistake, an error can be raised by setting *stopOnDuplicates* to `TRUE`. This forces the function to stop and raise an error, if duplicated attributes are present.

#### Value

NodeAttributes or RCX object with added node attributes

#### See Also

EdgeAttributes, NetworkAttributes

#### Examples

```
## For NodeAttributesAspects:
## prepare some aspects:
nodeAttributes1 = createNodeAttributes(
  propertyOf=c(1,1,1,1),
  name=c("A","A","B","B"),
  value=list(c("a1","a2"),
             "a with subnetwork",
             "b",
             "b with subnetwork"),
  isList=c(TRUE,FALSE,TRUE,FALSE),
  subnetworkId=c(NA,1,NA,1)
)

## A is updated, C is new
nodeAttributes2 = createNodeAttributes(
  propertyOf=c(1,1,1),
  name=c("A","A","C"),
  value=list("new a",
             "new a with subnetwork",
             c(1,2)),
  subnetworkId=c(NA,1,NA)
)

## Simply update with new values
nodeAttributes3 = updateNodeAttributes(nodeAttributes1, nodeAttributes2)

## Ignore already present keys
nodeAttributes4 = updateNodeAttributes(nodeAttributes1, nodeAttributes2,
                                       replace=FALSE)

## Raise an error if duplicate keys are present
try(updateNodeAttributes(nodeAttributes1, nodeAttributes2,
                         stopOnDuplicates=TRUE))
## =>ERROR:
## Elements of "propertyOf", "name" and "subnetworkId" (in addNodeAttributes)
## must not contain duplicates!

## For RCX
## prepare RCX object:
nodes = createNodes(name = c("a","b","c","d","e","f"))
edges = createEdges(source=c(1,2,0,0,0,2),
                    target=c(2,3,1,2,5,4))
rcx = createRCX(nodes, edges)
```

```
cySubNetworks = createCySubNetworks(
  id = c(1,2),
  nodes = list("all", c(1,2,3)),
  edges = list("all", c(0,2))
)
rcx = updateCySubNetworks(rcx, cySubNetworks)

## add the node attributes, even if no subnetworks are present
rcx = updateNodeAttributes(rcx, nodeAttributes1, checkReferences=FALSE)

## add the node attributes
rcx = updateNodeAttributes(rcx, nodeAttributes1)

## add additional node attributes and update existing
rcx = updateNodeAttributes(rcx, nodeAttributes2)

## create node attributes for a not existing node...
nodeAttributes3 = createNodeAttributes(propertyOf=9,
                                       name="A",
                                       value="a")
## ...and try to add them
try(updateNodeAttributes(rcx, nodeAttributes3))
## =>ERROR:
## Provided IDs of "additionalAttributes$propertyOf" (in addNodeAttributes)
## don't exist in "rcx$nodes$id"
```

---

updateNodes *Update nodes*

---

#### Description

This functions add nodes in the form of a Nodes object to an other Nodes or an RCX-object.

#### Usage

```
updateNodes(x, nodes, stopOnDuplicates = FALSE, keepOldIds = TRUE)

## S3 method for class 'NodesAspect'
updateNodes(x, nodes, stopOnDuplicates = FALSE, keepOldIds = TRUE)

## S3 method for class 'RCX'
updateNodes(x, nodes, stopOnDuplicates = FALSE, keepOldIds = TRUE)
```

#### Arguments

| | |
|---|---|
| x | RCX-object or Nodes object; (to which the new Nodes will be added) |
| nodes | Nodes object; (the Nodes, that will be added) |

stopOnDuplicates
: logical (optional); whether to stop, if duplicates in id column are found, or re-assign ids instead.

keepOldIds
: logical (optional); if ids are re-assigned, the original ids are kept in the column oldId

#### Details

When nodes should be added to a Nodes or a RCX-object object some conflicts may rise, since the aspects might use the same IDs. If the aspects do not share any IDs, the two aspects are simply combined. Otherwise, the IDs of the new nodes are re-assinged continuing with the next available ID (i.e. maxId(nodesAspect) + 1 and maxId(rcx$nodes) + 1, respectively).

To keep track of the changes, it is possible to keep the old IDs of the newly added nodes in the automatically added column *oldId*. This can be omitted by setting *keepOldIds* to FALSE. Otherwise, if a re-assignment of the IDs is not desired, this can be prevented by setting *stopOnDuplicates* to TRUE. This forces the function to stop and raise an error, if duplicated IDs are present.

#### Value

Nodes or RCX object with added nodes

#### Examples

```
## create some nodes
nodes1 = createNodes(name = c("EGFR","AKT1","WNT"))
nodes2 = createNodes(name=c("CDK1", "CDK2", "CDK3"),
                     represents=c("HGNC:CDK1",
                                  "Uniprot:P24941",
                                  "Ensembl:ENSG00000250506"))

## simply add the nodes and keep old ids
nodes3 = updateNodes(nodes1, nodes2)

## add the nodes
nodes4 = updateNodes(nodes1, nodes2, keepOldIds=FALSE)

## force an error because of duplicated ids
try(updateNodes(nodes1, nodes2, stopOnDuplicates=TRUE))
## =>Error:
## Elements of "id" (in updateNodes) must not contain duplicates!

## create an RCX object with nodes
rcx = createRCX(nodes1)

## add additional nodes
rcx = updateNodes(rcx, nodes2, keepOldIds=FALSE)

## force an error becauses of duplicated ids
try(updateNodes(rcx, nodes2, stopOnDuplicates=TRUE))
## =>Error:
## Elements of "id" (in updateNodes) must not contain duplicates!
```

validate *Validate RCX and its aspects*

#### Description

Validate RCX objects and its aspects.

#### Usage

```
validate(x, verbose = TRUE)

## Default S3 method:
validate(x, verbose = TRUE)

## S3 method for class 'NodesAspect'
validate(x, verbose = TRUE)

## S3 method for class 'EdgesAspect'
validate(x, verbose = TRUE)

## S3 method for class 'NodeAttributesAspect'
validate(x, verbose = TRUE)

## S3 method for class 'EdgeAttributesAspect'
validate(x, verbose = TRUE)

## S3 method for class 'NetworkAttributesAspect'
validate(x, verbose = TRUE)

## S3 method for class 'CartesianLayoutAspect'
validate(x, verbose = TRUE)

## S3 method for class 'CyGroupsAspect'
validate(x, verbose = TRUE)

## S3 method for class 'CyVisualPropertiesAspect'
validate(x, verbose = TRUE)

## S3 method for class 'CyVisualProperty'
validate(x, verbose = TRUE)

## S3 method for class 'CyVisualPropertyProperties'
validate(x, verbose = TRUE)

## S3 method for class 'CyVisualPropertyDependencies'
validate(x, verbose = TRUE)
```

```
## S3 method for class 'CyVisualPropertyMappings'
validate(x, verbose = TRUE)

## S3 method for class 'CyHiddenAttributesAspect'
validate(x, verbose = TRUE)

## S3 method for class 'CyNetworkRelationsAspect'
validate(x, verbose = TRUE)

## S3 method for class 'CySubNetworksAspect'
validate(x, verbose = TRUE)

## S3 method for class 'CyTableColumnAspect'
validate(x, verbose = TRUE)

## S3 method for class 'RCX'
validate(x, verbose = TRUE)
```

#### Arguments

| | |
|---|---|
| `x` | object to validate; RCX object or an aspect |
| `verbose` | logical; whether to print the test results. |

#### Details

Different tests are performed on aspects and the RCX network. This includes checks of the correct aspect structure, data types, uniqueness of IDs and attribute names, presence of NA values, and references between the aspects.

#### Value

logical; whether the object passed all tests.

#### Methods (by class)

- `default`: Default
- `NodesAspect`: Nodes
- `EdgesAspect`: Edges
- `NodeAttributesAspect`: Node attributes
- `EdgeAttributesAspect`: Edge attributes
- `NetworkAttributesAspect`: Network attributes
- `CartesianLayoutAspect`: Cartesian layout
- `CyGroupsAspect`: Cytoscape Groups
- `CyVisualPropertiesAspect`: Cytoscape Visual Properties
- `CyVisualProperty`: Cytoscape Visual Properties
- `CyVisualPropertyProperties`: Cytoscape visual property: Properties

- `CyVisualPropertyDependencies`: Cytoscape visual property: Dependencies
- `CyVisualPropertyMappings`: Cytoscape visual property: Mappings
- `CyHiddenAttributesAspect`: Cytoscape hidden attributes
- `CyNetworkRelationsAspect`: Cytoscape network relations
- `CySubNetworksAspect`: Cytoscape sub-networks
- `CyTableColumnAspect`: Cytoscape table column aspect
- `RCX`: The whole RCX object with all its aspects

#### Examples

```
## Read from a CX file
## reading the provided example network of the package
cxFile <- system.file(
 "extdata",
 "Imatinib-Inhibition-of-BCR-ABL-66a902f5-2022-11e9-bb6a-0ac135e8bacf.cx",
 package = "RCX"
)

rcx = readCX(cxFile)

## validate the network
validate(rcx)

## validate a single aspect
validate(rcx$nodes)
```

---

`visualize` *Visualize a Network*

---

#### Description

Visualize RCX and CX networks in RStudio or in an external browser.

#### Usage

```
visualize(x, layout = NULL, openExternal = FALSE)

## S3 method for class 'RCX'
visualize(x, layout = NULL, openExternal = FALSE)

## S3 method for class 'CX'
visualize(x, layout = NULL, openExternal = FALSE)
```

#### Arguments

| | |
|---|---|
| `x` | network; RCX or CX object |
| `layout` | named character or list; e.g. `c(name="random")` |
| `openExternal` | logical; whether to open in an external browser instead of the RStudio viewer |

#### Details

This function uses the Java Script library used by the NDEx platform (https://ndexbio.org/) to visualize the RCX or CX network from toCX. In the first case, the RCX is converted to CX (JSON) using toCX.

By default the visualization is opened in RStudio in the *Viewer* panel. If this function is not executed in RStudio, the visualization is opened in the standard web-browser. This also can be forced from within RStudio using *openExternal*.

If the network contains the necessary Cytoscape styles (see http://manual.cytoscape.org/en/stable/Styles.html) the network is visualized as seen on the NDEx platform.

To define the layout of the network the coordinate from CartesianLayout are used to determine the location of the nodes. If this aspect is missing, or the the coordinates should be ignored, the *layout* parameter can be used to set a different layout.

*layout* follows therefore the definition of Cytoscape.js (see https://js.cytoscape.org/#layouts). A simple definition can be setting only the *name* of the desired layout, e.g. `random`. Additional options can be passed as named list, where the values are passed without quoting. This allows for even passing Java Script functions to Cytoscape.js.

The visualization can also be saved as HTML file using the writeHTML function instead of this one.

#### Value

NULL

#### See Also

rcxToJson, readCX, writeCX

#### Examples

```
## prepare RCX
rcx = createRCX(
  createNodes(name = c("a","b","c")),
  createEdges(
    source=c(0,0,1),
    target=c(1,2,2)
  )
)

## visualize the network
 visualize(rcx)

## force a different layout
 visualize(rcx, c(name="cose"))

## force a different layout with Java Script parameters
 visualize(rcx, layout = c(name="random",animate="true"))

## even pass a Java Script function
```

```
 visualize(
   rcx,
   layout = c(
     name="random",
     animate="true",
     animateFilter="function ( node, i ){ return true; }"
   )
 )

## open the visualization in an external browser
 visualize(
   rcx,
   layout = c(name="cose"),
   openExternal = TRUE
 )
```

---

writeCX *Write RCX to file*

---

#### Description

These function write an RCX object or a CX object to a file.

#### Usage

```
writeCX(x, file, verbose = FALSE, pretty = FALSE)

## S3 method for class 'RCX'
writeCX(x, file, verbose = FALSE, pretty = FALSE)

## S3 method for class 'CX'
writeCX(x, file, verbose = FALSE, pretty = FALSE)
```

#### Arguments

| | |
|---|---|
| x | RCX or CX object |
| file | character; the name of the file to which the data are written |
| verbose | logical; whether to print what is happening |
| pretty | logical; adds indentation whitespace to JSON output. Can be TRUE/FALSE or a number specifying the number of spaces to indent. See `jsonlite::prettify()` |

#### Value

file character; the name of the file to which the data were written

#### See Also

toCX, rcxToJson, readCX

#### Examples

```
NULL
```

---

`writeHTML` *Save network visualization as HTML file*

---

#### Description

Save an interactive single page visualization of RCX and CX networks as an HTML file containing all necessary Java Script.

#### Usage

```
writeHTML(x, file, layout = NULL, verbose = FALSE)

## S3 method for class 'RCX'
writeHTML(x, file, layout = NULL, verbose = FALSE)

## S3 method for class 'CX'
writeHTML(x, file, layout = NULL, verbose = FALSE)
```

#### Arguments

| | |
|---|---|
| `x` | network; RCX or CX object |
| `file` | character; path, where the html file should be saved |
| `layout` | named character or list; e.g. `c(name="random")` |
| `verbose` | logical; whether to print what is happening |

#### Details

This function uses the Java Script library used by the NDEx platform (`https://ndexbio.org/`) to visualize the RCX or CX network. The RCX is therefore converted to CX (JSON) using toCX.

If the network contains the necessary Cytoscape styles (see `http://manual.cytoscape.org/en/stable/Styles.html`) the network is visualized as seen on the NDEx platform.

To define the layout of the network the coordinate from CartesianLayout are used to determine the location of the nodes. If this aspect is missing, or the the coordinates should be ignored, the *layout* parameter can be used to set a different layout.

*layout* follows therefore the definition of Cytoscape.js (see `https://js.cytoscape.org/#layouts`). A simple definition can be setting only the *name* of the desired layout, e.g. `random`. Additional options can be passed as named list, where the values are passed without quoting. This allows for even passing Java Script functions to Cytoscape.js.

To visualize the network in RStudio the visualize function can be used instead.

#### Value

file character; path, where the html file has been saved

#### See Also

rcxToJson, readCX, writeCX

#### Examples

```
## prepare RCX
rcx = createRCX(
  createNodes(name = c("a","b","c")),
  createEdges(
    source=c(0,0,1),
    target=c(1,2,2)
  )
)

cx = toCX(rcx)

htmlFile = tempfile(fileext = ".html")

## save the html
writeHTML(rcx, htmlFile)

## or
writeHTML(cx, htmlFile)

## force a different layout
writeHTML(rcx, htmlFile, c(name="cose"))

## force a different layout with Java Script parameters
writeHTML(rcx, htmlFile, layout = c(name="random",animate="true"))

## even pass a Java Script function
writeHTML(
   rcx,
   htmlFile,
   layout = c(
     name="random",
     animate="true",
     animateFilter="function ( node, i ){ return true; }"
   )
)
```

### Index

∗ **datasets**
aspectClasses, 3

aspectClass2Name (Convert-Names-and-Classes), 5
aspectClasses, 3, 44, 58, 60
aspectName2Class (Convert-Names-and-Classes), 5

base::print(), 9

cartesian layout, 35, 39
CartesianLayout, 4, 52, 66, 67, 106, 108
Convert-Names-and-Classes, 5
countElements, 6, 43
createCartesianLayout (CartesianLayout), 4
createCyGroups (CyGroups), 9
createCyHiddenAttributes (CyHiddenAttributes), 11
createCyNetworkRelations (CyNetworkRelations), 13
createCySubNetworks (CySubNetworks), 14
createCyTableColumn (CyTableColumn), 16
createCyVisualProperties (CyVisualProperties), 17
createCyVisualProperty (CyVisualProperty), 19
createCyVisualPropertyDependencies (CyVisualPropertyDependencies), 22
createCyVisualPropertyMappings (CyVisualPropertyMappings), 24
createCyVisualPropertyProperties (CyVisualPropertyProperties), 26
createEdgeAttributes (EdgeAttributes), 28
createEdges (Edges), 30
createNetworkAttributes (NetworkAttributes), 45
createNodeAttributes (NodeAttributes), 47
createNodes (Nodes), 50
createRCX (RCX-object), 52
custom-print, 8
CX, 107, 108
CyGroups, 9, 52, 68, 69
CyHiddenAttributes, 11, 52, 71, 72
CyNetworkRelations, 4, 13, 15, 17, 52, 74, 75, 78
CySubNetworks, 4, 14, 53, 75, 77, 78, 81, 84, 88
CyTableColumn, 16, 53, 80, 81
Cytoscape visual properties, 22, 24, 26, 82
CyVisualProperties, 7, 17, 18–20, 22, 24, 26, 32, 52, 82–84, 86
CyVisualProperty, 17, 18, 19, 20–24, 26, 27, 32, 33, 57, 82, 84, 86, 88
CyVisualPropertyDependencies, 18, 20, 22, 23, 24, 26, 27, 32, 84, 86, 88
CyVisualPropertyMappings, 18, 20, 22, 23, 24, 25–27, 32, 84, 86, 88
CyVisualPropertyProperties, 18, 20, 22–24, 26, 27, 32, 84, 86, 88

edge, 34, 35, 39
edge id, 15
edge ids, 10, 28
EdgeAttributes, 28, 46, 49, 52, 90, 91, 97, 100
edgeAttributes, 35, 39
Edges, 10, 29, 30, 52, 69, 78, 91, 93, 94
edges, 35, 39

fromGraphNEL (graphNEL), 34
fromIgraph (Igraph), 38

getCyVisualProperty, 19, 32, 84, 88

graph vertex attributes, *35*
graphNEL, *34*, 34, *35*, *40*

hasIds, 36, *43*
hasIds(), *7*, *38*, *43*, *60*, *62*

idProperty, 37
idProperty(), *7*, *37*, *43*, *60*, *62*
Igraph, *35*, 38
igraph, *38–40*
igraph graph attributes, *40*
igraph vertex attributes, *39*
igraph::as_graphnel(), *35*

jsonlite, *42*, *58*
jsonlite::prettify(), *65*, *107*
jsonToRCX, 40, *57*, *59*

maxId, 42, *43*, *69*, *78*, *93*, *102*
maxId(), *7*, *37, 38*, *43*, *60*, *62*
Meta-data, 43

NetworkAttributes, 45, *49*, *52*, *91*, *95–97*, *100*
node, *4*, *34*, *39*
node id, *15*, *31*
node ids, *4*, *10*, *48*
NodeAttributes, *42*, *46*, 47, *52*, *91*, *97–100*
nodeAttributes, *35*, *39*
Nodes, *7*, *10*, *31*, *35*, *39*, *48*, 50, *52*, *69*, *78*, *99*, *101, 102*
nodes, *35*, *39*, *52*

parseJSON *(*readCX*)*, 58
print.CartesianLayoutAspect *(*custom-print*)*, 8
print.CyGroupsAspect *(*custom-print*)*, 8
print.CyHiddenAttributesAspect *(*custom-print*)*, 8
print.CyNetworkRelationsAspect *(*custom-print*)*, 8
print.CySubNetworksAspect *(*custom-print*)*, 8
print.CyTableColumnAspect *(*custom-print*)*, 8
print.CyVisualPropertiesAspect *(*custom-print*)*, 8
print.CyVisualProperty *(*custom-print*)*, 8
print.CyVisualPropertyDependencies *(*custom-print*)*, 8
print.CyVisualPropertyMappings *(*custom-print*)*, 8
print.CyVisualPropertyProperties *(*custom-print*)*, 8
print.EdgeAttributesAspect *(*custom-print*)*, 8
print.EdgesAspect *(*custom-print*)*, 8
print.MetaDataAspect *(*custom-print*)*, 8
print.NetworkAttributesAspect *(*custom-print*)*, 8
print.NodeAttributesAspect *(*custom-print*)*, 8
print.NodesAspect *(*custom-print*)*, 8
print.RCX *(*custom-print*)*, 8
processCX, *42*
processCX *(*readCX*)*, 58

RCX, *7–9*, *12*, *16*, *29*, *34, 35*, *38–40*, *42–46*, *48*, 51, *53*, *57–59*, *62*, *64–69*, *71, 72*, *74, 75*, *77, 78*, *80–84*, *90, 91*, *93–100*, *102*, *104–108*
RCX-object, 52
rcxToJson, *42*, 55, *65*, *106, 107*, *109*
readCX, *42*, *57*, 58, *65*, *106, 107*, *109*
readJSON *(*readCX*)*, 58
referredBy, 60
referredBy(), *7*, *37, 38*, *43*, *62*
refersTo, 61
refersTo(), *7*, *37, 38*, *43*, *60*

subAspectClasses *(*aspectClasses*)*, 3
subnetwork, *20*, *35*, *40*
subnetwork id, *4*, *13*, *16*, *28*, *35*, *40*, *46*, *48*
subnetworks, *12*, *14*, *16*, *29*, *46*, *48*
summary, *9*, 62

toCX, *42*, *57*, *65*, 65, *106–108*
toGraphNEL *(*graphNEL*)*, 34
toIgraph *(*Igraph*)*, 38

updateAspectClasses *(*aspectClasses*)*, 3
updateCartesianLayout, *5*, 66
updateCyGroups, *10*, 68
updateCyHiddenAttributes, *12*, 71
updateCyNetworkRelations, *14*, 74
updateCySubNetworks, *15*, 77
updateCyTableColumn, *17*, 79
updateCyVisualProperties, *19*, *21*, *23*, *25*, *28*, *33*, 82, *88*

updateCyVisualProperty, *19*, *21*, *23*, *25*, *28*, *33*, *84*, 86
updateEdgeAttributes, *29*, 90
updateEdges, *31*, *59*, 93
updateMetaData *(*Meta-data*)*, 43
updateMetaDataProperties, *44, 45*, 94
updateNetworkAttributes, *46*, 95
updateNodeAttributes, *49*, 98
updateNodes, *51*, *59*, 101

validate, 103
vertex, *35*, *39*
visualize, 105, *108*

writeCX, *42*, *57*, *59*, *65*, *106*, 107, *109*
writeHTML, *106*, 108
