## Supplementary material for "RCX – an R package adapting the Cytoscape Exchange format for biological networks": RCX Cheat Sheet

```
createRCX(
  nodes,
    createNodes(id=NULL, name=NULL, represents=NULL)
  edges,
    createEdges(id=NULL, source, target, interaction=NULL)
  nodeAttributes,
    createNodeAttributes(propertyOf, name, value, dataType=NULL, isList=NULL, subnetworkId=NULL)
  edgeAttributes,
    createEdgeAttributes(propertyOf, name, value, dataType=NULL, isList=NULL, subnetworkId=NULL)
  networkAttributes,
    createNetworkAttributes(name, value, dataType=NULL, isList=NULL, subnetworkId=NULL)
  cartesianLayout,
    createCartesianLayout(node, x, y, z=NULL, view=NULL)
  cySubNetworks,
    createCySubNetworks(id, nodes=NULL, edges=NULL)
  cyGroups,
    createCyGroups(id=NULL, name, nodes=NULL, externalEdges=NULL, internalEdges=NULL, collapsed=NULL)
  cyHiddenAttributes,
    createCyHiddenAttributes(name, value, dataType=NULL, isList=NULL, subnetworkId=NULL)
  cyNetworkRelations,
    createCyNetworkRelations(child, parent=NULL, name=NULL, isView=FALSE)
  cyTableColumn,
    createCyTableColumn(appliesTo, name, dataType=NULL, isList=NULL, subnetworkId=NULL)
  cyVisualProperties,
    createCyVisualProperties(network=NULL, nodes=NULL, edges=NULL, defaultNodes=NULL, defaultEdges=NULL)

      createCyVisualProperty(properties=NULL, dependencies=NULL, mappings=NULL, appliesTo=NULL, view=NULL)

  checkReferences=TRUE
)
```

```
readCX(file, verbose=FALSE, aspectClasses=NULL)
```

```
updateMetaData(
  x,
  version = NULL,
  consistencyGroup = NULL,
  properties=NULL,
  aspectClasses=NULL
)
```

```
createCyVisualPropertyMappings(
  type,
  definition,
  name=NULL
)
```

```
createCyVisualPropertyDependencies(
  value,
  name=NULL
)
```

```
createCyVisualPropertyProperties(
  value,
  name=NULL
)
```

```
validate(x, verbose=TRUE)
```

```
visualize(
  x,
  layout=NULL,
  openExternal=FALSE
)
```

```
writeCX(
  x,
  file,
  verbose=FALSE,
  pretty=FALSE
)
```

```
toGraphNEL(rcx, directed=FALSE)
```
→ graphNEL

```
toIgraph(rcx, directed=FALSE)
```
→ igraph

### DATA MODEL STRUCTURE

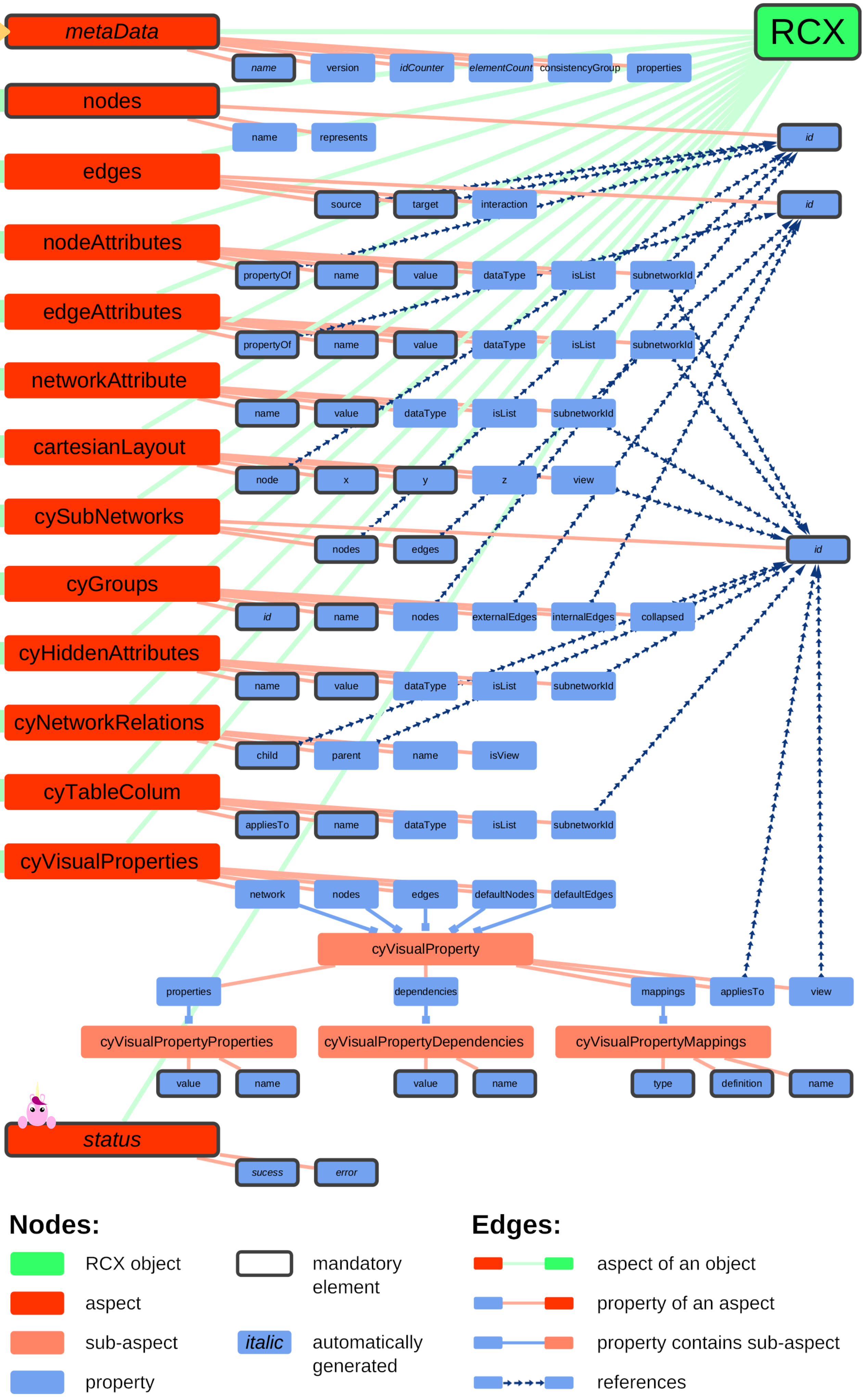
